# PEX39 facilitates the peroxisomal import of PTS2 proteins

**DOI:** 10.1101/2024.04.30.591961

**Authors:** Walter W. Chen, Tony A. Rodrigues, Daniel Wendscheck, Ana G. Pedrosa, Chendong Yang, Tânia Francisco, Till Möcklinghoff, Alexandros Zografakis, Bernardo Nunes-Silva, Reut Ester Avraham, Ana R. Silva, Maria J. Ferreira, Hirak Das, Julian Bender, Silke Oeljeklaus, Varun Sondhi, Maya Schuldiner, Einat Zalckvar, Kay Hofmann, Hans R. Waterham, Ralph J. DeBerardinis, Jorge E. Azevedo, Bettina Warscheid

**Affiliations:** Division of Neonatal-Perinatal Medicine, Department of Pediatrics, University of Texas Southwestern Medical Center, Dallas, TX, USA; Children’s Medical Center Research Institute, University of Texas Southwestern Medical Center, Dallas, TX, USA; Instituto de Investigação e Inovação em Saúde (i3S), Universidade do Porto, Porto, Portugal; Instituto de Biologia Molecular e Celular (IBMC), Universidade do Porto, Porto, Portugal; Instituto de Ciências Biomédicas Abel Salazar (ICBAS), Universidade do Porto, Porto, Portugal; Biochemistry II, Theodor Boveri-Institute, Biocenter and Faculty of Chemistry and Pharmacy, University of Würzburg, 97074 Würzburg, Germany; Biochemistry and Functional Proteomics, Institute of Biology II, Faculty of Biology, University of Freiburg, 79104 Freiburg, Germany; Department of Molecular Genetics, Weizmann Institute of Science, Rehovot 7610001, Israel; Division of Pulmonary and Critical Care Medicine, Department of Internal Medicine, UT Southwestern Medical Center, Dallas, TX, USA; The Mina and Everard Goodman Faculty of Life Sciences, Bar-Ilan University, Ramat-Gan 52900, Israel; Institute for Genetics, University of Cologne, 50674 Cologne, Germany; Laboratory Genetic Metabolic Diseases, Amsterdam UMC location University of Amsterdam, Amsterdam, the Netherlands; Amsterdam Gastroenterology, Endocrinology and Metabolism, Amsterdam, the Netherlands; Simmons Comprehensive Cancer Center, UT Southwestern Medical Center, Dallas, TX, USA; Howard Hughes Medical Institute, University of Texas Southwestern Medical Center, Dallas, TX, USA; German Cancer Research Center, Systems Biology of Signal Transduction, 69120 Heidelberg, Germany

**Keywords:** Peroxisome, biogenesis, peroxin, protein import, PTS2, PEX7, PEX39, PEX13, [R/K]PWE, protein complexes

## Abstract

Peroxisomes are metabolic organelles essential for human health. Defects in peroxisomal biogenesis proteins (peroxins/PEXs) cause devastating disease. PEX7 binds newly synthesized proteins containing a type 2 peroxisomal targeting signal (PTS2) to enable their import from the cytosol into peroxisomes, although many aspects of this import pathway remain enigmatic. Utilizing *in vitro* assays, yeast, and human cells, we show that PEX39, a previously uncharacterized protein, is a cytosolic peroxin that facilitates PTS2-protein import by binding PEX7 and stabilizing its interaction with PTS2 cargo. PEX39 and PEX13, a peroxisomal membrane translocon protein, both possess a KPWE motif necessary for PEX7 binding. Sequential binding of PEX7 to this motif in PEX39 and PEX13 provides a novel paradigm for how PTS2 cargo engage the translocation machinery. Collectively, our work uncovers an ancient and functionally important relationship among PEX39, PEX7, and PEX13, offering insights that will advance our understanding of peroxisomal biogenesis and disease.

## INTRODUCTION

Peroxisomes are organelles that are present in nearly all eukaryotes and provide a cellular niche for important biochemical processes.^1^ Peroxisomal enzymes participate in the oxidation of fatty acids, detoxification of reactive oxygen species, and, in mammals, the synthesis of bile acids and myelin.^2^ Peroxins (PEXs) are proteins essential for peroxisomal biogenesis, with multiple peroxins facilitating in mammalian cells the import of more than 60 different enzymes from the cytosol into the peroxisomal matrix (*i.e.*, lumen).^3^ Defects in peroxins cause devastating human diseases, such as Zellweger Spectrum Disorders.^4^

All peroxisomal matrix proteins are encoded in the nucleus and synthesized in the cytosol, so peroxisomal biogenesis and functions depend on specific protein import mechanisms. The import of peroxisomal enzymes occurs via a remarkable mechanism through which even folded proteins and large protein complexes can reach the organellar matrix.^5–7^ This process differs from protein import into the mitochondria or endoplasmic reticulum, which can only translocate unfolded monomers that are then folded within the organelle.^8,9^ Peroxisomal enzymes harbor a type 1 peroxisomal targeting signal (PTS1) (*i.e.*, C-terminal tripeptide SKL or variants of it) or, less commonly, a type 2 peroxisomal targeting signal (PTS2) [*i.e.*, N-terminal R-(L/V/I/Q)-X-X-(L/V/I/H)-(L/S/G/A)-X-(H/Q)-(L/A)].^10,11^ Although few in number, PTS2 proteins serve crucial roles in cellular metabolism, with defects in the PTS2 proteins phytanoyl-CoA 2-hydroxylase (PHYH) (*e.g.*, fatty acid α-oxidation) and alkyl-DHAP synthase (AGPS) (*e.g.*, myelin synthesis) causing human diseases.^12,13^ In the cytosol, the receptors PEX5 and PEX7 bind cargo through their PTS1 and PTS2, respectively, to initiate the import cycle. Notably, cargo-loaded PEX7 also requires binding of a co-receptor (*e.g.*, Pex18/Pex21 in *Saccharomyces cerevisiae* and the long isoform of PEX5 in humans) to engage the downstream import machinery.^10,11^ Cargo-loaded PEX5 or PEX7-coreceptor complexes are then recruited to the peroxisomal docking/translocation module containing the membrane proteins PEX13 and PEX14,^14–22^ and in yeast PEX17.^23–25^ Notably, PEX13 has been recently proposed to form the conduit through which peroxisomal matrix proteins are translocated.^26,27^ However, our understanding of various aspects of peroxisomal protein import remains incomplete.

Here, we demonstrate that the previously uncharacterized *S. cerevisiae* (yeast) protein Yjr012c and human protein C6ORF226 are orthologs of a peroxin we have named PEX39, which facilitates the import of PTS2 proteins. PEX39 and PEX13 both possess an ancient, highly conserved [R/K]PWE motif that is necessary for the binding of PEX7 and proper functioning of each protein. Collectively, our results show that PEX39 is a previously unknown peroxin and reveal how PEX39 and [R/K]PWE motifs contribute to the biogenesis of peroxisomes.

## RESULTS

PEX39 is a cytosolic protein that interacts with members of the PTS2-protein import pathway To better understand the import of PTS2 proteins, we examined PEX7 in the interactome databases BioGRID^28^ and BioPlex.^29,30^ Our analysis revealed that two proteins of unknown function, *S. cerevisiae* Yjr012c and human C6ORF226, interact with PEX7 orthologs despite lacking a PTS2. The C6ORF226 interactome data also contain peroxisomal phytanoyl-CoA 2-hydroxylase (PHYH) and peroxisomal 3-ketoacyl-CoA thiolase (ACAA1), two PTS2 proteins involved in fatty acid oxidation (Figure 1A).^2^

**Figure 1.**
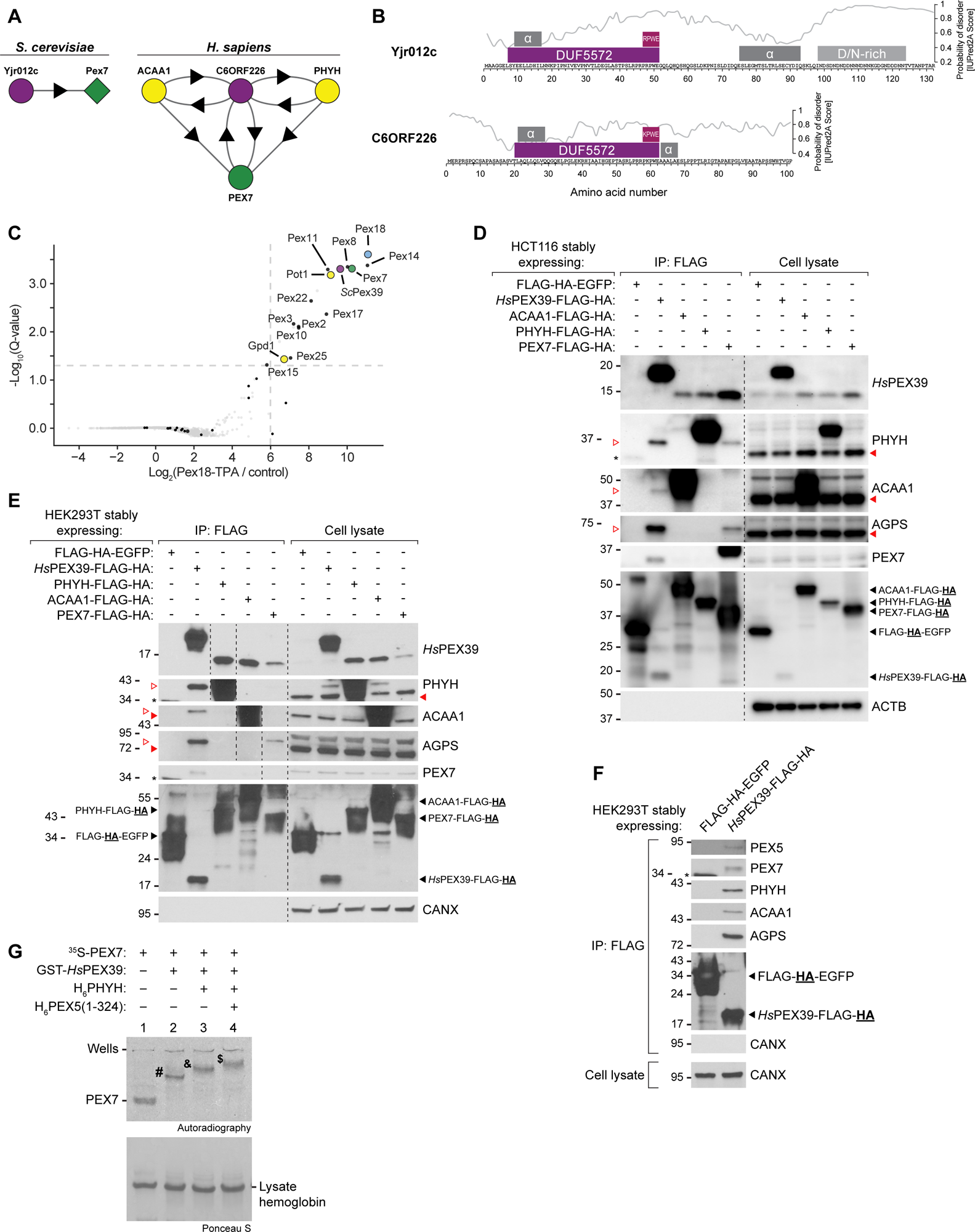
PEX39 interacts with members of the PTS2-protein import pathway. (A) Identification of Yjr012c and C6ORF226 as uncharacterized proteins that interact with the PTS2-protein import system via BioGRID^28^ and BioPlex,^29,30^ respectively. A circle represents the purified protein (bait) and a diamond represents the accompanying interactor (prey). Arrows point from bait to prey. Two circles with bidirectional arrows between them indicate that both proteins have served as baits and also been identified as accompanying interactors. For *S. cerevisiae*, only protein-protein interactions found in at least two independent studies per BioGRID were considered. The combined BioPlex interactome from HCT116 and HEK293T cells is shown for human/*Homo sapiens* (*H. sapiens*). (B) Domain analysis and alignment of Yjr012c and C6ORF226. Probabilities of disorder (determined using IUPred2A^61^), the DUF5572, [R/K]PWE motifs, and further regions of interest are indicated. α, α-helical. (C) *Sc*Pex39 is a specific component of Pex18 complexes as determined by label-free quantitative affinity purification-mass spectrometry. Pex18 complexes were affinity purified from soluble fractions of oleic acid-grown wild-type and Pex18-TPA-expressing cells (n = 3). Enrichment of proteins in Pex18 complexes and Q-values were determined using the rank sum method.^62^ Known peroxins and PTS2 proteins are labelled and/or marked by black dots. Dashed lines indicate a Q-value threshold of 0.05 and a fold-enrichment of 64. (D) *Hs*PEX39 interacts with PEX7, PHYH, ACAA1, and AGPS per assessment with human HCT116 cells. Anti-FLAG immunoprecipitates and cell lysates were prepared from HCT116 cells stably expressing the indicated proteins. Samples were analyzed by immunoblotting for the indicated proteins; detection of HA denoted by “<u>HA</u>” in the labelled black arrowheads identifying the corresponding proteins. Numbers along the left side indicate molecular weights (kD). Dashed lines indicate where different lanes of the same membrane were brought together. For the PHYH, ACAA1, and AGPS blots, the solid and open red arrowheads indicate the mature and precursor forms of these proteins, respectively. An asterisk indicates a non-specific band. (E) *Hs*PEX39 interacts with PEX7, PHYH, ACAA1, and AGPS per assessment with human HEK293T cells. Anti-FLAG immunoprecipitates and cell lysates were prepared from HEK293T cells stably expressing the indicated proteins. Samples were analyzed by immunoblotting for the indicated proteins. Occasionally, it was necessary to have blanks sandwiching a given sample to prevent spillover of immunoblot signal to other samples. Annotation of the immunoblots is otherwise the same as described for (D). (F) *Hs*PEX39 interacts with PEX5 per assessment with HEK293T cells. Anti-FLAG immunoprecipitates and cell lysates were prepared from HEK293T cells stably expressing the indicated proteins. Samples were analyzed by immunoblotting for the indicated proteins; detection of HA denoted by “<u>HA</u>” in the labelled black arrowheads identifying the corresponding proteins. An asterisk indicates a non-specific band. (G) *Hs*PEX39 can complex with PEX7, PHYH, and PEX5 *in vitro*. Radiolabeled H6PEX7 was pre-incubated or not with the recombinant proteins GST-*Hs*PEX39, H6PHYH, and H6PEX5(1-324), as indicated. Samples were analyzed by native-PAGE and autoradiography. In-gel migration of PEX7 alone (PEX7), lysate hemoglobin, and complexes PEX7-*Hs*PEX39 (#), PEX7-PHYH-*Hs*PEX39 (&), PEX7-PEX5-PHYH-*Hs*PEX39 ($) are indicated. The autoradiograph and the corresponding Ponceau S-stained membrane are shown. See also Figure S1.

A comparison of the C6ORF226 and Yjr012c sequences revealed notable similarities, such as the presence of a Domain of Unknown Function (DUF5572) and an [R/K]PWE motif at the DUF5572 C-terminus, indicating that these two proteins are orthologs (Figure 1B). A broader examination of numerous organisms revealed that the most conserved feature of Yjr012c/C6ORF226 orthologs is the [R/K]PWE motif, with a predominance of K over R at the first position (Figure S1A, left). Despite the absence of a DUF5572 domain, PEX13 also possesses a conserved KPWE motif at its N-terminus, which had been observed previously^31^ and recently predicted to interact directly with yeast Pex7.^32^ Of note, Yjr012c/C6ORF226 orthologs occur in all eukaryotic kingdoms, suggesting a similar evolutionary age to known peroxins (Figure S1A, right). Interestingly, no ortholog is present in *Caenorhabditis elegans* and *Drosophila melanogaster*, which lack the PTS2 pathway.^33,34^ Remarkably, outside of opisthokonts (*e.g.,* fungi and animals), Yjr012c/C6ORF226-related sequences are fused to PEX14. Collectively, these data suggested to us that Yjr012c/C6ORF226 are not merely cargo proteins but peroxins involved in protein import. We hereafter refer to human/*Homo sapiens* C6ORF226 as *Hs*PEX39 and yeast/*S. cerevisiae* Yjr012c as *Sc*Pex39.

We next validated and explored the interactions reported for *Sc*Pex39 and *Hs*PEX39 (Figure 1A). In *S. cerevisiae*, quantitative affinity purification-mass spectrometry experiments using Pex18 C-terminally fused with a TPA tag (Pex18-TPA) as bait revealed *Sc*Pex39 as a specific interactor, along with the known binding partners Pex7, other peroxins, and peroxisomal 3-ketoacyl-CoA thiolase (Pot1), which is a PTS2 protein (Figures 1C and S1B, Table S2). The PTS2 protein glycerol-3-phosphate dehydrogenase (Gpd1) was only slightly enriched with Pex18-TPA, confirming the specificity of this interactome data given that Gpd1 requires Pex7 and Pex21 for peroxisomal import.^35,36^ To examine *Hs*PEX39, we expressed and immunoprecipitated FLAG-HA-tagged versions of EGFP (negative control), *Hs*PEX39, PHYH, ACAA1, and PEX7 using the human cell lines HCT116 (Figure 1D) and HEK293T (Figure 1E). The control proteins β-actin (ACTB) or calnexin (CANX) were not detected in immunoprecipitates. In both cell lines, FLAG-HA-tagged PHYH, ACAA1, and PEX7 co-immunoprecipitated endogenous *Hs*PEX39 while the EGFP control did not. *Hs*PEX39-FLAG-HA also co-immunoprecipitated endogenous PEX7 and the three PTS2 proteins PHYH, ACAA1, and alkyl-DHAP synthase (AGPS). In addition, PHYH, ACAA1, and AGPS co-immunoprecipitating with *Hs*PEX39-FLAG-HA or PEX7-FLAG-HA migrated at the expected molecular weights of their precursor forms, consistent with PEX7 only being able to bind precursor PTS2 proteins due to removal of the PTS2 when the protein is converted to the mature form in the peroxisomal lumen; of note, this maturation process for PTS2 proteins does not occur in yeast.^11^ In agreement with the interaction between Pex18 and *Sc*Pex39 in yeast (Figure 1C), *Hs*PEX39-FLAG-HA also co-immunoprecipitated the co-receptor PEX5 from human cell lysates (Figure 1F). Consistent with PEX39 interacting with PEX7, PEX5/Pex18, and PTS2 cargo, fluorescence microscopy revealed that a *Sc*Pex39-mNeonGreen fusion protein localized to both the cytosol and peroxisomes (Figure S1C), while cellular fractionation revealed that endogenous *Hs*PEX39 is entirely cytosolic (Figure S1D).

To further explore the observed interactions, we produced recombinant *Hs*PEX39 as a GST-tagged protein and analyzed radiolabeled ^35^S-PEX7-containing complexes by native-PAGE.^37^ Incubation of ^35^S-PEX7 with GST-*Hs*PEX39 led to a shift in the migration of the radiolabeled protein, indicating that *Hs*PEX39 can interact with PEX7 to form a stable PEX7-*Hs*PEX39 complex (“#” in Figure 1G, lane 1 vs 2). Shifts in migration also reveal that trimeric PEX7-PHYH-*Hs*PEX39 (“&” in Figure 1G, lane 3) and tetrameric PEX7-PEX5-PHYH-*Hs*PEX39 (“$” in Figure 1G, lane 4) complexes are formed. Collectively, these data demonstrate that PEX39 is a cytosolic protein that interacts with the different members of the PTS2-protein import pathway.

### Loss of PEX39 impairs import of PTS2 proteins

For a protein to be considered a peroxin, it must participate in a process needed for peroxisomal biogenesis, such as the import of peroxisomal proteins.^38,39^ To explore the functional importance of PEX39 in peroxisomal biogenesis, we generated loss-of-function models in both yeast and human cells. We first assessed the growth of *Scpex39*-null (*Scpex39*Δ yeast in liquid media with glucose or oleic acid as the primary energy source; in the latter condition, peroxisomal biogenesis is stimulated and cellular proliferation requires β-oxidation of fatty acids, which occurs exclusively in peroxisomes in *S. cerevisiae*.^40^ Because the PTS2 protein Pot1 is a thiolase necessary for β-oxidation, defects in PTS2-protein import should impair fitness selectively in medium with oleic acid but not glucose.^41^ Consistent with this, while *Scpex39*Δ cells had similar fitness to wild-type cells when grown on glucose (Figure S2A), these cells grew considerably worse on oleic acid (Figure 2A). As expected, *pex7*Δ and *pot1*Δ yeast also grew poorly on oleic acid. Importantly, the fitness defect observed for *Scpex39*Δ cells grown on oleic acid could be rescued via introduction of a plasmid with the *Scpex39* gene under the control of its native promoter, thus demonstrating that the observed phenotype is not an off-target effect.

**Figure 2.**
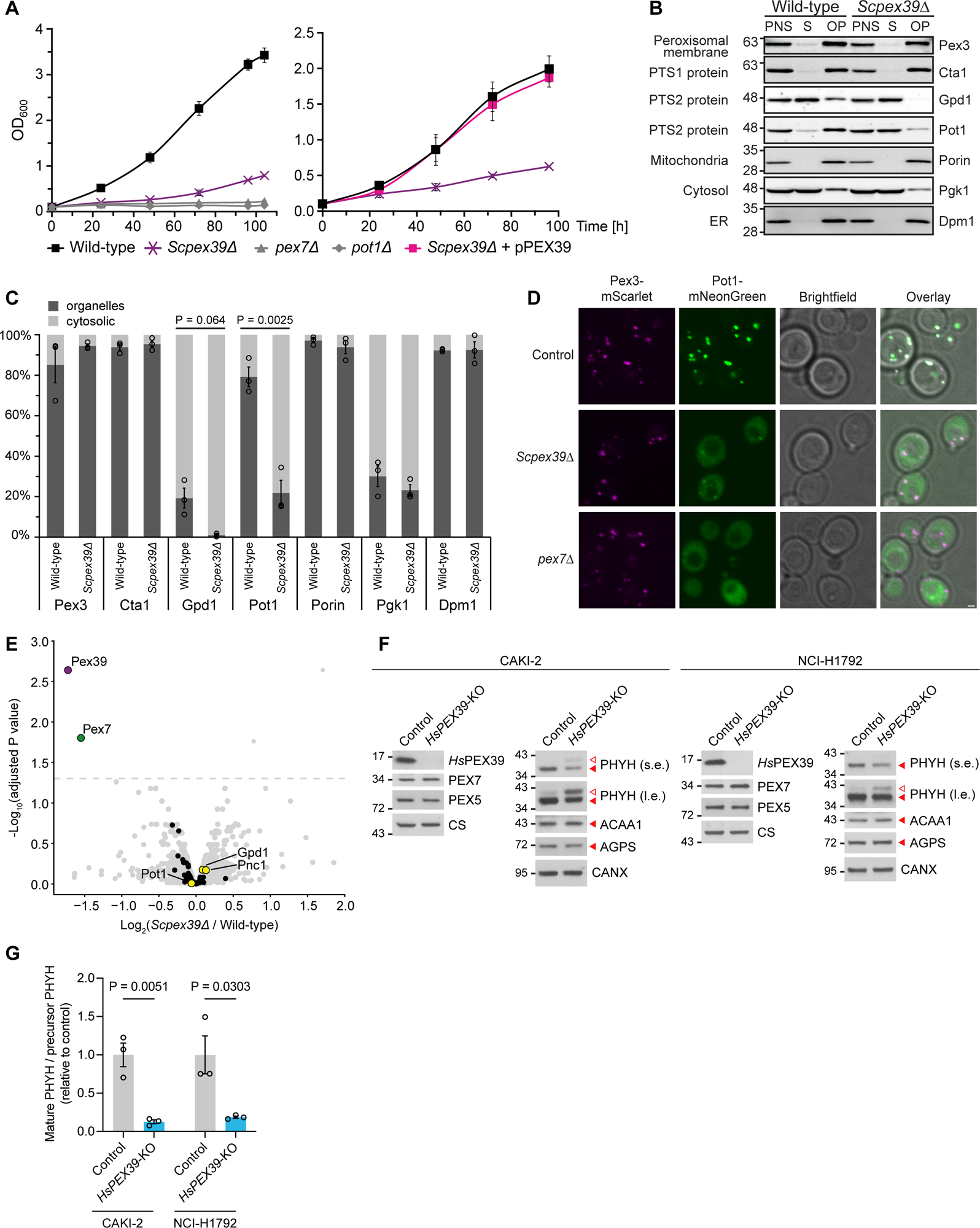
Loss of PEX39 impairs import of PTS2 proteins. (A) Loss of *Sc*Pex39 confers a fitness defect in yeast grown in oleic acid. Cells were precultured in 0.3% glucose medium for 16 h at 30°C and then shifted to oleic-acid medium for further growth. For complementation, *Scpex39Δ* cells were transformed with a plasmid containing *Scpex39* under control of its endogenous promoter (pPEX39). Cell growth was monitored by measuring the OD600 at the indicated time points. Data are mean ± standard deviation (SD) (n = 4). Error bars may not be visible if the SD is very small. (B) Loss of *Sc*Pex39 specifically impairs PTS2-protein import in yeast per cellular fractionation. A post-nuclear supernatant (PNS) was prepared from oleic acid-grown wild-type and *Scpex39Δ* cells and was further separated into a cytosolic fraction (supernatant, S) and an organellar pellet (OP). Equal volumes of the fractions were analyzed by immunoblotting for the indicated proteins. The experiment was performed in three independent replicates (see also Figure S2B). (C) Quantification of changes in the subcellular distribution of PTS2 proteins upon loss of *Sc*Pex39. Signal intensities of immunoblots shown in (B) and Figure S2B were quantified using ImageJ. For each protein, intensities for the cytosolic supernatant (S) and the organellar pellet (OP) were normalized to the PNS and the sum was set to 100%. Data are mean ± standard error of the mean (SEM) (n = 3), and P values were calculated using unpaired, two-tailed t-tests. (D) Loss of *Sc*Pex39 impairs import of Pot1 in yeast per fluorescent microscopy. Localization of Pot1-mNeonGreen (C-terminal tagging) was examined in control, *Scpex39Δ*, and *pex7Δ* cells after 8 h of growth on oleic acid. Peroxisomes were visualized with Pex3-mScarlet. Scale bar: 1 μm. (E) Loss of *Sc*Pex39 results in decreased levels of Pex7. Whole cell lysates of wild-type (WT) and *Scpex39Δ* cells were analyzed by SILAC-based quantitative mass spectrometry (n = 4 biological replicates). Shown are proteins quantified in at least three biological replicates, except for *Sc*Pex39, which was quantified in only one replicate. PTS2 proteins are highlighted in yellow; further peroxisomal proteins are marked by black dots. Adjusted P values were determined using the “linear models for microarray data” (limma) approach. Dashed horizontal line indicates an adjusted P value threshold of 0.05. (F) Precursor and mature PHYH increase and decrease, respectively, in human *HsPEX39*-knockout cells. CRISPR-Cas9 was used to generate knockouts (*HsPEX39*-KO) and matched controls (Control) in the CAKI-2 and NCI-H1792 cell lines (see STAR Methods). Cellular lysates were analyzed by immunoblotting for the indicated proteins. For the PHYH, ACAA1, and AGPS blots, the solid and open red arrowheads indicate the mature and precursor forms of these proteins, respectively. Precursor forms of ACAA1 and AGPS were undetectable in these experiments. CANX and CS are loading controls. Short and long exposures are denoted s.e. and l.e., respectively. (G) Quantification of changes in mature and precursor PHYH in *HsPEX39*-knockout cells. Band intensities of immunoblots prepared per (F) were quantified using ImageJ. Data are mean ± SEM (n = 3), and P values were calculated using unpaired, two-tailed t-tests. See also Figure S2.

Cellular fractionation of wild-type and *Scpex39*Δ yeast revealed a redistribution from the organellar pellet to the cytosol of Pot1 and to a lesser extent Gpd1, which localizes to the cytosol, nucleus and peroxisomes,^35^ while there were no changes noted for a peroxisomal membrane protein (Pex3) or PTS1 protein (Cta1) (Figures 2B, 2C, S2B). Using Pot1 fused to mNeonGreen, we also observed a redistribution of this reporter PTS2 protein from peroxisomes to the cytosol in *Scpex39*Δ yeast, although to a lesser degree than in *pex7*Δ yeast (Figure 2D). Collectively, these findings demonstrate that loss of *Sc*Pex39 specifically impairs the peroxisomal import of PTS2 proteins.

To more comprehensively assess for changes in the yeast proteome upon loss of *Sc*Pex39, we performed quantitative proteomics on wild-type and *Scpex39*Δ yeast grown on oleic acid and found that Pex7 was greatly decreased in *Scpex39*Δ yeast but PTS2 proteins or other peroxins including Pex5 and Pex18 were not (Figure 2E, Table S3). This reduction in Pex7, which was confirmed by immunoblot analysis (Figure S2C, lanes 1-3), likely contributes to the impairment of PTS2-protein import seen with *Sc*Pex39 loss (Figures 2B, 2C, S2B).

To explore the consequences of *Hs*PEX39 loss, we used CRISPR-Cas9 to generate *HsPEX39* knockouts and matched controls in human CAKI-2 and NCI-H1792 cells (see STAR Methods for rationale and methodology). Firstly, we observed that loss of *Hs*PEX39 in either cell line did not affect the levels of the interacting peroxins PEX7 and PEX5 (Figures 2F and S2D). The lack of a reduction in PEX7 levels with *Hs*PEX39 loss stands in contrast to what we observed in *S. cerevisiae*. Importantly though, loss of *Hs*PEX39 in either cell line led to an increase in precursor PHYH and reduction in mature PHYH, consistent with a defect in the peroxisomal import of PHYH (Figures 2F and 2G). However, no changes in ACAA1 or AGPS were appreciated (Figures 2F and S2D), demonstrating that PTS2 proteins can exhibit differential sensitivity to *Hs*PEX39 loss. Because our knockout cells were generated using one sgRNA against *HsPEX39*, we further validated our findings using two additional sgRNAs targeting distinct sites in the *HsPEX39* gene. In both cell lines, lentiviral CRISPR-Cas9 with these three independent sgRNAs partially depleted *Hs*PEX39 and increased precursor PHYH and reduced mature PHYH, while no differences were appreciated in PEX7, PEX5, ACAA1, and AGPS (Figure S2E). Importantly, cellular fractionation confirmed that the whole-cell increase in precursor PHYH and decrease in mature PHYH seen with *Hs*PEX39 loss reflected an accumulation of precursor PHYH in the cytosolic fraction and reduction of mature PHYH in the organellar fraction, thus demonstrating that this pattern of whole-cell changes does indeed reflect impaired PTS2-protein import in our system (Figure S2F). To conclude, our work in yeast and human cells demonstrates that loss of PEX39 impairs the import of PTS2 proteins.

### Overexpression of PEX39 impairs PTS2-protein import

Having assessed the consequences of depleting PEX39, we next investigated the effects of increasing it. Surprisingly, overexpression of *Hs*PEX39 in wild-type HEK293T cells, which also express endogenous *Hs*PEX39, increased the precursor forms and decreased the mature forms of all three PTS2 proteins, consistent with a defect in PTS2-protein import (Figures 3A and 3B), which is particularly noteworthy since loss of PEX39 also impairs PTS2-protein import. Similar to loss of *Hs*PEX39, overexpression did not alter PEX5 or PEX7 levels (Figure S3A).

**Figure 3.**
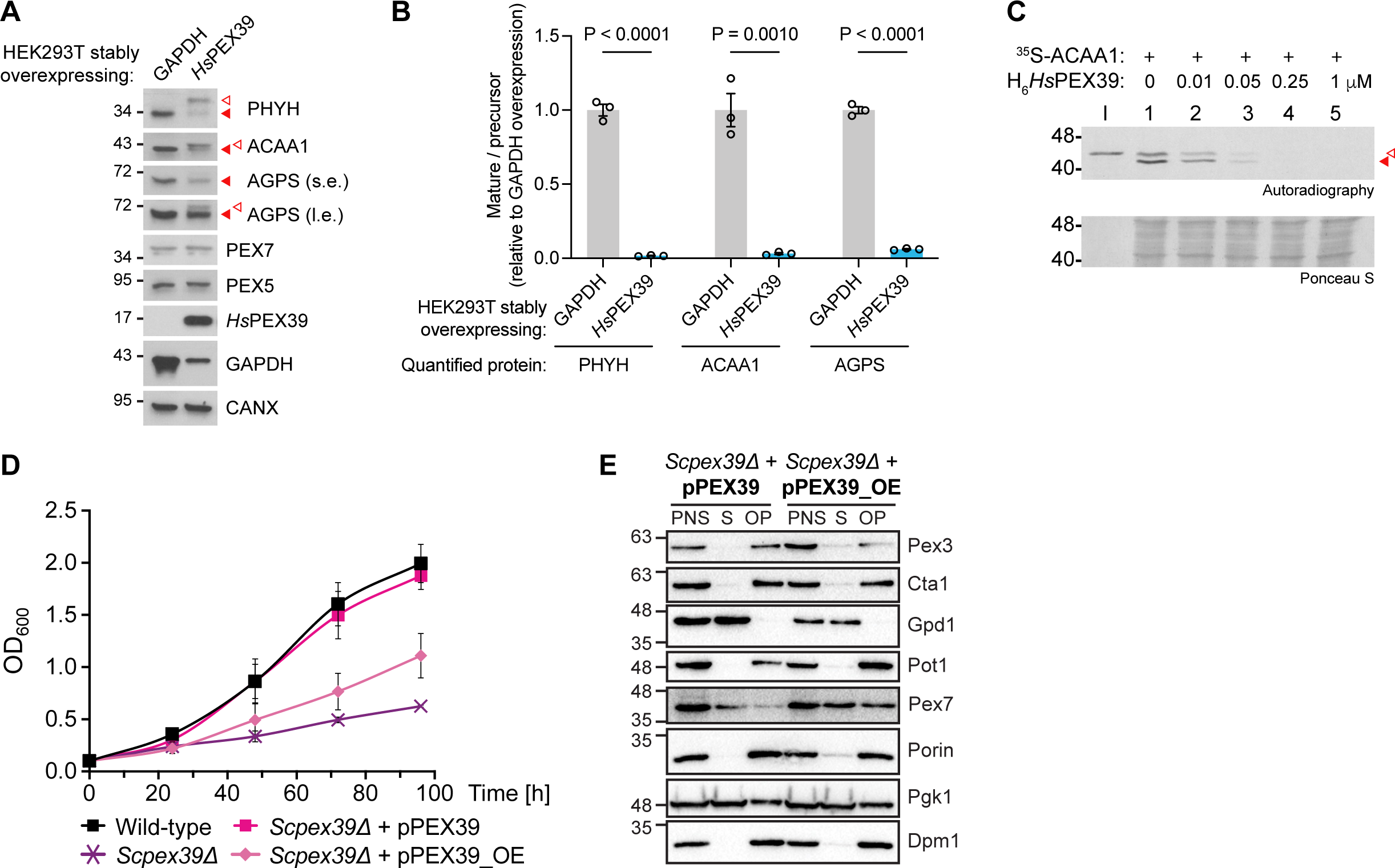
Overexpression of PEX39 impairs PTS2-protein import. (A) Overexpression of *Hs*PEX39 increases precursor forms and decreases mature forms of PHYH, ACAA1, and AGPS in human cells. GAPDH (negative control) or *Hs*PEX39 were stably overexpressed in HEK293T cells and cellular lysates analyzed by immunoblotting for the indicated proteins. For the PHYH, ACAA1, and AGPS blots, the solid and open red arrowheads indicate the mature and precursor forms of these proteins, respectively. CANX is a loading control. (B) Quantification of changes in mature and precursor forms of PHYH, ACAA1, and AGPS upon *Hs*PEX39 overexpression. Band intensities of immunoblots prepared per Figure 3A were quantified using ImageJ. Data are mean ± SEM (n = 3), and P values were calculated using unpaired, two-tailed t-tests. (C) Exogenously added *Hs*PEX39 inhibits peroxisomal import of ACAA1 *in vitro*. ^35^S-ACAA1 *in vitro* import assays at 37°C in the presence of increasing concentrations of H6*Hs*PEX39 as indicated. After incubation, reactions were treated with trypsin and organelles were isolated by centrifugation and analyzed by SDS-PAGE and autoradiography; protection from trypsin and maturation of ACAA1 reflects import into peroxisomes. Precursor and mature forms of ACAA1 denoted by open and solid red arrowheads, respectively. Numbers along the left side of each image indicate molecular weights (kD). See STAR Methods for detailed description of this assay. (D) Overexpression of *Sc*Pex39 confers a fitness defect on yeast grown on oleic acid. Experiment performed as described in Figure 2A. *Scpex39Δ* cells were transformed with a plasmid containing *Scpex39* under control of a TEF2 promoter (pPEX39_OE) for overexpression. Data for wild-type, *Scpex39Δ* and *Scpex39Δ* + pPEX39 are the same as shown in Figure 2A (right plot). Data are mean ± SD (n = 4). Error bars may not be visible if the SD is very small. (E) Cellular fractionation of yeast overexpressing *Sc*Pex39. Experiment performed as described in Figure 2B using *Scpex39Δ* cells transformed with plasmid pPEX39 (endogenous promoter) or pPEX39_OE (TEF2 promoter) for *Sc*Pex39 overexpression. PNS, post-nuclear supernatant; S, cytosolic supernatant; OP, organellar pellet. See also Figure S3.

To investigate this overexpression phenotype further, we examined how exogenously added *Hs*PEX39 affects peroxisomal import using an established *in vitro* import system, in which a radiolabeled reporter protein (*e.g.*, a PTS2 protein or PEX5) is incubated with post-nuclear supernatant containing cytosol and peroxisomes.^42^ Import of radiolabeled PTS2 proteins is assessed by the acquisition of an organelle-associated protease-resistant status and conversion from the precursor to mature form. Given the low amounts of endogenous PTS2 proteins in post-nuclear supernatants, PTS1-protein import can be assessed via the association of radiolabeled PEX5 with organelles and PEX5 monoubiquitination and extraction into the supernatant.

In line with our observations in human cells, exogenous addition of recombinant *Hs*PEX39 strongly inhibited *in vitro* import of ^35^S-ACAA1 at low nanomolar concentrations (Figure 3C), whereas PTS1-protein import was not affected (Figure S3B, reactions 3 vs 1). Given that excess *Hs*PEX39 can potently inhibit PTS2-protein import, we wondered whether cells may maintain low levels of the protein. Indeed, in human cell lines originating from various tissues, *Hs*PEX39 appears to be a protein of low abundance as it was undetectable in a whole-cell proteomic analysis of cells from the Cancer Cell Line Encyclopedia,^43^ even though *HsPEX39* mRNA is present (Figure S3C).

Consistent with *Hs*PEX39 overexpression inhibiting PTS2-protein import, yeast overexpressing *Sc*Pex39 proliferated worse than wild-type yeast when grown on oleic acid (Figure 3D), but grew similarly on glucose (Figure S3D). Notably, PEX7 levels were not altered in yeast overexpressing *Sc*Pex39 relative to wild-type cells (Figure S2C, lane 1 vs 5). Interestingly, cellular fractionation of yeast overexpressing *Sc*Pex39 revealed increased Pex7 in the organellar fraction relative to wild-type yeast, but no such effect was observed for PTS2 (Pot1, Gpd1) or PTS1 (Cta1) proteins (Figures 3E and S3E). Based on these findings, we reason that a fraction of Pex7 with its PTS2 cargo is stalled at the peroxisomal membrane - but not imported - in the presence of excess *Sc*Pex39. Collectively, our results with *in vitro* assays, human and yeast cells demonstrate that overexpression of PEX39 impairs PTS2-protein import.

### PEX39 stabilizes the PEX7-PTS2 protein interaction

To gain insight into the mechanisms underlying the biology of PEX39, we utilized our native-PAGE assay with radiolabeled PEX7. Using this approach, we observed that the interactions between PEX7 and PEX5 or PEX7 and PHYH are too weak to be detected by native-PAGE (Figure 4A, lanes 2 and 3, respectively), whereas the trimeric complex PEX7-PEX5-PHYH is stable and readily detected (“*” in Figure 4A, lane 4, see also^37^). Of note, the fact that the addition of recombinant *Hs*PEX39 allowed for a PEX7-PHYH-*Hs*PEX39 complex to form (“&” in Figure 4A, lane 7) demonstrates that *Hs*PEX39 stabilizes the weak interaction between PEX7 and PHYH, thus helping PEX7 bind the PTS2 cargo.

**Figure 4.**
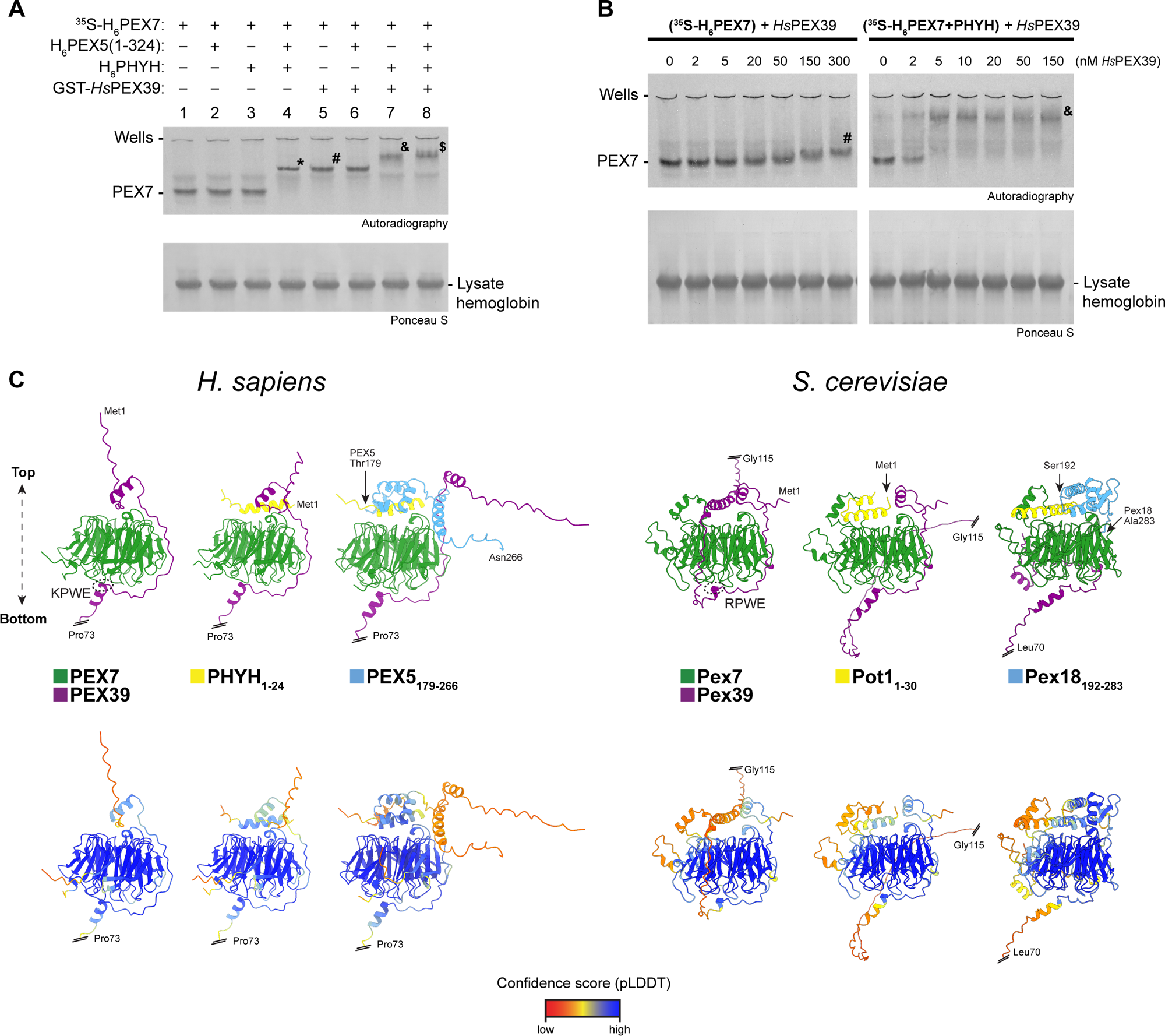
PEX39 stabilizes the PEX7-PTS2 protein interaction. (A) *Hs*PEX39 stabilizes the PEX7-PTS2 protein interaction *in vitro*. Radiolabeled H6PEX7 was pre-incubated or not with the recombinant proteins H6PEX5(1-324), H6PHYH, and/or GST-*Hs*PEX39 as indicated. Samples were analyzed by native-PAGE and autoradiography. In-gel migration of PEX7 alone (PEX7), lysate hemoglobin, PEX7-PEX5-PHYH complex (*), PEX7-*Hs*PEX39 complex (#), PEX7-PHYH-*Hs*PEX39 complex (&), and PEX7-PEX5-PHYH-*Hs*PEX39 complex ($) are indicated. (B) Assessments of KD,app for *Hs*PEX39-PEX7 interaction in different complexes *in vitro*. ^35^S-labeled H6PEX7 was incubated with increasing amounts of recombinant *Hs*PEX39 in the absence (^35^S-H6PEX7) or presence of PHYH (^35^S-H6PEX7+PHYH) as indicated and analyzed by native-PAGE and autoradiography. In-gel migration of PEX7 alone (PEX7), lysate hemoglobin, the PEX7-*Hs*PEX39 complex (#), and PEX7-PHYH-*Hs*PEX39 complex (&) are indicated. (C) AlphaFold predictions of human and yeast PEX39-containing complexes. The top and bottom faces of PEX7 are oriented as indicated. (Top) In the PEX7/PEX39 dimers, the [R/K]PWE motifs of PEX39 are marked with a dotted circle. Structure predictions were performed with full-length PEX39. However, for visualization, PEX39 was C-terminally shortened as indicated by a double line and the last amino acid. (Bottom) AlphaFold confidence scores (*i.e.*, strength of structural predictions) for the corresponding models above are shown according to the predicted local distance difference test (pLDDT). See also Figure S4.

Using this native-PAGE assay, we next determined the apparent dissociation constants (K_D,app_s) for the interactions between *Hs*PEX39 and PEX7 in the presence or absence of PHYH. While these K_D,app_s are not direct measurements of PEX7 and *Hs*PEX39 binding, they can qualitatively offer valuable information. Both the PEX7-*Hs*PEX39 and PEX7-PHYH-*Hs*PEX39 complexes exhibited K_D,app_s for *Hs*PEX39 in the nanomolar range, reflecting high-affinity interactions (Figure 4B). Interestingly, the PEX7-PHYH-*Hs*PEX39 complex formed at lower *Hs*PEX39 concentrations (Figure 4B, right) than those seen for formation of the PEX7-*Hs*PEX39 complex (Figure 4B, left), which supports our observation that *Hs*PEX39 stabilizes the PEX7-PTS2-protein interaction.

To gain further insight into the molecular function of PEX39, we used AlphaFold^44,45^ to predict the structure of *Hs*PEX39 in various complexes with human PEX7, PHYH(1-24) (PTS2-containing portion), and PEX5(179-266) (PEX7-binding residues^46^) (Figures 4C and S4A). These structural predictions corroborate our experimental finding that *Hs*PEX39 stabilizes the interaction between PEX7 and PHYH by predicting that *Hs*PEX39 binds the base of PEX7 via the conserved KPWE motif and the N-terminal half wraps around PEX7 to interact with the PTS2, thereby potentially acting as a clamp to strengthen the interaction between PEX7 and PHYH. Also of note, beginning with the model of the PEX7-*Hs*PEX39 complex, the addition of PHYH(1-24) is predicted to introduce a new binding surface for the N-terminal region of *Hs*PEX39, which agrees with the observed increase in binding affinity between PEX7 and *Hs*PEX39 (*i.e.*, decreased K_D,app_) when PHYH is present (Figure 4B). Interestingly, PEX5(179-266) is also predicted to interact with the PTS2 and displace the N-terminal portion of *Hs*PEX39, but still allows for *Hs*PEX39 to bind PEX7 via the KPWE motif, which agrees with our observation that a PEX7-PHYH-*Hs*PEX39-PEX5 complex can form (Figure 1G). Of note, the structural predictions generated by AlphaFold are consistent with the crystal structure of a yeast Pex7-Pex21-PTS2 complex.^47^

AlphaFold modeling of *Sc*Pex39 in various complexes with yeast Pex7, Pot1(1-30) (PTS2-containing portion), and Pex18(192-283) (Pex7-binding residues^46^) revealed similar predictions (Figures 4C and S4B). The RPWE motif of *Sc*Pex39 is predicted to bind the bottom face of Pex7, with the N-terminal half wrapping around Pex7 to interact with the PTS2 of Pot1, potentially stabilizing the Pex7-Pot1 interaction. In addition, Pex18(192-283) is predicted to interact with the PTS2 and displace the N-terminal portion of *Sc*Pex39 but still allows *Sc*Pex39 to bind Pex7 via the RPWE motif. In summary, our *in vitro* data and structural predictions indicate that PEX39 stabilizes the PEX7-PTS2 protein interaction, thereby providing a mechanism by which PEX39 facilitates PTS2-protein import.

### The N-terminal region and the [R/K]PWE motif are essential for PEX39 function

The evolutionary conservation of the [R/K]PWE motif in PEX39 orthologs (Figure S1A) and the AlphaFold predictions of the motif being a PEX7-binding site (Figure 4C) suggested that it contributes to PEX39 function. To investigate this, we utilized an *Hs*PEX39 variant that had the KPWE motif mutated to AAAA [*Hs*PEX39(4A)], as well as different truncated forms of *Hs*PEX39 that were designed, in part, by insights obtained from the AlphaFold predictions of the various complexes.

Using our native-PAGE assays with ^35^S-PEX7, we examined the ability of these *Hs*PEX39 variants to complex with PEX7, PHYH, and PEX5 (Figure 5A). *Hs*PEX39(4A) could not form any interactions with PEX7 and failed to stabilize the PEX7-PTS2 protein interaction, as the only detectable species were PEX7 alone or the PEX7-PHYH-PEX5 complex (Figure 5A, lanes 6-10). A C-terminally truncated version of *Hs*PEX39 containing residues 1-70 [*Hs*PEX39(ΔC)] could still bind PEX7 and complex with PEX7 and PHYH, as well as PEX7, PHYH, and PEX5 (Figure 5A, lanes 11-15); this is consistent with the deleted C-terminal residues of *Hs*PEX39 not being predicted to be involved in the formation of these protein complexes per AlphaFold (Figure 4C), although we cannot exclude the importance of the deleted residues in other aspects of *Hs*PEX39 biology. Using an N-terminally truncated variant containing residues 48-101 [(*Hs*PEX39(ΔN)], we observed formation of a PEX7-*Hs*PEX39(ΔN) complex (Figure 5A, lane 17). Interestingly, this variant did not form a trimeric complex with PEX7 and PHYH (Figure 5A, lane 18 vs 17), which indicates that the missing N-terminal region (*i.e.*, residues 1-47) is important for the ability of *Hs*PEX39 to stabilize the weak PEX7-PHYH interaction, in agreement with the AlphaFold predictions (Figure 4C). However, *Hs*PEX39(ΔN) seemed to retain the ability to interact with the PEX7-PHYH-PEX5 complex (Figure 5A, compare lanes 18-20), raising the possibility that the KPWE motif is sufficient for this interaction. Consistent with this hypothesis, *Hs*PEX39(ΔN) with a mutated KPWE motif [*Hs*PEX39 (ΔN,4A)] failed to interact with PEX7, PHYH, and PEX5 (Figure 5A, lanes 21 to 25), thus underscoring the importance of the KPWE motif.

**Figure 5.**
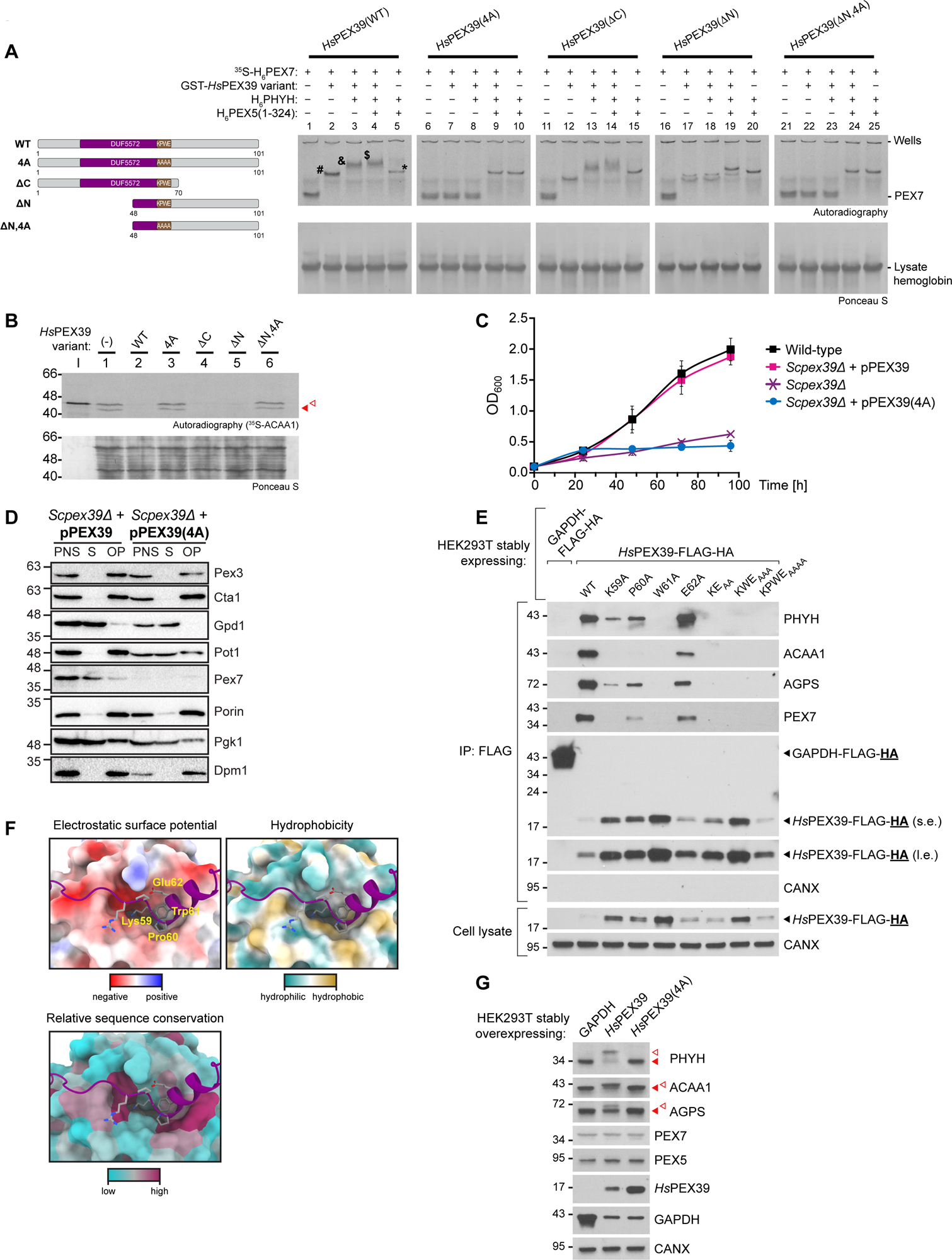
The N-terminal region and the [R/K]PWE motif are essential for PEX39 function. (A) Investigation of *Hs*PEX39 truncation and mutated variants using native-PAGE and radiolabeled PEX7. Depictions of the different variants are shown on the left. ^35^S-H_6_PEX7 was incubated or not with the indicated recombinant proteins and analyzed by native-PAGE and autoradiography. In-gel position of PEX7 alone (PEX7), lysate hemoglobin, and of the complexes PEX7-*Hs*PEX39 (#), PEX7-*Hs*PEX39-PHYH (&), PEX7-PEX5-PHYH-*Hs*PEX39 ($) and PEX7-PEX5-PHYH (*) are indicated for the full-length wild-type *Hs*PEX39 variant. Double bands in *Hs*PEX39(ΔN) complexes are due to co-migration with hemoglobin from the lysate. (B) Investigation of *Hs*PEX39 truncation and mutated variants using an *in vitro* import assay. ^35^S-ACAA1 was subjected to *in vitro* import assays at 37°C in the absence (-) or presence of the indicated recombinant *Hs*PEX39 proteins [see depictions of the indicated variants in left side of (A)]. After incubation, reactions were treated with trypsin and organelles were isolated by centrifugation and analyzed by SDS-PAGE and autoradiography. Precursor and mature forms of ACAA1 denoted by open and solid red arrowheads, respectively. I, 5% of the reticulocyte lysate containing the ^35^S-labeled protein used in each reaction. (C) Mutation of the RPWE motif in *Sc*Pex39 prevents rescue of the fitness defect of *Scpex39Δ* cells grown on oleic acid. Experiment performed as described in Figure 2A analyzing the growth of *Scpex39Δ* cells transformed with a plasmid for expression of an *Sc*Pex39 RPWE-to-AAAA mutant under control of the endogenous promoter [pPEX39(4A)] in oleic acid medium. Data for wild-type, *Scpex39Δ*, and *Scpex39Δ* + pPEX39 are the same as shown in Figure 2A (right plot). Data are mean ± SD (n = 4). Error bars may not be visible if the SD is very small. (D) Cellular fractionation of *Scpex39Δ* yeast expressing wild-type or mutant *Sc*Pex39. Experiment performed as described in Figure 2B using *Scpex39*Δ cells transformed with plasmids for expression of wild-type *Sc*Pex39 (pPEX39) or the *Sc*Pex39 RPWE-to-AAAA mutant [pPEX39(4A)], each under control of the endogenous promoter. PNS, post-nuclear supernatant; S, cytosolic supernatant; OP, organellar pellet. (E) Mutations of the KPWE motif of *Hs*PEX39 have deleterious effects on protein interactions per assessment with HEK293T cells. Anti-FLAG immunoprecipitates and cell lysates were prepared from wild-type HEK293T cells stably expressing the indicated proteins. Samples were analyzed by immunoblotting for the indicated proteins; detection of HA denoted by “<u>HA</u>” in the labeled black arrowheads identifying the corresponding proteins. GAPDH-FLAG-HA is a negative control for the immunoprecipitations. Different *Hs*PEX39 variants denoted as WT (wild-type) or by single or multiple alanine replacements of the indicated residue(s). (F) Close-up views of the interactions between the KPWE motif of *Hs*PEX39 and PEX7. Shown are different surface properties and relative sequence conservation of the *Hs*PEX39 binding region at the bottom face of PEX7. Images are based on the same predicted structural model as shown in Figure 4C (*H. sapiens*). The individual amino acids of the KPWE motif are labeled. (G) Mutation of the KPWE motif prevents overexpressed *Hs*PEX39 from impairing the import of PHYH, ACAA1, and AGPS in human cells. GAPDH (negative control), *Hs*PEX39, or *Hs*PEX39 with KPWE motif replaced by AAAA [*Hs*PEX39(4A)] were stably overexpressed in wild-type HEK293T cells and cellular lysates analyzed by immunoblotting for the indicated proteins. For the PHYH, ACAA1, and AGPS blots, the solid and open red arrowheads indicate the mature and precursor forms of these proteins, respectively. CANX is a loading control. See also Figure S5.

We next investigated the behavior of the *Hs*PEX39 variants using our *in vitro* peroxisomal import assays. Here, all exogenously added variants that had an intact KPWE motif and could bind PEX7 *in vitro* (Figure 5A) almost completely inhibited import of the reporter ACAA1 (Figure 5B, lanes 2, 4, 5), while variants with a mutated KPWE motif and inability to bind PEX7 *in vitro* did not affect import (Figure 5B, lanes 3 and 6). Taken together, these results demonstrate that the KPWE motif is necessary for the inhibition of peroxisomal import *in vitro* by excess *Hs*PEX39.

We next extended our investigations into yeast and human cells. Expression of *Sc*Pex39 with the RPWE motif mutated to AAAA [*Sc*Pex39(4A)] via the endogenous promoter failed to rescue the fitness defect of *Scpex39*Δ cells grown on oleic acid (Figure 5C), while growth on glucose remained unaffected (Figure S5A). *Sc*Pex39(4A) also failed to restore Pex7 levels (Figure S2C, lanes 1-4) and PTS2-protein import in *Scpex39*Δ yeast (Figures 5D and S5B). Using wild-type HEK293T cells, we then examined the effects of mutating single or multiple residues within the KPWE motif on protein interactions (Figure 5E). We observed that alanine replacement of any single residue within the KPWE motif was generally deleterious on the ability of *Hs*PEX39 to interact with its binding partners, with the W61A variant most and the E62A variant least impaired. Consistent with single mutations generally being deleterious, combinations of multiple mutations severely impacted the ability of *Hs*PEX39 to interact with binding partners as well. The AlphaFold structural prediction of the PEX7 binding site and the KPWE motif corroborates the importance of the motif residues by illustrating how conserved PEX7 residues with matching biochemical properties are positioned to interact (Figures 5F and S5C). To confirm the importance of the KPWE motif in human cells, we overexpressed in wild-type HEK293T cells, which also express endogenous *Hs*PEX39, either wild-type *Hs*PEX39 or a variant with the KPWE motif mutated to AAAA [*Hs*PEX39(4A)]. While overexpression of wild-type *Hs*PEX39 increased precursor and decreased mature forms of the PTS2 proteins PHYH, ACAA1, and AGPS, overexpression of *Hs*PEX39(4A) did not (Figures 5G and S5D); importantly, this is not simply due to the mutant protein being unstable and not achieving sufficient levels for inhibition to occur as there is greater overexpression of mutant relative to wild-type *Hs*PEX39. Lastly, overexpression of wild-type or *Hs*PEX39(4A) did not alter levels of PEX7 or PEX5 (Figures 5G and S5E). In summary, our investigation of different regions of PEX39 demonstrates that the [R/K]PWE motif and the N-terminal region are essential for PEX39 function. The N-terminal KPWE motif of PEX13 is evolutionarily linked to PEX39 and is necessary for PEX7 binding and proper peroxisomal biogenesis

As mentioned previously, there is also a highly conserved KPWE motif in the N-terminus of PEX13, including human and yeast orthologs (Figures S1A and 6A). However, this KPWE motif has yet to be experimentally characterized. Interestingly, the presence or absence of PEX39 is mirrored by the presence or absence of the PEX13 KPWE motif across eukaryotes (Figure S1A), strongly suggesting that there is a functional connection between PEX39 biology and the PEX13 KPWE motif throughout evolution. In addition, amongst PEX7 interactors in yeast and humans, the [R/K]PWE motif is almost exclusively found in PEX39 and PEX13 (Figure S6A). These insights are notable for several reasons: firstly, PEX13 is essential for peroxisomal protein import^48^ and is thought to form a conduit to allow proteins to translocate across the peroxisomal membrane;^26,27^ in addition, prior work has shown that the first N-terminal 55 amino acids of Pex13, which contains the KPWE motif, can bind Pex7 and are essential for PTS2-protein import in yeast;^49^ and recently, computational predictions suggested that the PEX13 KPWE motif could bind PEX7.^32^ Given our observations that PEX39 facilitates PTS2-protein import and contains a KPWE motif essential for PEX7 binding and PEX39 function, we hypothesized that the PEX13 KPWE motif is necessary for binding PEX7 and plays an important role in PEX13 function.

To test this hypothesis, we produced a recombinant protein comprised of the first 36 residues of the *Hs*PEX13 N-terminus fused to the small ubiquitin-related modifier 1 (Sumo1) with a hexa-histidine tag (NtPEX13), as well as a variant with the KPWE motif mutated to AAAA [NtPEX13(4A)], and investigated whether these proteins could interact with ^35^S-PEX7 in our native-PAGE assay (Figure 6B). We observed that PEX7 can indeed interact with NtPEX13 (lanes 5 vs 1 and 10 vs 9) and this complex remains unchanged by the presence of either PEX5(1-324) or PHYH (lanes 5-7). Only in the presence of both PEX5 and PHYH do we observe the appearance of a slower migrating complex, although we cannot discern whether this is PEX7-PHYH-PEX5 alone or with PEX13 bound as well (lane 4 vs 8). Importantly, NtPEX13(4A) failed to interact with PEX7 (lane 10 vs 11), thus demonstrating the importance of the KPWE motif for the N-terminus of *Hs*PEX13 to bind PEX7. Using our native-PAGE assay, we also determined the K_D,app_ for the interaction between PEX7 and NtPEX13 to be in the low micromolar range (Figure 6C), which is larger than the one observed for the PEX7-*Hs*PEX39 interaction (Figure 4B, left). However, interpretation of this value should be with the understanding that we are not using full-length *Hs*PEX13 and we have not reconstituted the complete peroxisomal docking and translocation module, in which PEX13 should be present in multiple copies.^19,50^ Consistent with our *in vitro* data, AlphaFold modeling of the interaction between PEX7 and the N-terminus of PEX13 for human and yeast orthologs predicts that the N-terminal KPWE motif binds to the bottom of PEX7 (Figures 6D and S6B). Lastly, given that NtPEX13 shares properties with *Hs*PEX39 (*i.e.*, soluble proteins able to bind PEX7 in KPWE-dependent fashion), we investigated the effects of NtPEX13 on PTS2-protein import and observed that NtPEX13 could indeed inhibit import of ^35^S-ACAA1 *in vitro* in a KPWE-dependent fashion (Figure S6C), but not PTS1-protein import (Figure S6D); this is similar to what we observed for *Hs*PEX39 and provides additional evidence that the role of the KPWE motif may be restricted to PTS2-protein import.

**Figure 6.**
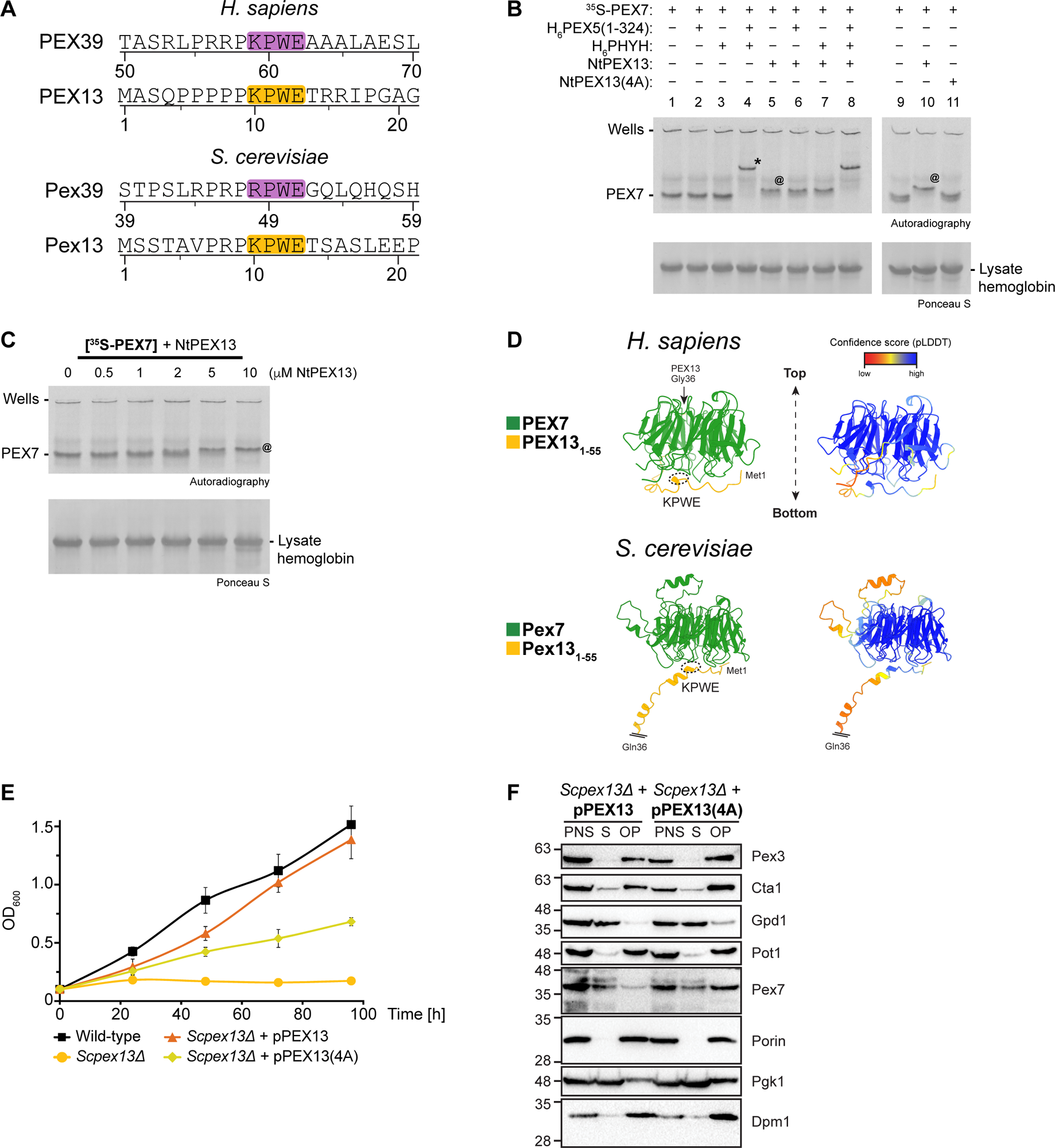
The N-terminal KPWE motif of PEX13 is evolutionarily linked to PEX39 and is necessary for PEX7 binding and proper peroxisomal biogenesis. (A) Schematic demonstrating that PEX39 and the N-terminus of PEX13 both possess [R/K]PWE motifs in yeast and humans. Select portions of the respective amino acid sequences are shown, with the [R/K]PWE motifs highlighted in purple (PEX39) or orange (PEX13). (B) The KPWE motif is necessary for the N-terminus of *Hs*PEX13 to bind PEX7 *in vitro*. Radiolabeled H6PEX7 was pre-incubated with the recombinant proteins as indicated. NtPEX13, first 36 residues of *Hs*PEX13 fused to the N-terminus of the small ubiquitin-related modifier 1 (Sumo1) with a hexa-histidine tag at the C-terminal end; NtPEX13(4A), NtPEX13 with KPWE motif mutated to AAAA. Samples were analyzed by native-PAGE and autoradiography. In-gel migration of PEX7 alone (PEX7), lysate hemoglobin, and of the complexes PEX7-PEX5-PHYH (*) and PEX7-*Hs*PEX13 (@) are indicated. (C) Assessment of KD,app for the interaction between N-terminus of *Hs*PEX13 and PEX7. ^35^S-labeled H6PEX7 was incubated with increasing amounts of NtPEX13 as indicated and analyzed by native-PAGE and autoradiography. In-gel migration of PEX7 (PEX7), lysate hemoglobin, and of the complex PEX7-*Hs*PEX13 (@) are indicated. (D) AlphaFold prediction of interactions between the PEX13 N-terminus and PEX7 in human and yeast. Top and bottom faces of PEX7 are oriented as indicated. (Left) The KPWE motifs of PEX13 are marked with a dotted circle. AlphaFold prediction was performed using full-length sequences of PEX7 and amino acids 1-55 of PEX13. For visualization, PEX13 was C-terminally shortened at residue 36 as indicated. (Right) AlphaFold confidence scores of corresponding models. pLDDT, predicted local distance difference test. (E) Mutation of the N-terminal KPWE motif in *Sc*Pex13 confers a fitness defect in yeast grown on oleic acid. Experiment performed as described in Figure 2A. *Scpex13Δ* cells were transformed with plasmids expressing wild-type *Sc*Pex13 (pPEX13) or a Pex13 mutant in which the KPWE motif was converted to AAAA [pPEX13(4A)] via the endogenous promotor. Data are mean ± SD (n = 4). Error bars may not be visible if the SD is very small. (F) Cellular fractionation of *Scpex13Δ* yeast expressing wild-type or mutant *Sc*Pex13. Experiment performed as described in Figure 2B using cells expressing plasmid-encoded wild-type *Sc*Pex13 (pPEX13) or the KPWE-to-AAAA mutant [pPEX13(4A)] described in (E). PNS, post-nuclear supernatant; S, cytosolic supernatant; OP, organellar pellet. See also Figure S6.

Having determined that the KPWE motif of the PEX13 N-terminus is necessary for binding to PEX7, we investigated the importance of this motif in PTS2-protein import. Expression of *Sc*Pex13 with the KPWE motif mutated to AAAA via the endogenous promoter could not fully rescue the fitness defect of *Scpex13*Δ yeast grown on oleic acid (Figure 6E). Interestingly, cellular fractionation of these yeast revealed an increased amount of Pex7 in the organellar fraction relative to control yeast (Figures 6F and S6E). These results resemble those observed for yeast overexpressing *Sc*Pex39 and we similarly reason that a fraction of Pex7 with its PTS2 cargo is stalled at the peroxisomal membrane - but not imported - upon loss of the *Sc*Pex13 KPWE motif. In summary, our investigation of the KPWE motif of PEX13 reveals that it is necessary for PEX7 binding, proper PEX13 function, and peroxisomal biogenesis.

### Dissociation of PEX39 from PEX7 allows the N-terminus of PEX13 to bind

The presence of [R/K]PWE motifs in both PEX39 and PEX13 that are important for binding to PEX7 suggests that their interaction with PEX7 is mutually exclusive. Indeed, structural modeling predicts that the [R/K]PWE motifs of PEX39 and the N-terminus of PEX13 bind the same site on PEX7 in both human and yeast (Figure 7A). Furthermore, using FLAG-HA-tagged proteins stably expressed in HEK293T cells, we observed that *Hs*PEX39-FLAG-HA did not co-immunoprecipitate *Hs*PEX13, whereas FLAG-HA-tagged PEX7, ACAA1, and PHYH did (Figure 7B); importantly, this was not due to less PEX7 being co-immunoprecipitated with FLAG-HA-tagged *Hs*PEX39.

**Figure 7.**
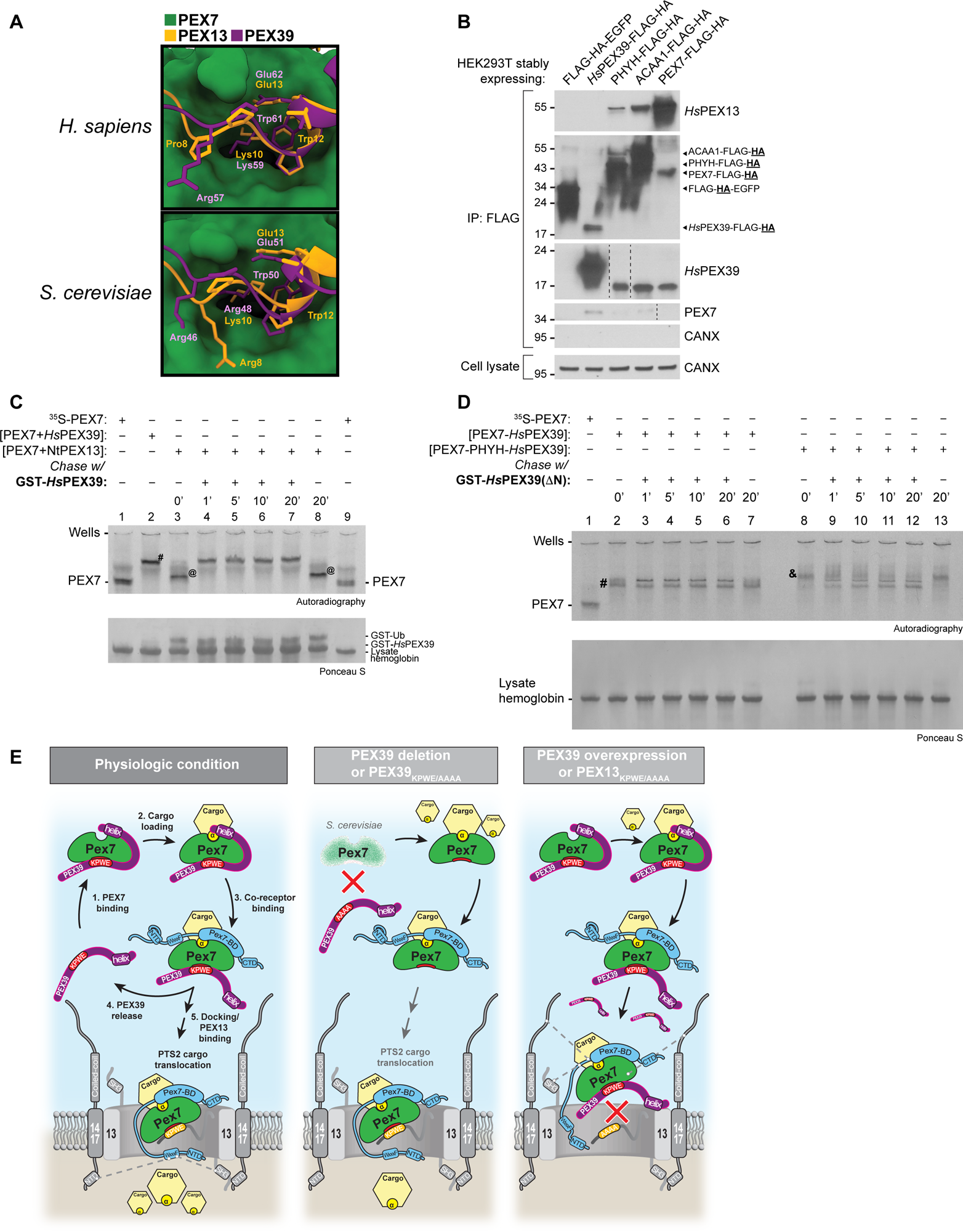
Dissociation of PEX39 from PEX7 allows the N-terminus of PEX13 to bind. (A) Structural modeling predicts that [R/K]PWE motifs of PEX39 and N-terminus of PEX13 both bind to same site of PEX7 in both human and yeast. Shown are superimpositions of the predicted models shown in Figures 4C and 6D. The residues of the PEX39 and PEX13 [R/K]PWE motifs of human and yeast are shown. (B) FLAG-HA-tagged *Hs*PEX39 does not co-immunoprecipitate *Hs*PEX13 per assessment with HEK293T cells. Anti-FLAG immunoprecipitates and cell lysates were prepared from HEK293T cells stably expressing the indicated proteins. Samples were analyzed by immunoblotting for the indicated proteins; detection of HA denoted by “<u>HA</u>” in the labeled black arrowheads identifying the corresponding proteins. Dashed lines indicate where different lanes of the same membrane were brought together; occasionally, it was necessary to have blanks sandwiching a given sample to prevent spillover of immunoblot signal to other samples. (C) *Hs*PEX39 binds to PEX7 and displaces the N-terminus of *Hs*PEX13 with fast kinetics *in vitro*. A mixture of ^35^S-H6PEX7 and NtPEX13 (PEX7+NtPEX13) was incubated with a 5-fold molar excess of GST-*Hs*PEX39 or GST-Ub at 23°C. Aliquots before (0’) and during incubations were collected at the indicated time-points. Radiolabeled PEX7 in a mixture with GST-*Hs*PEX39 (PEX7+*Hs*PEX39) was also analyzed. Samples were analyzed by native-PAGE and autoradiography. In-gel migration of PEX7 alone (PEX7), the complexes PEX7-*Hs*PEX39 (#) and PEX7-NtPEX13 (@), and other proteins are indicated. (D) *Hs*PEX39(ΔN) can rapidly exchange with wild-type *Hs*PEX39 for PEX7 binding *in vitro*. Mixtures of ^35^S-H6PEX7 and GST-*Hs*PEX39 in the absence (PEX7-*Hs*PEX39) or presence of PHYH (PEX7-PHYH-*Hs*PEX39) were incubated with a 100-fold molar excess of either GST-*Hs*PEX39(ΔN) or GST-Ub at 23°C. Aliquots before (0’) and during incubations were collected at the indicated time-points. Samples were analyzed by native-PAGE and autoradiography. In-gel migration of PEX7 alone (PEX7), lysate hemoglobin, and the complexes PEX7-*Hs*PEX39 (#) and PEX7-PHYH-*Hs*PEX39 (&) are indicated. Double bands in PEX7-*Hs*PEX39(ΔN) complexes are caused by co-migration with hemoglobin from the lysate (see also Figure 5A). (E) Model depicting how PEX39 facilitates PTS2-protein import and the consequences of perturbations explored in this study. Proteins and their respective motifs/domains are indicated: cargo with α (PTS2), PTS2 cargo; PEX7-BD, PEX7 binding domain; 13, PEX13; 14/17, PEX14/PEX17; blue, PTS2 co-receptor (*e.g.*, PEX5/Pex18/Pex21); NTD, N-terminal domain; CTD, C-terminal domain; WxxxF, di-aromatic motif. Dashed lines highlight known protein-protein interactions.^49,51,63-66^ See also Figure S7.

To investigate further, we used our native-PAGE assay to assess the time-dependent displacement of NtPEX13 from a PEX7-NtPEX13 complex by GST-*Hs*PEX39. Impressively, NtPEX13 was readily replaced by *Hs*PEX39 even at the first time point (Figure 7C, lane 3 vs 4), while in the absence of *Hs*PEX39, the PEX7-NtPEX13 complex was unchanged during the 20-min incubation (lane 3 vs 8). These results thus indicate that binding of *Hs*PEX39 and the PEX13 N-terminus to PEX7 are mutually exclusive. The fast kinetics by which *Hs*PEX39 replaced NtPEX13 is noteworthy and suggests that the interaction of PEX7 with KPWE motifs may generally be labile.

To further examine the lability of PEX7 binding with KPWE motifs, we assessed the time-dependent replacement of wild-type *Hs*PEX39 in PEX7-*Hs*PEX39 or PEX7-PHYH-*Hs*PEX39 complexes with a distinguishable variant, the aforementioned N-terminally truncated *Hs*PEX39(ΔN) (Figure 7D). For the PEX7-*Hs*PEX39 complex, *Hs*PEX39 was completely replaced by *Hs*PEX39(ΔN) already at the first time point (lane 2 vs 3). For the PEX7-PHYH-*Hs*PEX39 complex, switching of *Hs*PEX39 species was slower (lanes 8-12), which is not unexpected given the lower K_D,app_ for this trimeric complex. Regardless, these results indicate that the interaction of *Hs*PEX39 with PEX7, even with PHYH present, is quite labile. In summary, our results demonstrate that dissociation of PEX39 from PEX7 allows the N-terminus of PEX13 to bind and that this is facilitated by a labile PEX7-PEX39 interaction.

## DISCUSSION

In this work, we identify and characterize PEX39, an ancient and previously unknown component of the peroxisomal protein import machinery, and elucidate a new paradigm in peroxisomal biogenesis, namely the sequential engagement of PEX7 by [R/K]PWE-containing peroxins. Based on the results of this study, we propose a model for how PEX39 facilitates PTS2-protein import, as well as the sequelae of PEX39 loss or overexpression (Figure 7E). In a physiologic condition, PEX39 can directly bind PEX7 via the [R/K]PWE motif (step 1) and then stabilize the PEX7-PTS2 cargo interaction via an N-terminal region (step 2). Binding of a co-receptor (*e.g.*, PEX5/Pex18) to the complex displaces the N-terminal region of PEX39 but independently stabilizes the PEX7-PTS2 cargo interaction (step 3). Because of the labile nature of the PEX39-PEX7 interaction via the [R/K]PWE motif, PEX39 can be exchanged with the N-terminal region of PEX13, which also contains a KPWE motif (steps 4 and 5). PEX13 can now fully participate in the remaining steps required for PTS2-cargo translocation. In the setting of PEX39 loss or mutation of its KPWE motif, the import of PTS2 cargo is impaired due to destabilization of the PEX7-PTS2 cargo interaction, with an additional contribution in *S. cerevisiae* being decreased abundance of PEX7, which suggests that *Sc*Pex39 stabilizes Pex7. Why this is not the case for human PEX7 might stem from different structural properties of the yeast and human orthologs, such as the presence of three loops in yeast but not human PEX7 (Figure S7A). In the setting of PEX39 overexpression or mutation of the PEX13 KPWE motif, PTS2-cargo import is also impaired because even though cargo-loaded, co-receptor-bound PEX7 can be recruited to PEX13, the N-terminus of PEX13 cannot bind PEX7 and facilitate the translocation of PTS2 cargo. The observation that PEX39 impairs import when depleted or overexpressed is surprising for a facilitating factor and is, to the best of our knowledge, unique amongst cytosolic peroxins; by extension, our results suggest that the dosage of PEX39 must be well-controlled to maximize PTS2-protein import.

Based on the insights from our study, we envision multiple important areas for future investigation. A deeper understanding of the mechanistic role of the PEX13 KPWE motif in peroxisomal import will be of significant value, although our proposal that the KPWE motif binds PEX7 to facilitate cargo translocation is consistent with prior studies. Recent data on the membrane topology of PEX13 suggest that at least half of the PEX13 molecules present at the peroxisomal membrane direct their N-termini towards the cytosol.^26,51^ In addition, considering a recent model for the architecture of the docking/translocation module in which the flexible N-terminal domain of PEX13 rests at the center of the protein translocation pore,^6,26^ it is possible that the PEX13 KPWE motif interacts with the cargo-loaded PEX7-coreceptor complex to facilitate cargo translocation.

Structural determination of the PEX39, PEX7, and PEX13 complexes investigated in this study will also provide valuable insight. PEX7 belongs to the large family of WD40 domain proteins, which have three potential interaction sites: the top and bottom face and the circumference, of which interactions with the top and bottom faces are the most and least frequently observed, respectively.^52^ Indeed, the crystal structure of yeast Pex7 reveals that the PTS2 binds to the top face,^47^ but an interactor for the bottom face has yet to be identified. Our experimental data and structural predictions strongly suggest that the [R/K]PWE motifs of PEX39 and PEX13 are interactors of the bottom face and that the function of the bottom face is to allow PEX7 to cycle between these two peroxins.

Additional investigation of the physiological consequences of PEX39 loss will also be of value. *Scpex39*Δ yeast exhibit defects in Pot1 import and grow poorly in conditions requiring fatty-acid oxidation, while *HsPEX39*-knockout cells have impaired import of PHYH, a fatty-acid oxidation enzyme whose deficiency causes adult Refsum disease.^53^ Comparison with prior studies^54,55^ however indicates that loss of PEX7 is more severe than loss of PEX39 in human cells, a phenomenon we also observed in yeast (Figure 2A). Why *Hs*PEX39 loss only affects PHYH and not ACAA1 or AGPS is surprising and may reflect differences in the stability of the corresponding PTS2-PEX7 interactions. Given the role of PHYH in human physiology, it will be important to examine the consequences of PEX39 loss in mice and search for pathogenic human alleles, the latter of which is facilitated by our identification of critical residues in *Hs*PEX39.

Looking beyond yeast and human PEX39, our observation that a subset of eukaryotes, including plants, possesses a PEX39 sequence fused to the C-terminus of PEX14 is fascinating and deserves further investigation. PEX14 is a major component of the docking/translocation module that exposes its C-terminus to the cytosol.^51,56,57^ As such, in these organisms, a pool of PEX7 may be constantly bound to PEX14 via the PEX39 sequence and consequently recognition of PTS2 proteins may occur at the docking/translocation module.

In conclusion, almost all peroxins known up until now were identified using forward genetic approaches in yeast^58^ and mammalian cells.^59^ Although these approaches were extremely successful, they were not designed to identify genes with redundant functions or more subtle loss-of-function phenotypes.^39^ Perhaps for this reason, after the identification of PEX26 more than 20 years ago,^60^ no new human peroxin has been discovered. The work described here breaks this long pause by using functional proteomics instead of functional genomics, and demonstrates both the power of alternative approaches for studying peroxins and the abundance of mysteries still surrounding the import of peroxisomal proteins.

## ACKNOWLEDGMENTS

This article is subject to HHMI’s open access to publications policy. HHMI lab heads have previously granted a nonexclusive CC BY4.0 license to the public and a sublicensable license to HHMI in their research articles. Pursuant to those licenses, the author-accepted manuscript of this article can be made freely available under a CC BY4.0 license immediately upon publication. Work included in this study has been performed in partial fulfilment of the doctoral thesis of D.W., A.Z. and H.D. at the University of Würzburg as well as of the Master thesis of T.M. at the University of Freiburg, all under supervision of B.W. We thank the following people: Ralf Erdmann, Wolfgang Schliebs (Ruhr University Bochum) for scientific discussion, antibodies, and yeast strains; Ann-Kathrin Richard, Chiara Urban for excellent technical assistance; the PRIDE team for data deposition to the ProteomeXchange Consortium; all members of the R.J.D. lab and Hoi See Tsao for helpful suggestions; Wade Harper (Harvard Medical School), in whose lab W.W.C. initially made the link between C6ORF226, PEX7, and PTS2 proteins. This research was funded as follows: AAP Marshall Klaus Perinatal Research Award, Thrasher Research Fund Early Career Award, CMC Fellow Research Scholar Award to W.W.C.; ERC CoG OnTarget (864068) and an Israeli Science Foundation grant 914/22 to M.S.; Portuguese Funds through FCT - Fundação para a Ciência e a Tecnologia, I.P., under the projects UIDB/04293/2020 and 2022.08378.PTDC (doi.org/10.54499/2022.08378.PTDC) to J.E.A.; 2023.02428.RESTART to T.F.; FEDER (Fundo Europeu de Desenvolvimento Regional), through COMPETE 2020-Operational Programme for Competitiveness and Internationalization (POCI), Portugal 2020; Portuguese funds through Fundação para a Ciência e Tecnologia (FCT)/Ministério da Ciência, Tecnologia e Ensino Superior in the framework of the projects: “Institute for Research and Innovation in Health Sciences” (POCI-01-0145-FEDER-007274); T.A.R., A.G.P., T.F, A.R.S. and M.J.F. are supported by Fundação para a Ciência e Tecnologia; Deutsche Forschungsgemeinschaft (DFG, German Research Foundation), Project-ID 278002225 - RTG 2202 and the European Union’s Horizon 2020 research and innovation programme under the Marie Skłodowska-Curie grant agreement No 812968 (PERICO) to B.W.; and the Howard Hughes Medical Institute Investigator Program (R.J.D.).

## AUTHOR CONTRIBUTIONS

Conceptualization, W.W.C., T.A.R., D.W., A.G.P., T.F., S.O., R.J.D., J.E.A., B.W.; methodology, W.W.C., T.A.R., D.W., A.G.P., T.F., R.E.A., V.S., J.E.A., B.W.; software, D.W.; validation, W.W.C., T.A.R., D.W., A.G.P., A.R.S., M.J.F, C.Y., T.M., A.Z., R.E.A.; formal analysis, W.W.C., T.A.R., D.W., A.G.P., T.F., B.N.-S., A.R.S., M.J.F., A.Z., H.D., J.B., S.O., R.E.A., K.H., J.E.A., B.W.; investigation, W.W.C., T.A.R., D.W., A.G.P., T.F., B.N.-S., C.Y., T.M., A.Z., H.D., R.E.A., K.H., H.R.W.; resources, W.W.C., T.A.R., D.W., A.G.P., T.F., B.N.-S., R.E.A., H.R.W., B.W.; data curation, D.W., J.B.; writing - original draft, W.W.C., T.A.R., D.W., A.G.P., M.S., E.Z., K.H., S.O., R.J.D., J.E.A., B.W.; writing - reviewing & editing, W.W.C., T.A.R., D.W., A.G.P., T.F., B.N.-S., A.R.S., M.J.F., M.S., E.Z., S.O., H.R.W., R.J.D., J.E.A., B.W.; visualization, W.W.C., T.A.R., D.W., A.G.P., T.F., R.E.A., K.H., H.D., J.B., B.W.; supervision, W.W.C., T.A.R., D.W., T.F., M.S., E.Z., S.O., R.J.D., J.E.A., B.W.; project administration, W.W.C., R.J.D., J.E.A., B.W.; funding acquisition, W.W.C., T.F., M.S., E.Z., R.J.D., J.E.A., B.W.

## STAR METHODS

### RESOURCE AVAILABILITY

#### Lead contact

Further information and requests for resources and reagents should be directed to and will be fulfilled by the lead contact Walter W. Chen.

#### Materials availability

All unique and stable reagents generated in this study are available upon reasonable request, but may require a completed Materials Transfer Agreement.

#### Data and code availability

- The mass spectrometry proteomics data have been deposited to the ProteomeXchange Consortium via the PRIDE^67^ partner repository with the dataset identifiers PXD051501 (Pex18-TPA affinity purification experiments) and PXD051550 (*Scpex39*Δ yeast SILAC experiments).
- Original code for the analysis of mass spectrometric data is available at https://github.com/ag-warscheid/Pex39_Manuscript and https://doi.org/10.5281/zenodo.10986227.
- Any additional information required to reanalyze the data reported in this paper is available from the lead contact upon request.

### EXPERIMENTAL MODEL AND STUDY PARTICIPANT DETAILS

#### Experiments with yeast not related to fluorescent microscopy

##### Yeast culture conditions and metabolic labeling

Yeast cells were cultured at 30°C and 160 rpm in synthetic complete (SC) medium (pH 6.0) containing 0.17% yeast nitrogen base lacking amino acids, 0.5% ammonium sulfate, 0.3% glucose, 0.002% histidine, methionine, adenine and uracil, 0.003% tryptophane, isoleucine and tyrosine, 0.005% arginine, lysine and phenylalanine, 0.01% leucine, 0.015% valine, and 0.02% threonine (% in w/v each) unless stated otherwise. Depending on the selection markers, uracil, histidine, or both were omitted. To induce peroxisome proliferation, cells were grown in SC medium until an optical density at 600 nm (OD_600_) of 1 - 1.5, shifted to YNO medium (SC medium containing 0.1% oleic acid and 0.05% Tween 40; % in v/v), and cultivated for further 12 - 16 h or as indicated. For metabolic labeling using stable isotope labeling by amino acids in cell culture (SILAC),^68^ the media contained stable isotope-coded ‘heavy’ arginine (^13^C_6_/^15^N_4_, Arg10) and lysine (^13^C_6_/^15^N_2_, Lys8) instead of the unlabeled ‘light’ counterparts (^12^C_6_/^14^N_x_, Arg0/Lys0). Cells were grown for at least six cell doublings in SILAC-SC medium to ensure complete labeling of the proteome with the stable isotope-coded amino acids. For growth of yeast cells in glucose medium, SC medium contained 2% (w/v) glucose. Cells were harvested by centrifugation (10 min, 7,000 g, 4°C) and washed three times with ultrapure water (*i.e.*, Milli-Q quality) before further use unless stated otherwise. For growth assays, strains were pre-cultured in SC medium containing 0.3% glucose. Depending on the selection markers present in the strains, uracil, histidine or both were omitted. Cells were cultivated for 16 h and then shifted to either glucose (2%)- or oleate-containing medium, adjusting the OD_600_ to approximately 0.1. To prevent contamination of the oleate medium with residual glucose from the pre-culture, cells were collected by centrifugation and resuspended in YNO medium. Aliquots of the cultures were taken at distinct time points as indicated and the OD_600_ was determined. Cells grown in YNO medium were washed twice with ultrapure water and resuspended in an appropriate volume of ultrapure water before the OD_600_ was measured.

Yeast cells used for transformation (*i.e.*, in complementation studies and sedimentation assays) were grown overnight in YPD medium (1% yeast extract, 2% peptone, 2% glucose, 0.002% uracil, 0.002% adenine; % in w/v each) at 30°C and 160 rpm, diluted with YPD medium to an OD_600_ of 0.1 in 20 mL, and incubated for further 4 - 5 h at 30°C and 160 rpm until an OD_600_ of 0.4 - 0.7 was reached.

#### Fluorescent microscopy of yeast

##### Yeast growth media

Yeast were grown on synthetic media containing 6.7 g/L yeast nitrogen base with ammonium sulfate (Conda Pronadisa 1545) and 2% glucose (SD) or 0.2% oleic acid (S-oleate) (Sigma-Aldrich) + 0.1% Tween 80 (Sigma-Aldrich) with a complete amino acid mix (oMM composition).^69^ When Geneticin antibiotic was used, the media contained 0.17 g/L yeast nitrogen base without ammonium sulfate (Conda Pronadisa 1553) and 1 g/L of monosodium glutamic acid (Sigma-Aldrich G1626). The strains were selected using a dropout mix (same composition as “SD” above, without the specific amino acid for selection) or with antibiotics using the following concentrations: 500 mg/L Geneticin (Formedium G418), and 200 mg/L Nourseothricin (WERNER BioAgents “ClonNat”).

#### Experiments with human cells

##### Human cell lines

For routine culturing and experiments, all human cell lines were grown at 37°C, 5% CO_2_ and in media supplemented with 10% fetal bovine serum (FBS, GeminiBio 100-106). HEK293T (ATCC CRL-3216) and HCT116 cells (ATCC CCL-247) were grown in Dulbecco′s modified Eagle′s medium with high glucose (Sigma-Aldrich D5796) supplemented with 1 mM sodium pyruvate (Sigma-Aldrich S8636). CAKI-2 cells (gift from Gregory Wyant and William Kaelin Jr.) were grown in McCoy’s 5A (modified) medium (Gibco 16600-082) and NCI-H1792 cells (ATCC CRL-5895) were grown in RPMI-1640 medium (Sigma-Aldrich R8758). All complete media was passed through a 0.22 μm filter prior to use. Cells were routinely tested for mycoplasma.

#### Experiments with radiolabeled proteins

##### Human cell lines

For routine culturing and experiments, HEK293 cells were grown at 37°C, 5% CO2 and in Minimum Essential Medium Eagle (Sigma-Aldrich M2279) supplemented with 2 mM L-alanyl-L-glutamine dipeptide (Gibco 35050061), 10% fetal bovine serum (FBS, Gibco GA3160802), and MycoZap Plus-CL (Lonza 195261). All complete media was passed through a 0.22 μm filter prior to use. Cells were routinely tested for mycoplasma.

### METHOD DETAILS

#### Experiments with yeast not related to fluorescent microscopy

##### Yeast strains and plasmids

Yeast strains and plasmids used in this study are listed in the key resource table. Oligonucleotides used to generate yeast strains and plasmids are listed in Table S1. Genomic manipulation of yeast cells was performed by homologous recombination following transformation of the cells with the respective PCR product. For genomic C-terminal tagging of Pex18 with the tobacco etch virus protease (TEV) cleavage site and Protein A (*i.e.*, the TPA tag), the DNA sequence coding for the TPA tag followed by the selection marker kanMX4 was inserted at the 3’ end of the *pex18* gene.^70^ To generate the *Scpex39* deletion strain, the *yjr012c/Scpex39* gene was replaced with the URA3 marker cassette.^71^ Transformants were selected for the respective marker, and correct integration of the TPA tag and disruption of the *Scpex39* gene, respectively, were confirmed by PCR and sequencing of the PCR products.

To generate the plasmid pRS313-*Sc*Pex39 (P797) for expression of *Sc*Pex39 at endogenous levels, the *Scpex39* ORF and the promoter and terminator regions were amplified from yeast genomic DNA using the primer pair SF1/SF2 and cloned into pRS313^72^ using the SalI restriction enzyme.

The plasmid pRS313-TEF2p-*Sc*Pex39 for *Sc*Pex39 overexpression (P798) using the strong TEF2 promoter was generated by inserting the ORF of *Scpex39*, amplified using the primers SF3 and SF4, into a modified pRS313 backbone comprising the TEF2 promoter and ADH1 terminator. Correct integration was confirmed by sequencing.

The amino acids of the RPWE motif of *Sc*Pex39 in pRS313-*Sc*Pex39 (P797) were mutated to AAAA by site-directed, ligase-independent PCR-mediated mutagenesis.^73^ To this end, two primer pairs (O2145/O2143 and O2144/O2142) were used to amplify the P797 plasmid DNA by PCR. A 1:1 mixture of DpnI-treated O2145/O2143 and O2144/O2142 PCR products was mixed with hybridization buffer (50 mM Tris, 300 mM NaCl, 20 mM EDTA, pH 8.0), and hybridization was performed in a PCR cycler (two cycles of 99°C for 3 min, 65°C for 5 min, and 30°C for 40 min, followed by cool down to 4°C). *E. coli* cells were transformed with hybridization mixture using the heat-shock method, and successful mutagenesis, resulting in the plasmid pRS313-*Sc*Pex39(RPWE/AAAA mutant) (P815), was confirmed by sequencing.

The plasmids pRS313-*Sc*Pex13 (P801) and pRS313-*Sc*Pex13(KPWE/AAAA mutant) (P807) were generated as described for P797 and P815 using the following primer pairs: SF5/SF6 (introducing a XhoI restriction site) as well as O2181/O2179 and O2180/O2178.

##### Transformation of yeast cells

Yeast cells were grown in YPD medium as described above in “Yeast culture conditions and metabolic labeling,” harvested by centrifugation (5 min, 500 g, room temperature), washed first with 20 mL ultrapure water and then with 10 mL SORB (100 mM lithium acetate, 10 mM Tris-HCl, pH 8.0, 1 M sorbitol). The cell pellet was resuspended in 360 μL of SORB, mixed with 40 μL denatured salmon sperm, and directly used for transformation applying the heat-shock method.^70^ For this, 10 μL of the cell suspension were mixed with 100 - 200 ng of purified plasmid DNA in ultrapure water and 66 μL of PEG solution [100 mM lithium acetate, 10 mM Tris-HCl (pH 8.0), 1 mM EDTA (pH 8.0), 40% (w/v) PEG 3350] and incubated for 30 min at room temperature. Next, 9 μL of DMSO were added and the cell suspension was incubated for 15 min at 42°C. Subsequently, the cell suspension was cooled for 2 min on ice, and cells were plated onto an SC agar plate lacking histidine, uracil or both, depending on the selection marker(s) of the plasmid used for transformation. Plates were incubated at 30°C for 3 - 4 days and stored at 4°C until use.

##### Affinity purification of Pex18 complexes from yeast

Native Pex18 complexes were affinity-purified using Pex18-TPA-expressing yeast cells grown under peroxisome-proliferating conditions as described before with some modifications.^19,74^ Isogenic cells expressing the native, non-tagged version of Pex18 and grown under the same conditions served as control. Cells were lysed in a buffer consisting of 20 mM HEPES/80 mM NaCl (pH 7.5) supplemented with protease and phosphatase inhibitors [8 μM antipain, 0.3 μM aprotinin, 1 mM benzamidine, 1 μM bestatin, 10 μM chymostatin, 5 μM leupeptin, 1.5 μM pepstatin A, 1 mM phenylmethylsulfonyl fluoride (PMSF), 9.5 mM NaF] using glass beads and an MM400 mixer mill (Retsch, Haan, Germany) at 20 Hz for 8 min, 4°C. Glass beads and cell debris were removed by two rounds of centrifugation (10 min, 2,000 g, 4°C). Following centrifugation for 90 min at 110,000 g at 4°C, supernatants were carefully removed, supplemented with glycerol [10% (v/v) final concentration], and mixed with human IgG-coupled Sepharose (37.5 μL of IgG Sepharose beads per 50 mg of protein) equilibrated with lysis buffer. Samples were incubated overnight at 4°C with slight agitation. Beads were collected by centrifugation (5 min, 100 g, 4°C), transferred to MobiCols, and washed with elution buffer [same as lysis buffer containing 10% (v/v) glycerol] to remove proteins nonspecifically bound to the beads. Pex18 complexes were eluted by incubation with AcTEV protease in twice the bead volume of elution buffer (1 Unit AcTEV per μL of IgG Sepharose beads) for 2.5 h at 16°C and 1,100 rpm. Eluted proteins were collected by centrifugation (2 min, 100 g, 4°C). To increase the yield, the beads were resuspended in twice the bead volume of elution buffer and incubated for 10 min at 1,100 rpm. The samples were collected again by centrifugation as described above and combined with the corresponding eluate. The experiment was performed in three biological replicates.

##### Preparation of yeast cell lysates

Oleate-induced *Scpex39*Δ and isogenic wild-type cells, labeled with ‘light’ or ‘heavy’ arginine and lysine, were harvested by centrifugation (2 min, 1600 g, 4°C) and washed twice with ultrapure water. Cells were resuspended in 500 μL of lysis buffer (8 M urea, 75 mM NaCl, 50 mM Tris-HCl, 1 mM EDTA, pH 8.0), and equal amounts of differentially labeled *Scpex39*Δ and wild-type cells were mixed based on the cell wet weight. Cells were lysed using glass beads (300 mg) and a Minilys homogenizer (Bertin Technologies, Montigny-le-Bretonneux, France), applying two cycles of 4 min at 5,000 rpm with at least 2 min cooling on ice between cycles. Glass beads and cell debris were removed by centrifugation (5 min, 15,000 g, 4°C). The protein concentration of the cell lysates was adjusted to 1 μg/μL using urea buffer [8 M urea in 50 mM ammonium bicarbonate (AmBic)]. The experiment was performed in four independent replicates with light/heavy label switch.

##### Tryptic in-gel digestion

Affinity-purified Pex18 protein complexes and proteins of the control purifications were acetone-precipitated and resuspended in 0.1 M NaOH/1% (w/v) SDS. Proteins were separated by SDS-PAGE using NuPAGE BisTris gradient gels (4 - 12%) and visualized by colloidal Coomassie Blue staining. Lanes were cut into 11 slices, followed by destaining of the gel slices by alternate incubation with 10 mM AmBic and 5 mM AmBic/50% (v/v) ethanol for 10 min at room temperature. The destaining step was performed three times. To reduce cysteine residues, gel slices were incubated with 10 mM DTT/10 mM AmBic for 45 min at 56°C, followed by alkylation of the thiol groups using 55 mM 2-iodoacetamide/10 mM AmBic (30 min at room temperature). Gel slices were washed with 10 mM AmBic and 100% ethanol (10 min at room temperature each) and dried *in vacuo*. Proteolytic digestion with trypsin (dissolved in 10 mM AmBic; 60 - 120 ng of trypsin per slice depending on the staining intensity) was performed overnight at 37°C. Peptides were eluted by incubating the gel slices for 10 min with 0.05% (v/v) trifluoroacetic acid (TFA)/50% (v/v) acetonitrile (ACN) in an ice-cooled ultrasonic bath. This step was repeated once. Eluted peptides of each sample were combined, dried *in vacuo*, desalted using StageTips^75^ and dried again *in vacuo*.

##### Proteolytic in-solution digestion

Proteins (300 μg) of yeast whole-cell lysates prepared from SILAC-labeled *Scpex39*Δ and wild-type cells were digested in solution using LysC and trypsin as described before^76^ with slight modifications. In brief, cysteine residues were reduced using 5 mM Tris(2-carboxy-ethyl)phosphine (30 min at 37°C) and subsequently alkylated with 50 mM iodoacetamide (30 min at room temperature in the dark). The reaction was quenched by adding DTT (25 mM final concentration) and the urea concentration was adjusted to 1.6 M using 50 mM AmBic. Proteins were first digested with LysC, added at a protease-to-protein ratio of 1:100 (1 h at 37°C and 1,400 rpm), followed by digestion with trypsin (protease-to-protein ratio of 1:50; overnight at 37°C and 1,400 rpm). Proteolytic reactions were stopped by addition of 100% TFA at a final concentration of 2% (v/v). Peptides were subsequently desalted using C18-SD 7 mm/3 mL extraction disc cartridges as described previously^77^, dried *in vacuo*, and further fractionated by high pH reversed-phase liquid chromatography (RP-LC).

##### High pH reversed-phase liquid chromatography

Peptide fractionation by high pH RP-LC^78^ was performed essentially as described before.^77^ Dried peptides were reconstituted in 200 μL of 1% (v/v) ACN/10 mM NH_4_OH (pH 10) by sonication, followed by centrifugation (5 min, 12,000 g, room temperature) to remove insoluble material, and further purification using 0.2 μm PTFE membrane syringe filter (Phenomenex, Aschaffenburg, Germany). Peptides were separated using an Ultimate 3000 HPLC system (Thermo Fisher Scientific, Dreieich, Germany) operated with an NX 3u Gemini C18 column (150 mm x 2 mm inner diameter, particle size of 3 μM, pore size of 110 Å; Phenomenex) at 40°C and a flow rate of 200 μL/min. Peptide elution was performed using a binary solvent system consisting of 10 mM NH_4_OH (solvent A1) and 90% (v/v) ACN/10 mM NH_4_OH (solvent B1). Peptides were loaded onto the column at 1% solvent B1 for 5 min and separated by increasing the percentage of solvent B1 from 1 - 40% in 37 min and 40 - 78% in 3 min, followed by 5 min at 78% B1, before the column was re-equilibrated with 1% solvent B1. Starting at min 1.5 until min 65.5, 45-sec fractions were collected in a concatenated manner resulting in a total of 8 fractions per sample. Peptides were dried *in vacuo*, desalted using StageTips^75^ and dried again.

##### LC-MS analysis

Prior to LC-MS analysis, dried peptides were resuspended in 0.1% (v/v) TFA and insoluble material was removed by centrifugation (12,000 g, 5 min, room temperature). Peptides were analyzed by nano-HPLC-ESI-MS/MS using a Q Exactive Plus mass spectrometer (Thermo Fisher Scientific, Bremen, Germany) connected to an UltiMate 3000 RSLCnano HPLC system (Thermo Fisher Scientific, Dreieich, Germany). The RSLC system was operated at 40°C with C18 trapping columns (μPAC^TM^, 10 mm in length x 2 mm inner diameter; PharmaFluidics, Ghent, Belgium) at a flow rate of 10 μL/min and a C18 endcapped analytical column (μPAC^TM^, 500 mm x 0.3 mm; PharmaFluidics) at a flow rate of 300 nL/min. A binary solvent system consisting of 0.1% (v/v) formic acid (FA) (solvent A2) and 86% (v/v) ACN/0.1% (v/v) FA (solvent B2) was employed for peptide separation. For the analysis of peptides from Pex18-TPA affinity purification experiments, peptides were loaded onto the precolumn, washed and preconcentrated for 3 min at 1% solvent B2 and eluted by applying the following gradient: 1 - 4% B2 in 2 min, 4 - 25% B2 in 20 min, 25 - 44% B2 in 11 min, 44 - 90% B2 in 2 min, and 4 min at 90% B2. The same solvent system was used for the analysis of samples from *Scpex39*Δ-versus-wild-type SILAC experiments. Peptides equivalent to 1 μg of protein were loaded, washed and preconcentrated for 5 min using 1% solvent B2. For peptide elution, a gradient ranging from 1 - 5% B2 in 3 min, 5 - 22% B2 in 103 min, 22 - 42% B2 in 50 min, and 42 - 80% B2 in 5 min was applied.

Mass spectrometric data were acquired in data-dependent mode. MS spectra were recorded in a mass-to-charge (*m/z*) range of 375 - 1,700 with a resolution of 70,000 (at *m/z* 200). The automatic gain control (AGC) was set to 3 x 10^6^ and the maximum injection time (IT) to 60 ms. The 12 most intense precursor ions (*z* ≥ +2) were selected for fragmentation by higher-energy collisional dissociation applying a normalized collision energy of 28%, a resolution of 35,000, and AGC of 10^5^, a maximum IT of 120 ms, and a dynamic exclusion time of 45 s.

##### MS data processing and analysis

Proteins present in Pex18-TPA pulldowns were identified using the Andromeda search engine^79^ implemented in MaxQuant (version 2.0.1.0)^80^ through comparison with the UniProt reference proteome of *S. cerevisiae* (including isoforms, release 02/2024, 6079 entries) and the default contaminants list of MaxQuant, extended by the sequences of the TEV protease and immunoglobulins used in this study. Trypsin/P (no cleavage before proline) was specified as digestion enzyme, oxidation of methionine and N-terminal acetylation as variable modifications, and carbamidomethylation of cysteine as fixed modification. Identifications were transferred between samples using match between runs with standard settings. For subsequent analysis, protein quantities were reported as intensity-based absolute quantification (iBAQ) values from MaxQuant. To ensure that only reliably identified protein groups were reported, the minimum number of unique peptides per protein groups was set to 1.

The autoprot Python module (v0.2)^81^ was used for data analysis and processing. Decoy and contaminant entries as well as protein groups without quantitative information were removed. Moreover, protein groups with a sequence coverage of less than 10% were removed. For protein groups from each gel slice of Pex18-TPA-expressing or control cells, the median log_10_ iBAQ value was calculated. To maximize instrument sensitivity, the maximum sample amount within linear quantification range was injected for each gel slice for Pex18-TPA and control samples. To correct for this, the mean of medians between Pex18-TPA and control cells was calculated for each corresponding gel slice pair and subtracted from all iBAQ values of both slices. The median-corrected iBAQ values were exponential-transformed to gain non-log values and iBAQ values for protein groups of all slices of a replicate were summed. Protein groups with less than two valid values in the Pex18-TPA replicates were removed (missing values in the control samples were not considered as the control is expected to lack specific binders). Missing intensity values were imputed by drawing random values from a distribution matching the intensity distribution of the existing values but shifted by 1.8 standard deviations and scaled to 30% width. Statistical significance of differences in protein group intensity was computed using the ranksum test implemented in the R package RankProd^82^ and results were visualized in autoprot. Data analysis was documented as Jupyter notebook and is available at https://github.com/agwarscheid/Pex39_Manuscript and https://doi.org/10.5281/zenodo.10986227. Information about proteins identified and quantified in Pex18-TPA affinity purification experiments are given in Table S2.

For protein identification and SILAC-based relative quantification in *Scpex39*Δ- versus-wild-type experiments, MaxQuant/Andromeda version 2.4.4.0 was employed. MS/MS data were searched against the *S. cerevisiae* reference proteome provided by *Saccharomyces* Genome Database (SGD; http://sgd-archive.yeastgenome.org/sequence/S288C_reference/orf_protein; downloaded 08/2023) using MaxQuant default parameters with the following exceptions: Arg10/Lys8 was selected as ‘heavy’ label, multiplicity was set to 2; Trypsin and LysC were set as proteolytic enzymes, allowing a maximum of 3 missed cleavages, and the options ‘match between runs’ and ‘requantify’ were enabled. ‘Requantify’ enables the determination of peptide abundance ratios in cases when only one variant of a peptide, either ‘light’ or ‘heavy’, is present. This is the case, for example, for *Sc*Pex39 peptides, which are absent in *Scpex39*Δ cells. Using ‘requantify’, the algorithm assigns a peptide intensity for the missing counterpart from the background signals in MS spectra at the expected *m/z* value. Protein identification required the detection of at least one unique peptide with a length of at least seven amino acids. A false discovery rate of 1% was applied at both the peptide and the protein level. Protein quantification was based on unique peptides and at least one ratio count. ‘Autoprot’ was used for further data analysis and visualization, considering only proteins quantified in at least three out of four replicates, except for *Sc*Pex39, which was quantified in only one replicate. To include *Sc*Pex39, which is only present in wild-type cells, in our data analysis and visualization, missing *Sc*Pex39 ratios were imputed by randomly drawing values deviating ± 0.05 from the single log_2_-transformed value reported by MaxQuant. The normalized protein abundance ratios calculated by MaxQuant were log_2_-transformed, followed by sequential imputation of missing values and cyclic-loess normalization.^83^ To identify proteins with differences in protein abundance between wild-type and *Scpex39*Δ cells, the ‘linear models for microarray data’ (limma) approach was employed. This method is a moderated two-sided t-test that adjusts a protein’s variance in ratios between replicates towards the average ratio variance of the entire dataset.^84,85^ P values were corrected for multiple testing according to Benjamini and Hochberg.^86^ Data analysis was documented as Jupyter notebook and is available at https://github.com/ag-warscheid/Pex39_Manuscript and https://doi.org/10.5281/zenodo.10986227. Information about proteins identified and quantified in *Scpex39*Δ-versus-wild-type SILAC experiments are provided in Table S3.

##### Cellular fractionation (sedimentation assay)

Freshly harvested oleate-induced yeast cells (5 g per experiment) were resuspended in 25 mL of DTT buffer (10 mM DTT in 100 mM Tris) and incubated for 20 min at 37°C and 60 rpm in a 250 mL Erlenmeyer flask. Cells were harvested by centrifugation (10 min, 600 g, room temperature), resuspended in fresh DTT buffer, and incubated again as described above. Cells were harvested and washed three times with 20 mL of 1.2 M sorbitol (in ultrapure water) preheated to 37°C. To prepare spheroplasts, cells were then resuspended in 35 mL of 1.2 M sorbitol buffer (1.2 M sorbitol in 20 mM potassium phosphate, pH 7.4, preheated to 37°C) containing 1000 U of lyticase per g cell wet weight, transferred to a 50 mL reaction tube and incubated for 30 min at 37°C and 60 rpm. Digestion of the cell wall was stopped by incubation on ice for 10 min, inverting the reaction tubes two to three times in between. Spheroplasts were washed three times with 15 mL of precooled 1.2 M sorbitol and collected by centrifugation (10 min, 600 g, 4°C) after each washing step. Spheroplasts were resuspended in 5 mL of homogenization buffer (5 mM MES, 0.5 mM EDTA, 1 mM KCl, 0.6 M sorbitol, pH 6.0) containing protease and phosphatase inhibitors (same set as described above for the Pex18-TPA affinity purification) using a Dounce homogenizer operated at 2x 100 rpm (2 min each), 3x 300 rpm (1 min each), 3x 500 rpm (1 min each), and 3x 800 rpm (1 min each). Cell debris and nuclei were removed by centrifugation (2x 10 min, 600 g, 4°C). Protein concentrations of the resulting post-nuclear supernatants (PNS) obtained from different strains within given experiments were adjusted using homogenization buffer. The PNS (1 mL) was loaded on top of a 200-μL sucrose cushion (0.5 M sucrose in homogenization buffer) and separated into an organellar pellet and a cytosolic fraction by centrifugation (20 min, 25,000 g, 4°C). The organellar pellet was resuspended in 1 mL of homogenization buffer. Equal volumes of PNS, cytosolic fraction, and organellar pellet were analyzed by SDS-PAGE and semi-dry Western blotting.

##### SDS-PAGE and immunoblotting

SDS-PAGE and semi-dry Western blotting were performed following standard protocols unless specified otherwise. Antibodies used for immunoblotting are listed in the Key Resources Table.

##### Quantification of immunoblots

Immunoblot signals were quantified using the software ImageJ (version 1.54d).^87^ Signal intensities were corrected for background intensities.

#### Fluorescent microscopy of yeast

##### Yeast strain construction

Genetic manipulations were performed using PCR-mediated homologous recombination with the lithium-acetate method.^88^ The correct tagging or deletion were verified in all strains by PCR. The primers in this study were either designed using the web tool Primers-4-Yeast^89^ or manually constructed in the case of the deletion of *Scpex39*. All relevant strains and plasmids are listed in the Key Resources Table and all relevant primers in Table S1.

##### PCR validation of genomic transformations

Freshly grown yeast cells were picked from agar plates and suspended in PCR tubes containing 50 μL of 20 mM NaOH with 0.1 mg/mL RNaseA. The suspension was then boiled at 100°C for 20 min in a PCR machine and spun down at a micro-centrifuge for 3 min. The supernatant was used as template DNA for a PCR reaction (2 μL), alongside 2X GoTaq Green Master Mix (5 μL, Promega), forward primer (0.2 μL from 10 μM concentration), reverse primer (0.2 μL from 10 μM concentration) and DDW up to a final volume of 10 μL. The DNA was amplified using the following thermocycling steps: 98°C for 3 min; 35 cycles of 98°C for 60 sec, 55°C for 90 sec and 72°C for 30 sec; 72°C for 60 sec. The resulting PCR product was then run on a 1% agarose gel and examined for the correct size of the amplicon.

##### Imaging of yeast strains

Yeast strains were grown overnight in an SD-based medium supplemented with amino acids in 96-well polystyrene plates and were then transferred to S-oleate for 4 h (for experiment shown in Figure S1C) or 8 h (for experiment shown in Figure 2D). The strains were then manually transferred into glass-bottom 384-well microscope plates (Matrical Bioscience) coated with Concanavalin A (Sigma-Aldrich). After 20 min, the wells were washed twice with DDW to remove non-adherent cells and obtain a cell monolayer. Imaging was performed in DDW. Images were taken using the Olympus IXplore SpinSR system, composed of an Olympus IX83 inverted microscope scanning unit (SCU-W1) operated by ScanR. When high-resolution images were taken (*i.e.*, Figure 2D), a high-resolution spinning disk module (Yokogawa CSU-W1 SORA confocal scanner with double microlenses and 50 μm pinholes) was used. Cells were imaged using an X60 oil lens (NA 1.42) and Hamamatsu ORCA-Flash 4.0 camera. Images were recorded in two channels: mNeonGreen (excitation wavelength 488 nm) and mScarlet (excitation wavelength 561 nm). For all micrographs, a single, representative focal plane was imaged and shown.

#### Experiments with human cells

##### Reagents and antibodies

Materials were obtained from the indicated sources: puromycin (InvivoGen ant-pr-1); doxycycline hyclate (Thermo Scientific Chemicals 446060050); antibodies to ACAA1 (HPA006764), AGPS (HPA030211), C6ORF226 (HPA045350), PEX5 (HPA039260), *Hs*PEX13 (HPA032142) were from Sigma-Aldrich; the antibody to ACTB (sc-69879) was from Santa Cruz Biotechnology; antibodies to PEX7 (20614-1-AP), PHYH (12858-1-AP), SCP2 (23006-1-AP) were from Proteintech; antibodies to CANX (2433), CS (14309), GAPDH (2118), HA (3724), Histone H3 (3638), and RPS6KB1 (2708), as well as HRP-conjugated anti-rabbit secondary antibody (7074) were from Cell Signaling Technology; HRP-conjugated anti-mouse secondary antibody for IP (ab131368) was from Abcam.

##### Genetic modification of human cells

pHAGE vectors encoding the following were obtained as indicated: C-terminally FLAG-HA-tagged human ACAA1, PHYH, PEX7, and wild-type *Hs*PEX39 were obtained from the human orfeome collection (version 8); C-terminally FLAG-HA-tagged human GAPDH and KPWE mutants of *Hs*PEX39 were generated by cloning gblocks (IDT) containing the corresponding cDNA into a pHAGE vector containing a C-terminal FLAG-HA tag [linearized with BsrGI-HF (NEB R3575)] via NEBuilder HiFi DNA assembly (NEB E2621); N-terminally FLAG-HA-tagged EGFP was a gift from Alban Ordureau; human GAPDH, wild-type and KPWE-mutated *Hs*PEX39 were generated by cloning gblocks containing the corresponding cDNA into a pHAGE vector (linearized with BsrGI-HF) via NEBuilder HiFi DNA assembly. An sgRNA against *AAVS1* (sg*AAVS1*) and 3 independent sgRNAs against *HsPEX39* (sg*HsPEX39*_1-3) were cloned into a lentiCRISPR-v2 vector (TLCV2, doxycycline-inducible Cas9, Addgene 87360^90^) linearized with BsmBI-v2 (NEB R0739) by ligation of annealed oligonucleotides (IDT) with Quick Ligase (NEB M2200). The oligonucleotide sequences used for the sgRNAs are provided in Table S1.

For the production of lentivirus, each of these constructs was transfected with the lentiviral packaging vectors psPAX2 and pMD2.G into HEK293T cells using PolyJet DNA transfection reagent (SignaGen Laboratories SL100688). Media was changed 24 h after transfection. The virus-containing supernatant was collected 48 h after transfection and passed through a 0.45 μm filter to eliminate cells.

The desired cell lines were then infected with lentivirus added to complete culture media containing 8 μg/ml of polybrene (Sigma-Aldrich TR-1003). 24 h after infection, the media was changed. 48 h after infection, cells were selected with puromycin (1 μg/mL for all cell lines except CAKI-2, which were 2 μg/mL) until an uninfected control was completely dead, at which point infected cells were freed from selection.

CAKI-2 and NCI-H1792 cells transduced with lentivirus harboring TLCV2 with sg*AAVS1* (negative control) or the 3 independent sgRNAs sg*HsPEX39*_1-3 were used for our lentiviral, doxycycline-inducible CRISPR-Cas9 system. After puromycin selection, these cells were treated with 1 μg/mL doxycycline hyclate, which stimulates the production of Cas9 and EGFP. Cells were treated with doxycycline hyclate for 9 days to achieve significant depletion of *Hs*PEX39 across the population before the drug was removed and cells were subsequently used for experimentation.

To generate *HsPEX39* knockouts with matched controls, we took CAKI-2 and NCI-H1792 cells with sg*AAVS1* or sg*HsPEX39*_1 on the 9th day of treatment with doxycycline hyclate and sorted single-cell clones into wells with conditioned media using FACS based on EGFP signal. The composition of this conditioned media was a 1:1 mixture of fresh and spent complete culture media with no doxycycline hyclate that was supplemented with 1% Penicillin-Streptomycin (Sigma-Aldrich P0781) and additional FBS to achieve a final concentration of 20%. Single-cell clones were expanded for 2 - 3 weeks and visible colonies for cells with sg*HsPEX39*_1 were screened by immunoblotting for the absence of *Hs*PEX39. Equal amounts of cells originating from 8 *HsPEX39*-knockout clones were then pooled together to make the final knockout line (*i.e.*, diversifying the population of cells). Equal amounts of cells originating from 8 clones with sg*AAVS1* were also pooled together to make the final matched controls. CAKI-2 and NCI-H1792 cells were chosen for their excellent performance in our workflow for isolating and expanding single-cell clones, and for their greater expression of *HsPEX39* and/or *PHYH*, *ACAA1*, and *AGPS* compared to HCT116 and HEK293T cells.

##### Anti-FLAG immunoprecipitation using human cells

The immunoprecipitation workflow was done at 4°C or on ice unless indicated and low-retention microcentrifuge tubes (Thermo Fisher Scientific 3448) were used. Fully confluent 10-cm dishes of HCT116 or HEK293T cells stably expressing FLAG-HA-tagged proteins were washed once with ice-cold Dulbecco’s Phosphate-Buffered Saline without calcium and magnesium, pH 7.4 (DPBS, Corning 21-031-CM), and immediately scraped into 500 μL lysis buffer [50 mM Tris-HCl, pH 7.5, 150 mM NaCl, 0.5% (v/v) NP40 alternative (Millipore 492016), one tablet cOmplete protease inhibitor cocktail (Roche 04693116001) per 25 mL lysis buffer, one tablet PhosSTOP (Roche 04906837001) per 10 mL lysis buffer, and 1 mM DTT (Sigma-Aldrich 43816)]. Lysates were incubated with rotation for 30 min at 4°C before being cleared by centrifugation at 17,000 g for 10 min at 4°C. Protein quantification was performed on the clarified lysate using the *DC* protein assay (Bio-Rad 500-0116). Anti-FLAG M2 magnetic beads (Millipore M8823) were washed 3 times with 10X packed bead volume of bead wash buffer [50 mM Tris-HCl, pH 7.5, 150 mM NaCl, 0.5% (v/v) NP40 alternative]. A DynaMag-2 magnet (Thermo Fisher Scientific 12321D) was used for collecting beads. Clarified lysate containing ∼3 mg protein was then added to 10 μL packed bead volume of the anti-FLAG beads for each immunoprecipitation, with the total solution volume brought up to 500 μL by adding lysis buffer. Samples were incubated with rotation for 4 h at 4°C. Following immunoprecipitation, beads were washed 3 times with 20X packed bead volume of lysis buffer containing 400 mM NaCl. Proteins were eluted from beads using 1X Laemmli sample buffer (Bio-Rad 1610747) containing 2.5% (v/v) 2-mercaptoethanol (Sigma-Aldrich M3148) and then heated at 72°C for 10 min for SDS-PAGE. For preparation of cell lysate samples for SDS-PAGE, clarified lysates were adjusted to have equal protein concentrations across samples and then mixed with 4X Laemmli sample buffer containing 10% (v/v) 2-mercaptoethanol to achieve a final 1X concentration and then heated at 72°C for 10 min.

##### Fractionation of human cells

Cellular fractionation was performed using the Cell Fractionation Kit (Cell Signaling Technology 9038) generally according to the manufacturer’s instructions. Wide bore p1000 tips were used for manipulating solutions containing cells and organelles so as to minimize damage. All steps were done on ice or at 4°C except those involving the kit’s nuclear isolation buffer, which was done at room temperature to avoid precipitation. 1/4 tablet cOmplete protease inhibitor cocktail per 5 mL of the kit’s cytosolic isolation buffer, organellar isolation buffer, and nuclear isolation buffer was used instead of the kit’s protease inhibitor cocktail.

Cellular fractions from HCT116 cells were generated by first preparing a suspension of 5 million live HCT116 cells in 500 μL ice-cold DPBS. From this, 100 μL of the cell suspension (*i.e.*, whole cells) was mixed with 4X Laemmli sample buffer containing 10% (v/v) 2-mercaptoethanol to achieve a final 1X concentration. The sample was then sonicated, heated at 95°C for 5 min, centrifuged at 15,000 g for 3 min at room temperature, and the supernatant was taken for SDS-PAGE. The remaining 400 μL of the cell suspension in DPBS was centrifuged at 500 g for 5 min at 4°C and the supernatant discarded. The pellet was resuspended in 400 μL of the cytosolic isolation buffer, vortexed for 5 sec, left on ice for 5 min, and then centrifuged at 500 g for 5 min at 4°C. 100 μL of the supernatant (*i.e.*, cytosolic fraction) was mixed with 4X Laemmli sample buffer containing 10% (v/v) 2-mercaptoethanol to achieve a final 1X concentration, then heated at 95°C for 5 min, centrifuged at 15,000 g for 3 min at room temperature, and the supernatant was taken for SDS-PAGE. The remainder of the cytosolic fraction was aspirated off and the pellet was resuspended in 400 μL of organellar isolation buffer, vortexed for 15 sec, left on ice for 5 min, and then centrifuged at 8,000 g for 5 min at 4°C. 100 μL of the supernatant (*i.e.*, organellar fraction) was mixed with 4X Laemmli sample buffer containing 10% (v/v) 2-mercaptoethanol to achieve a final 1X concentration, then heated at 95°C for 5 min, centrifuged at 15,000 g for 3 min at room temperature, and the supernatant was taken for SDS-PAGE. After aspiration of the remaining organellar fraction, the pellet was resuspended in 400 μL of nuclear isolation buffer (*i.e.*, nuclear fraction), sonicated, and mixed with 4X Laemmli sample buffer containing 10% (v/v) 2-mercaptoethanol to achieve a final 1X concentration, then heated at 95°C for 5 min, centrifuged at 15,000 g for 3 min at room temperature, and the supernatant was taken for SDS-PAGE. The same volumes - and thus the same whole-cell equivalents - of the whole-cell sample, cytosolic fraction, organellar fraction, and nuclear fraction were loaded on the SDS-PAGE gel.

CAKI-2 cells did not perform well using the aforementioned protocol for HCT116 cells (data not shown) and so modifications were made. A suspension of 3 million live CAKI-2 cells in 500 μL ice-cold DPBS was prepared. From this, 200 μL of the cell suspension (*i.e.*, whole cells) was centrifuged at 500 g for 5 min at 4°C. The supernatant was aspirated and the pellet was resuspended with lysis buffer [50 mM Tris-HCl, pH 7.5, 150 mM NaCl, 0.5% (v/v) NP40 alternative, 1/4 tablet cOmplete protease inhibitor cocktail per 5 mL lysis buffer, 1/2 tablet PhosSTOP per 5 mL lysis buffer], incubated with rotation for 30 min at 4°C, and cleared by centrifugation at 17,000 g for 10 min at 4°C. 150 μL of the clarified lysate was mixed with 4X Laemmli sample buffer containing 10% (v/v) 2-mercaptoethanol to achieve a final 1X concentration, then heated at 95°C for 5 min, centrifuged at 15,000 g for 3 min at room temperature, and the supernatant was taken for SDS-PAGE. The remaining 300 μL of the cell suspension in DPBS was centrifuged at 500 g for 5 min at 4°C and the supernatant discarded. The pellet was resuspended in 300 μL of the kit’s cytosolic isolation buffer, vortexed for 5 sec, left on ice for 5 min, and then centrifuged at 500 g for 5 min at 4°C. 150 μL of the supernatant (*i.e.*, cytosolic fraction) was mixed with 4X Laemmli sample buffer containing 10% (v/v) 2-mercaptoethanol to achieve a final 1X concentration, then heated at 95°C for 5 min, centrifuged at 15,000 g for 3 min at room temperature, and the supernatant was taken for SDS-PAGE. The remainder of the cytosolic fraction was aspirated off and the pellet was resuspended in 300 μL of lysis buffer, incubated with rotation for 30 min at 4°C, and cleared by centrifugation at 17,000 g for 10 min at 4°C. 150 μL of the clarified lysate (*i.e.*, organellar fraction) was mixed with 4X Laemmli sample buffer containing 10% (v/v) 2-mercaptoethanol to achieve a final 1X concentration, then heated at 95°C for 5 min, centrifuged at 15,000 g for 3 min at room temperature, and the supernatant was taken for SDS-PAGE. The same volumes - and thus the same whole-cell equivalents - of the whole-cell samples, cytosolic fractions, and organellar fractions were loaded on the SDS-PAGE gel.

##### Preparation of human cellular lysates

Preparation of cellular lysates from human cells was done at 4°C or on ice unless indicated. Fully confluent 10-cm dishes were washed once with ice-cold DPBS and immediately scraped into 500 μL lysis buffer [50 mM Tris-HCl, pH 7.5, 150 mM NaCl, 0.5% (v/v) NP40 alternative, one tablet cOmplete protease inhibitor cocktail per 25 mL lysis buffer, one tablet PhosSTOP per 10 mL lysis buffer]. Lysates were incubated with rotation for 30 min at 4°C before being cleared by centrifugation at 17,000 g for 10 min at 4°C. Protein quantification was performed on the clarified lysate using the *DC* protein assay. For preparation of samples for SDS-PAGE, clarified lysates were adjusted to have equal protein concentrations across samples and then mixed with 4X Laemmli sample buffer containing 10% (v/v) 2-mercaptoethanol to achieve a final 1X concentration and then heated at 72°C for 10 min.

##### Immunoblotting of samples from human cells

Samples were resolved by SDS-PAGE via 4% - 20% Mini-PROTEAN TGX gels (Bio-Rad 456-1096), transferred to 0.2 μm nitrocellulose membranes (Bio-Rad 1704158, 1704159) by using the Trans-Blot Turbo transfer system (Bio-Rad 1704150). Membranes were then blocked for 45 min at room temperature with 5% non-fat dry milk (Bio-Rad 1706404) prepared in 1X TBST (Bioworld 40120065-2) and incubated with primary antibodies in 1X TBST overnight at 4°C at the following dilutions: ACAA1 (1/250), AGPS (1/1000), C6ORF226 (1/1000), PEX5 (1/1000), PEX13 (1/1000), ACTB (1/2500), PEX7 (1/1000), PHYH (1/1000), SCP2 (1/500), CANX (1/1000), CS (1/1000), GAPDH (1/5000), HA (1/1000), Histone H3 (1/1500), and RPS6KB1 (1/1000). Afterwards, membranes were washed 3 times with 1X TBST, then incubated with HRP-conjugated secondary antibodies (1/5000) in 1X TBST for 45 min at room temperature. Membranes were then washed 3 more times with 1X TBST and subsequently visualized using enhanced chemiluminescence via Pierce ECL western blotting substrate (Thermo Fisher Scientific 32106) or, if increased sensitivity was needed, SuperSignal West Femto maximum sensitivity substrate (Thermo Fisher Scientific 34096). If membranes needed to be stripped and reprobed after enhanced chemiluminescence, membranes were washed twice with 1X TBST, incubated with Restore PLUS Western Blot Stripping Buffer (Thermo Fisher Scientific 46430) for 15 min at room temperature, washed once more with 1X TBST, and then blocked and immunoblotted as described already.

##### Quantification of immunoblots

Densitometric analyses of immunoblots were performed using the software ImageJ (version 1.54d),^87^ and background signals were corrected for. For quantification of immunoblots, band intensities were used as indicated in the respective figures and figure legends. With regards to the generation of mature/precursor ratios for PHYH, ACAA1, and AGPS in Figures 3B and S5D, the precursor forms were undetectable in all cells except for those overexpressing wild-type *Hs*PEX39. As such, to avoid dividing by zero when generating the ratios, a pseudonumbering strategy was adapted from prior work,^91^ in which all band intensities for a protein’s precursor form were adjusted by adding 0.1 x the smallest non-zero band intensity observed for that protein’s precursor form across all samples.

#### Experiments with radiolabeled proteins

##### Plasmids

The plasmids pET28a-PEX7, encoding an N-terminally histidine (His)-tagged version of human PEX7^64^; pET23a-PEX5(C11K), encoding a version of the large isoform of human PEX5 possessing a lysine at position 11^92^; pET28a-PEX5(1-324), encoding a His-tagged protein comprising residues 1 to 324 of the large isoform of PEX5^93^; pQE30-PEX5(315-639), encoding a His-tagged protein comprising residues 315 to 639 of the large isoform of PEX5^94^; pGEM4-ACAA1, which encodes the precursor form of human ACAA1^64^; pGEM4-PHYH, encoding the precursor form of human PHYH^37^ and pQE31-PHYH which encodes a His-tagged version of PHYH^95^ were all described before.

The following plasmids were produced:

1. pET28a-*Hs*PEX39 - encodes an N-terminally His-tagged version of *Hs*PEX39 and was obtained as follows: the *HsPEX39* cDNA was amplified from total genomic DNA from HepG2 cells using the primers Hs.C6ORF226_EcoRI_Fw and Hs.C6ORF226_NotI_Rv. The amplified fragment was inserted into EcoRI/NotI-digested pET28a;
2. pGEX-*Hs*PEX39 - encodes an N-terminally GST-tagged version of *Hs*PEX39. The *HsPEX39* DNA fragment described above was inserted into the EcoRI/NotI-sites of pGEX-6-P1 (GE Healthcare);
3. pGEX-*Hs*PEX39(1-70) - encodes an N-terminally GST-tagged *Hs*PEX39 containing residues 1 to 70. It was generated by introducing 2 stop codons by site-directed mutagenesis with the primers ORF70stop2x_Fw and _ ORF70stop2x_Rv;
4. pGEX-*Hs*PEX39(48-101) - encodes an N-terminally GST-tagged *Hs*PEX39 comprising residues 48 to 101. The corresponding DNA fragment was obtained by PCR with the primers Hs.C6ORF226_Glu48_EcoRI_Fw and Hs.C6ORF226_NotI_Rv and inserted into EcoRI/NotI sites of pGEX-6-P1 (GE Healthcare);
5. pGEX-*Hs*PEX39(KPWEmut) - encodes an N-terminally GST-tagged version of *Hs*PEX39 where the KPWE motif is replaced by AAAA. It was obtained by overlap-extension PCR. Two overlapping regions of *HsPEX39* cDNA, both encoding for the KPWE-to-AAAA mutation, were obtained by PCR with the primer pairs Hs.C6ORF226_EcoRI_Fw/P2_KPWE-Ala_Rv and P3_KPWE-Ala_Fw/Hs.C6ORF226_NotI_Rv. Resulting PCR products were denatured, annealed and used in a third PCR with the primers Hs.C6ORF226_EcoRI_Fw and Hs.C6ORF226_NotI_Rv. The amplified fragment was digested and inserted into the EcoRI/NotI-sites of pGEX-6-P1 (GE Healthcare);
6. pGEX-*Hs*PEX39(48-101,KPWEmut) - encodes an N-terminally GST-tagged *Hs*PEX39 protein comprising residues 48 to 101 that has the KPWE motif mutated to AAAA. It was obtained by PCR amplification of pGEX-*Hs*PEX39(KPWEmut) with primers Hs.C6ORF226_Glu48_EcoRI_Fw and Hs.C6ORF226_NotI_Rv. The resulting fragment was digested and inserted into EcoRI/NotI sites of pGEX-6-P1 (GE Healthcare);
7. pET23a-NtPEX13.Sumo.His - The cDNA encoding residues 1 to 36 of *Hs*PEX13 appended to the TEV recognition sequence SENLYFQG followed by residues 2-101 of human SUMO-1 protein with a C-terminal His-tag was synthesized and cloned into the NdeI/BamHI restriction sites of pET23a (Novagen) by GenScript.
8. pET23a-NtPEX13(KPWEmut).Sumo.His - Obtained by site directed mutagenesis of the pET23a-NtPEX13.Sumo.His construct from GenScript.

##### Expression and purification of recombinant proteins

Recombinant His-PEX5(1-324), His-PEX5(315-639) (also referred to as TPRs),^94^ and His-PHYH^95^ were expressed and purified exactly as described before.

The N-terminally His-tagged and all GST-tagged versions of *Hs*PEX39 were expressed in *E. coli* BL21(DE3) for 3 h at 37°C with 0.5 mM isopropyl β-D-1-thiogalactopyranoside (IPTG), and purified using either Ni Sepharose 6 Fast Flow or Glutathione Sepharose 4B affinity chromatography resins (GE Healthcare) according to the manufacturer’s instructions. All proteins were stored in 50 mM Tris-HCl, pH 8.0, 150 mM NaCl, 1 mM EDTA-NaOH, pH 8.0, 1 mM DTT at -80°C.

Where indicated, GST-*Hs*PEX39 variants were treated with His-tagged HRV-3C protease to remove the GST tag. Briefly, GST-tagged proteins were incubated with 3C protease [30:1 (w/w)] for 16 h at 4°C in 50 mM Tris-HCl, pH 8.0, 150 mM NaCl, 1 mM EDTA-NaOH, pH 8.0, 1 mM DTT. The His-tagged protease was removed from the preparation by affinity chromatography using Ni Sepharose 6 Fast Flow resin and collecting the flow through.

##### *In vitro* synthesis of radiolabeled proteins

Radiolabeled proteins were synthesized *in vitro* using the TNT® T7 Quick Coupled Transcription/Translation System (Promega) in the presence of EasyTag^TM^ L-[^35^S]methionine (specific activity > 1,000 Ci/mmol, Perkin Elmer) for 90 min at 30°C, according to the manufacturer’s instructions, with the exception of PEX7 which was synthesized for 4 h. Unlabeled (“cold”) PEX7 was synthesized in the same way but using unlabeled methionine provided in the kit. Aliquots of the rabbit reticulocyte lysates containing the synthesized proteins were snap-frozen in liquid nitrogen and stored at -80°C.

##### Cell-free *in vitro* assays

*In vitro* peroxisomal import reactions using HEK293-derived post-nuclear supernatants (PNS) were performed as previously described.^42^ Briefly, the rabbit reticulocyte lysates (RRL) containing the radiolabeled reporter protein (0.2-1 μL of RRL per import reaction) was diluted in 10 μL import buffer (0.25 M sucrose, 50 mM KCl, 20 mM MOPS-KOH, pH 7.2, 5 mM MgCl_2_, 2 μg/mL E-64, 48 μg/mL methionine, final concentration). For improved import yields, the diluted ^35^S-ACAA1 RRL contained also 70 nM PEX5(1-324) and 1 μL of RRL containing unlabeled PEX7 to form a PEX5-PEX7-PTS2 pre-import complex. Radiolabeled proteins were added to import reactions containing 600 μg of HEK293 PNS (previously primed by incubating at 37°C for 5 min with 0.3 mM ATP) in import buffer supplemented with 5 μM bovine ubiquitin, 2 mM GSH and either 3 mM ATP or AMP-PNP (100 μL final volume). Where specified, *in vitro* reactions were supplemented with 2 μM Ubiquitin aldehyde (HA-Ubal), TPRs (5 μM), NtPEX13.Sumo.His (up to 20 μM), His.*Hs*PEX39 or untagged *Hs*PEX39 variants (up to 1 μM). Import reactions were incubated for 30 min at 37°C. Import reactions with radiolabeled PTS2 proteins were treated with Trypsin (400 μg/mL final concentration) in order to degrade non-imported reporter protein. After a 40 min incubation on ice, proteases were inhibited with phenylmethylsulfonyl fluoride (500 μg/mL, final concentration). When monitoring radiolabeled PEX5, the protease treatment was omitted. Reactions were diluted with ice-cold SEMK (20 mM MOPS-KOH, pH 7.2, 0.25 M sucrose, 1 mM EDTA-NaOH, pH 8.0, 80 mM KCl). Organelles, and soluble fractions (if required), were collected by centrifugation (20 min, 16,000 g, 4°C). Proteins were precipitated with 10% (w/v) TCA, washed with acetone, and the resulting organelle and/or soluble fractions were analyzed by SDS-PAGE, followed by Western-blotting, Ponceau S staining, and autoradiography.

##### Native-PAGE analyses

The reticulocyte lysate containing radiolabeled His-PEX7 (0.4 μL) was incubated in the absence or presence of recombinant *Hs*PEX39 variants (4 μg), His-PEX5(1-324) (1 μg), His-PHYH (100 ng) and NtPEX13.Sumo.His variants (4 μg) as specified in figures, in 20 μL of 20 mM Tris-HCl, pH 8.0, 5 mM DTT, 0.025% (w/v) bovine serum albumin. After adding 5 μL of loading buffer [0.17% (w/v) bromophenol blue, 50% (w/v) sucrose], 6 μL of the samples were loaded onto Tris nondenaturing 8% polyacrylamide gels using a discontinuous buffer system^96^ and run at 250 V at 4°C for ∼1 h.

To determine the half-lives of protein complexes, ^35^S-His-PEX7 complexes were formed with 50 nM and 5 μM of GST-*Hs*PEX39 and NtPEX13.Sumo.His and supplemented with 5 μM of GST-HsPEX39(48-101) and 25 μM of GST-*Hs*PEX39, respectively. Aliquots were withdrawn at the indicated time points and immediately flash-frozen in liquid nitrogen. Samples were individually thawed and loaded in the gel from the longest to shortest time-points.

Gels were blotted onto nitrocellulose membranes, stained with Ponceau S, and subjected to autoradiography.

#### Bioinformatic analyses and structural predictions

##### Sequence analysis of [R/K]PWE-motif sequences

Sequences with significant relationship to human C6ORF226 were initially established by BLAST^97^ searches in the Uniprot database, using p < 0.001 as a significant threshold. The sequence set was further expanded by three rounds of iterative refinement using the generalized profile method^98^ and p < 0.001 as a threshold for including protein hits in the next iteration. Protein alignment were calculated using the L-Ins-I algorithm of the MAFFT package, v7.490.^99^ In the 2^nd^ and 3^rd^ iteration cycle, significant hits in members of the PEX13 family were picked up, based on conservation of the KPWE motif. Subsequently, the PEX13 family was analyzed by a separate set of generalized profiles, as described for the C6ORF226 family. The final hit lists after three rounds of iterative refinement were used for compiling Figure S1A.

The conserved KPWE core region and five flanking residues on either side were extracted from the final C6ORF226 and PEX13 family alignments and used for sequence logo generation using WebLogo v2.8.2 (https://weblogo.berkeley.edu).^100,101^ The final hit lists and alignment files for the C6ORF226 and PEX13 families will be provided by the authors upon request.

##### AlphaFold structural prediction of protein complexes

For the structural predictions of protein complexes, the amino acid sequences of the proteins were downloaded from Uniprot (https://www.uniprot.org/) and indicated parts were used for web-based predictions (https://colab.research.google.com/github/sokrypton/ColabFold/blob/main/AlphaFold2.ipynb) as described previously.^45,102^ The highest-ranking model of each prediction was visualized using ChimeraX (version 1.7.1.0).^103^ Parameters and output files of the predictions will be provided by the authors upon request.

### QUANTIFICATION AND STATISTICAL ANALYSIS

Statistics were calculated using GraphPad Prism 10 and Microsoft Excel unless specified otherwise. Definition of center and dispersion/precision measures, and statistical tests are described in the figure legends. A significance threshold (alpha) of 0.05 was used for all statistical tests for figures 1C, 2C, 2E, 2G, 3B, S2D, S3A, S5D, S5E. All experiments had at least two biological replicates. N refers to the number of biological replicates for all instances where n replicates are reported.

## SUPPLEMENTAL INFORMATION

Document S1. Figures S1-S7 with legends, and references Table S1. Oligonucleotide sequences Table S2. Proteomic analysis of Pex18 complexes, related to Figure 1C Table S3. Proteomic analysis of *Scpex39Δ* yeast, related to Figure 2E

## Key resources table

| REAGENT or RESOURCE | SOURCE | IDENTIFIER |
| --- | --- | --- |
| Antibodies |  |  |
| Rabbit polyclonal anti-ACAA1 | Sigma-Aldrich | Cat#HPA006764;<br>RRID:AB_1078085 |
| Rabbit polyclonal anti-AGPS | Sigma-Aldrich | Cat#HPA030211;<br>RRID:AB_10602195 |
| Rabbit polyclonal anti-C6ORF226 | Sigma-Aldrich | Cat#HPA045350;<br>RRID:AB_10964640 |
| Rabbit polyclonal anti-PEX5 | Sigma-Aldrich | Cat#HPA039260;<br>RRID:AB_2676417 |
| Rabbit polyclonal anti-HsPEX13 | Sigma-Aldrich | Cat#HPA032142;<br>RRID:AB_10669676 |
| Mouse monoclonal anti-ACTB | Santa Cruz<br>Biotechnology | Cat#sc-69879;<br>RRID:AB_1119529 |
| Rabbit polyclonal anti-PEX7 | Proteintech | Cat#20614-1-AP;<br>RRID:AB_10694826 |
| Rabbit polyclonal anti-PHYH | Proteintech | Cat#12858-1-AP;<br>RRID:AB_2163745 |
| Rabbit polyclonal anti-SCP2 | Proteintech | Cat#23006-1-AP;<br>RRID:AB_2879197 |
| Rabbit polyclonal anti-CANX | Cell Signaling<br>Technology | Cat#2433;<br>RRID:AB_2243887 |
| Rabbit monoclonal anti-CS | Cell Signaling<br>Technology | Cat#14309;<br>RRID:AB_2665545 |
| Rabbit monoclonal anti-GAPDH | Cell Signaling<br>Technology | Cat#2118;<br>RRID:AB_561053 |
| Rabbit monoclonal anti-HA | Cell Signaling<br>Technology | Cat#3724;<br>RRID:AB_1549585 |
| Mouse monoclonal anti-Histone H3 | Cell Signaling<br>Technology | Cat#3638;<br>RRID:AB_1642229 |
| Rabbit monoclonal anti-RPS6KB1 | Cell Signaling<br>Technology | Cat#2708;<br>RRID:AB_390722 |
| Goat HRP-conjugated polyclonal anti-rabbit | Cell Signaling<br>Technology | Cat#7074;<br>RRID:AB_2099233 |
| Rat HRP-conjugated monoclonal anti-mouse | Abcam | Cat#ab131368;<br>RRID:AB_2895114 |
| Goat polyclonal anti-CTA1 (*S. cerevisiae*) | Prior study<sup>1</sup> | N/A |
| Rabbit polyclonal anti-PEX7 (*S. cerevisiae*) | Prior study<sup>2</sup> | N/A |
| Rabbit polyclonal anti-PEX3 (*S. cerevisiae*) | Erdmann group RUB<br>Bochum | N/A |
| Rabbit polyclonal anti-PEX18 (*S. cerevisiae*) | Prior study<sup>3</sup> | N/A |
| Rabbit polyclonal anti-POT1 (*S. cerevisiae*) | Prior study<sup>4</sup> | N/A |
| Rabbit polyclonal anti-GPD1 (*S. cerevisiae*) | Prior study<sup>5</sup> | N/A |
| Rabbit polyclonal anti-Protein A (*S. aureus*) | Sigma-Aldrich | Cat#P3775 |
| Mouse monoclonal anti-POR1 (*S. cerevisiae*) | Invitrogen | Cat#459500 RRIP:<br>AB_2532239 |
| Mouse monoclonal anti-PGK1 (*S. cerevisiae*) | Invitrogen | Cat#459250 RRIP:<br>AB_2532235 |
| Mouse monoclonal anti-DPM1 (*S. cerevisiae*) | Invitrogen | Cat#A-6429 RRIP:<br>AB_2536204 |
| Goat HRP-conjugated polyclonal anti-rabbit | Sigma-Aldrich | Cat#A0545 |
| Rabbit HRP-conjugated polyclonal anti-goat | Sigma-Aldrich | Cat#A8919 |
| Rabbit HRP-conjugated polyclonal anti-mouse | Sigma-Aldrich | Cat#A9044 |
| Anti-FLAG M2 magnetic beads | Millipore | Cat#M8823 |
| Bacterial and virus strains |  |  |
| *E. coli* BL21-CodonPlus(DE3)-RIL | Agilent | Cat#AGLS230245 |
| *E. coli* One Shot TOP10 | Invitrogen | Cat#C404003 |
| Biological samples |  |  |
| Chemicals, peptides, and recombinant proteins |  |  |
| Ready Blue Protein Gel Stain | Merck/Sigma-Aldrich | Cat#RSB-1L |
| Quick Start Bradford 1x Dye Reagent | Biorad | Cat#5000205 |
| Lyticase from *Arthrobacter luteus* | Merck/Sigma-Aldrich | Cat#L4025 |
| Digitonin, High Purity | Merck/Millipore | Cat#300410 |
| ROTI Garose-His/Ni beads | Carl Roth | Cat#1308.1 |
| AcTEV protease | Invitrogen | Cat#12575-023 |
| <sup>13</sup>C<sub>6</sub>/<sup>15</sup>N<sub>4</sub>-L-Arginine (Arg10) | Eurisotop | Cat#CNLM-539-H |
| <sup>13</sup>C<sub>6</sub>/<sup>15</sup>N<sub>2</sub>-L-Lysine (Lys8) | Eurisotop | Cat#CNLM-291-H |
| LysC | Fujifilm/Wako Chemicals | Cat#125-05061 |
| Trypsin Sequencing Grade Modified | Promega | Cat#V5111 |
| Oleic acid | Sigma-Aldrich | Cat#27728-1L-R |
| ATP disodium salt hydrate | Sigma-Aldrich | Cat#A2383 |
| AMP-PNP tetralithium salt | Jena Bioscience | Cat#NU-407 |
| Critical commercial assays |  |  |
| Cell Fractionation Kit | Cell Signaling Technology | Cat#9038 |
| Platinum SuperFi II DNA Polymerase Kit | Invitrogen | Cat#12361010 |
| Deposited data |  |  |
| Quantitative proteomics datasets (MS/MS raw files and MaxQuant result files) | This study | ProteomeXchange: PXD051501, PXD051550 |
| Experimental models: Cell lines |  |  |
| Human: CAKI-2 cells | William Kaelin Jr. | RRID:CVCL_0235 |
| Human: HCT116 cells | ATCC | Cat#CCL-247; RRID:CVCL_0291 |
| Human: HEK293T cells | ATCC | Cat#CRL-3216; RRID:CVCL_0063 |
| Human: NCI-H1792 cells | ATCC | Cat#CRL-5895; RRID:CVCL_1495 |
| Human: HEK293 cells | Hans R. Waterham | RRID:CVCL_0045 |
| Experimental models: yeast strains |  |  |
| BY4741, wild-type strain (MATa *his3Δ1 leu2Δ0 met15Δ0 ura3Δ0*) | Euroscarf | Acc. No. Y00000 |
| BY4741 ΔScpex13 (MATa *his3Δ1 leu2Δ0 met15Δ0 ura3Δ0 Scpex13::kanMX4*) | Euroscarf | Acc. No. Y04140 |
| BY4741 Δlys2Δarg4 (MATa *his3Δ1 leu2Δ0 lys2Δ0 ura3Δ0 met15Δ0 arg4::kanMX4*) | Prior study<sup>6</sup> | SC174 (Intra-lab identifier for Warscheid group) |
| BY4741 Δlys2Δarg4ΔScpex39 (MATa *his3Δ1 leu2Δ0 lys2Δ0 ura3Δ0 met15Δ0 arg4::kanMX4 YJR012C::URA3*) | This study | SC601 (Intra-lab identifier for Warscheid group) |
| UTL7-A Δpex7 Pex18-TPA<br>(MATa *ura3-52 leu2-3/112 trp1 pex7::LEU2 PEX18:TPA-kanMX4*) | Prior study<sup>3</sup> | #2806 (Intra-lab identifier for Warscheid group) |
| UTL7-A Δfox3 Pex18-TPA<br>(MATa *ura3-52 leu2-3/112 trp1 fox3::loxP PEX18:TPA-kanMX4*) | Prior study<sup>3</sup> | #2821 (Intra-lab identifier for Warscheid group) |
| CB80 Δlys1Δarg4<br>(MATa *ura3-52 leu2-1 trp1-63 his3-200 arg4::loxP lys1::loxP*) | Prior study<sup>7</sup> | SC38 (Intra-lab identifier for Warscheid group) |
| CB80 Δlys1Δarg4 Pex18-TPA<br>(MATa *ura3-52 leu2-1 trp1-63 his3-200 arg4::loxP lys1::loxP PEX18:TPA-kanMX4*) | This study | SC377 (Intra-lab identifier for Warscheid group) |
| BY4741 Pex3-mScarlet<br>(MATa *his3Δ1 leu2Δ0 met15Δ0 ura3Δ0 Pex3-mScarlet::G418*) | This study | N/A |
| BY4741 Pex3-mScarlet Pex39-mNG<br>(MATa *his3Δ1 leu2Δ0 met15Δ0 ura3Δ0 Pex3-mScarlet::G418 Pex39-mNG::URA*) | This study | N/A |
| BY4741 Pex3-mScarlet Pot1-mNG<br>(MATa *his3Δ1 leu2Δ0 met15Δ0 ura3Δ0 Pex3-mScarlet::G418 Pot1-mNG::URA*) | This study | N/A |
| BY4741 ΔScpex39 Pex3-mScarlet Pot1-mNG<br>(MATa *his3Δ1 leu2Δ0 met15Δ0 ura3Δ0 Pex3-mScarlet::G418 ΔScpex39::NAT Pot1-mNG::URA*) | This study | N/A |
| BY4741 Δpex7 Pex3-mScarlet Pot1-mNG<br>(MATa *his3Δ1 leu2Δ0 met15Δ0 ura3Δ0 Pex3-mScarlet::G418 Δpex7::NAT Pot1-mNG::URA*) | This study | N/A |
| Oligonucleotides |  |  |
| See Table S1 for a list of oligonucleotides used in this study | This study | N/A |
| Recombinant DNA |  |  |
| Plasmid: pHAGE-ACAA1-FLAG-HA | This study | N/A |
| Plasmid: pHAGE-PHYH-FLAG-HA | This study | N/A |
| Plasmid: pHAGE-PEX7-FLAG-HA | This study | N/A |
| Plasmid: pHAGE-HsPEX39-FLAG-HA | This study | N/A |
| Plasmid: pHAGE-GAPDH-FLAG-HA | This study | N/A |
| Plasmid: pHAGE-HsPEX39(KPWE mutants)-FLAG-HA | This study | N/A |
| Plasmid: pHAGE-FLAG-HA-EGFP | This study | N/A |
| Plasmid: pHAGE-GAPDH | This study | N/A |
| Plasmid: pHAGE-HsPEX39 | This study | N/A |
| Plasmid: pHAGE-HsPEX39(KPWE mutant) | This study | N/A |
| Plasmid: TLCV2 | Prior study,<sup>8</sup> Addgene | Addgene Plasmid#87360 |
| Plasmid: TLCV2-sgAAVS1 | This study | N/A |
| Plasmid: TLCV2-sgHsPEX39_1 | This study | N/A |
| Plasmid: TLCV2-sgHsPEX39_2 | This study | N/A |
| Plasmid: TLCV2-sgHsPEX39_3 | This study | N/A |
| Plasmid: pRS313 PEX39Pro-ScPex39-PEX39Term | This study | P797 |
| Plasmid: pRS313 TEF2Pro-ScPex39-ADH1Term | This study | P798 |
| Plasmid: pRS313 PEX13Pro-ScPex13-PEX13Term | This study | P801 |
| Plasmid: pRS313 PEX13Pro-ScPex13(KPWE/AAAA mutant)-PEX13Term | This study | P807 |
| Plasmid: pRS313 PEX39Pro-ScPex39(RPWE/AAAA mutant)-PEX39Term | This study | P815 |
| Plasmid: pET28a-PEX7 | Prior study<sup>9</sup> | N/A |
| Plasmid: pET23-PEX5(C11K) | Prior study<sup>10</sup> | N/A |
| Plasmid: pGEM4-ACAA1 | Prior study<sup>9</sup> | N/A |
| Plasmid: pGEM4-PHYH | Prior study<sup>11</sup> | N/A |
| Plasmid: pQE31-PHYH | Prior study<sup>12</sup> | N/A |
| Plasmid: pET28a-PEX5(1-324) | Prior study<sup>13</sup> | N/A |
| Plasmid: pGEX-HsPEX39 | This study | N/A |
| Plasmid: pET28a-HsPEX39 | This study | N/A |
| Plasmid: pGEX-HsPEX39(1-70) | This study | N/A |
| Plasmid: pGEX-HsPEX39(48-101) | This study | N/A |
| Plasmid: pGEX-HsPEX39(KPWEmut) | This study | N/A |
| Plasmid: pGEX-HsPEX39(48-101,KPWEmut) | This study | N/A |
| Plasmid: pET23a-NtPEX13.Sumo.His | This study | N/A |
| Plasmid: pET23a-NtPEX13(KPWEmut).Sumo.His | This study | N/A |
| Plasmid: pQE30-PEX5(315-639) | Prior study<sup>14</sup> | N/A |
| Plasmid: pFA6a-NAT-MX6 (knockout-mediated plasmid with Nat cassette with TEF2 promoter and terminator) | This study | pMS49 (Intra-lab identifier from Zalckvar group) |
| Plasmid: mNG C-terminal tagging plasmid (plasmid for C-terminal tagging with mNeonGreen) | This study | pMS1190 (Intra-lab identifier from Zalckvar group) |
| Software and algorithms |  |  |
| MaxQuant/Andromeda, versions 2.0.1.0 and 2.4.4.0 | Prior studies<sup>15,16</sup> | https://www.maxquant.org/ |
| ImageJ (v 1.54d) | Prior study<sup>17</sup> | N/A |
| ColabFold (v1.5.5) | Prior study<sup>18</sup> | https://colab.research.google.com/github/sokrypton/ColabFold/blob/main/AlphaFold2.ipynb#scrollTo=UGUBLzB3C6WN |
| ChimeraX (v 1.7.1.0) | Prior study<sup>19</sup> | https://www.rbvi.ucsf.edu/chimerax |
| IUPred2A | Prior study<sup>20</sup> | https://iupred2a.elte.hu/ |
| Phyre2 | Prior study<sup>21</sup> | http://www.sbg.bio.ic.ac.uk/~phyre2/html/page.cgi?id=index |
| Other |  |  |
| C18-SD 7 mm/3 mL extraction disc cartridges | 3M Empore | Cat#4215SD |
| C18 extraction disks (StageTips) | 3M Empore | Cat#2215 |
| Glass beads, unwashed (0.425 - 0.6 mm) | Sigma-Aldrich | Cat#G9268 |
| 0.2 µm PTFE syringe filter | Phenomenex | Part#AF0-3202-12 |
| Gemini 3 µm NX C18 column | Phenomenex | Part#00F-4453-B0 |
| SuperSignal™ West Pico PLUS Chemiluminescent Substrate | Thermo Scientific | Cat#34580 |
| SuperSignal™ West Femto Substrate | Thermo Scientific | Cat#34096 |
| Mobicol spin columns with filter (35 µm pore size) | MoBiTec | Cat#M1003 and # M513515 |
| NuPAGE BisTris gradient gels (4% - 12%) | Invitrogen/Thermo Fisher Scientific | Cat#NP0321 |
| EasyTag™ L-[<sup>35</sup>S]methionine | Perkin Elmer | Cat#NEG709A |
| TnT® Quick Coupled Transcription/Translation System | Promega | Cat#L1170 |

