## Supplemental Figures S1-S7 for "PEX39 facilitates the peroxisomal import of PTS2 proteins"

### SUPPLEMENTAL FIGURE LEGENDS

**Figure S1. PEX39 and [R/K]PWE motif conservation, Pex18 enrichment, and PEX39 subcellular localization, related to Figure 1**

(A) Examination of [R/K]PWE motifs and Yjr012c/C6ORF226 orthologs across eukaryotes. Left: Sequence logos for the regions containing the conserved [R/K]PWE motifs in Yjr012c/C6ORF226 orthologs and PEX13 were generated by WebLogo. The height of the stack indicates the sequence conservation at that position, while the height of symbols within the stack indicates the relative frequency of each amino acid at that position. Right: Domain architecture of Yjr012c/C6ORF226 orthologs, PEX13, and PEX14 across major eukaryotic taxa. Colored boxes indicate sequence regions typical for PEX14 (green), Yjr012c/C6ORF226 orthologs (purple), and PEX13 (orange). Red boxes indicate the position of the [R/K]PWE motif. Grey boxes indicate additional domains only found in some taxa. In most fungi, invertebrates and fishes, Yjr012c/C6ORF226 orthologs possess an additional N-terminal region of unknown function (indicated with "X"), which should not be confused with DUF5572. SAR, *Stramenopiles-Alveolata-Rhizaria*.

(B) Pex18 enrichment via affinity purification. Shown are representative immunoblots of eluates (E) and bead (B) samples of the soluble fractions of wild-type (WT) cells and cells expressing Pex18-TPA with the corresponding Coomassie-stained membranes. TEV-protease was added to beads to cleave the TPA tag and elute Pex18 complexes.

(C) An *Sc*Pex39-mNeonGreen fusion protein localizes to the cytosol and peroxisomes in yeast per fluorescent microscopy. The localization of *Sc*Pex39-mNeonGreen (C-terminal tagging) was examined after 4 h of growth in media with oleic acid as the primary energy source, a condition that stimulates peroxisomal biogenesis [S1]. Peroxisomes were visualized with Pex3-mScarlet. Scale bar: 5 $\mu$m.

(D) *Hs*PEX39 is a cytosolic protein. Whole-cell and the indicated cellular fractions were prepared from HCT116 cells and analyzed by immunoblotting for the indicated proteins. In parentheses are the subcellular compartments that GAPDH, SCP2, and Histone H3 are markers for. SCP2 represents the ~15 kD protein that results from transcription of a downstream promoter for the *SCP2* gene and has been used previously as a marker of peroxisomes [S2, S3].

**Figure S2. Characterization of PEX39 loss in yeast and human cells, related to Figure 2**

(A) *Scpex39Δ* cells grow like wild-type yeast on glucose. Experiment performed as described in Figure 2A except that cells precultured in 0.3% glucose were shifted to medium containing 2% glucose instead of oleic acid. Data are mean ± standard deviation (SD) (n = 3). Error bars may not be visible if the SD is very small.
(B) Immunoblots of the two additional biological replicates used for quantification of the cellular fractionation data of *Scpex39Δ* yeast. Experiments performed as described in Figure 2B, quantification of the data is shown in Figure 2C. PNS, post-nuclear supernatant; S, cytosolic supernatant; OP, organellar pellet.
(C) Assessment of Pex7 levels in yeast cells exhibiting different manipulations of *Sc*Pex39. Post-nuclear supernatants generated from the indicated yeast strains were analyzed by immunoblotting using a Pex7-specific antibody. The corresponding Coomassie-stained membrane confirms equal loading of the samples. L, protein ladder.
(D) Quantification of mature ACAA1, mature AGPS, PEX7, and PEX5 in *HsPEX39*-knockout cells. Band intensities of immunoblots prepared per Figure 2F were quantified using ImageJ. Loading-control intensities were used to normalize protein abundance across samples. The names of quantified proteins are shown on the bottom. Precursor forms of ACAA1 and AGPS were undetectable and thus not quantified. Data are mean ± standard error of the mean (SEM) (n = 3), and P values were calculated using unpaired, two-tailed t-tests.
(E) Precursor and mature forms of PHYH increase and decrease, respectively, in human cells upon depletion of *Hs*PEX39. Cells were generated using lentiviral transduction of doxycycline-inducible CRISPR-Cas9 and multiple independent sgRNAs against *HsPEX39* (sg*HsPEX39*_1-3) or a negative control sgRNA against the *AAVS1* gene (sg*AAVS1*) in the CAKI-2 and NCI-H1792 cell lines. Cellular lysates were analyzed by immunoblotting for the indicated proteins after cells were treated with doxycycline. For the PHYH, ACAA1, and AGPS blots, the solid and open red arrowheads indicate the mature and precursor forms of these proteins, respectively. Precursor forms of ACAA1 and AGPS were undetectable. CANX is a loading control. Short and long exposures are denoted s.e. and l.e., respectively.
(F) Depletion of *Hs*PEX39 impairs the import of PHYH per cellular fractionation. Whole-cell (Cell), cytosolic (Cyto), and organellar (Org) fractions were prepared from the CAKI-2 cells expressing sg*AAVS1* (negative control) or sg*HsPEX39*_1 that are described in (E). For the PHYH blot, the solid and open red arrowheads indicate the mature and precursor forms, respectively. The cellular fractions that RPS6KB1 and CANX are markers for are indicated in parentheses.

**Figure S3. Investigation of the effects of excess PEX39 in human and yeast cells, related to Figure 3**

(A) Quantification of PEX7 and PEX5 upon *Hs*PEX39 overexpression. Band intensities of immunoblots prepared per Figure 3A were quantified using ImageJ. Data are mean ± SEM (n = 3), and P values were calculated using unpaired, two-tailed t-tests.

(B) *Hs*PEX39 does not inhibit PTS1-protein import *in vitro*. $^{35}$S-PEX5(C11K) was subjected to import assays at 37ºC in the presence or absence of $H_6$*Hs*PEX39 and TPRs, as indicated. PEX5(C11K) is used to improve detection of ubiquitination under reducing conditions. After incubation, organellar pellet (OP) and cytosolic supernatant (S) fractions were isolated by centrifugation and analyzed by SDS-PAGE and autoradiography. “PEX5” and “Ub-PEX5” indicate the migration of non-modified and monoubiquitinated PEX5(C11K), respectively. The presence of monoubiquitinated PEX5(C11K) in the supernatant reflects successful PEX5 processing by peroxisomes and PTS1-protein import. TPR is a PEX5 variant that inhibits PEX5 association with peroxisomes by sequestering PTS1 proteins, as described previously [S4]; as expected, upon addition of TPR, ubiquitinated PEX5 was no longer found in the supernatant fractions (reactions 2 vs 1 and 4 vs 3). “I” is 5% of the reticulocyte lysate containing the $^{35}$S-labeled protein used in each reaction. Numbers along the left side of each image indicate molecular weights (kD). See STAR Methods for detailed description of this assay.

(C) *HsPEX39* gene expression across cell lines of the Cancer Cell Line Encyclopedia. Shown are RNA-seq data (TPM) from the DepMap Portal (https://depmap.org/portal/) for 369 cell lines for which proteomics was performed and *Hs*PEX39 protein was undetectable [S5]. The dashed line indicates the cutoff (TPM = 1) for the *HsPEX39* gene to be considered expressed and grey circles represent cell lines that do not meet that cutoff.

(D) Yeast overexpressing *Sc*Pex39 grow like wild-type yeast on glucose. Experiment performed as described in Figure S2A. Data for wild-type, *Scpex39Δ* and *Scpex39Δ* + pPEX39 are the same as shown in Figure S2A. Data are mean ± SD (n = 3). Error bars may not be visible if the SD is very small.

(E) Immunoblots of two additional biological replicates of the cellular fractionation of yeast overexpressing *Sc*Pex39. Experiments performed as described for Figure 3E. PNS, post-nuclear supernatant; S, cytosolic supernatant; OP, organellar pellet.

**Figure S4. Confidence measures of predicted AlphaFold modelling, related to Figure 4**

(A) Confidence measures for human protein complex predictions. Predicted aligned error plots and predicted LDDT (local distance difference test) plots of human AlphaFold models. To predict human protein complexes, PEX39 (full-length), PEX7 (full-length), PTS2 sequence of PHYH (amino acids 1-24) and the PEX7 binding domain of the long isoform of PEX5 (amino acids 179-266) were used.

(B) Same plots as described in (A) for predictions of yeast protein complexes. Protein complexes from yeast were predicted using the amino acid sequence of *Sc*Pex39 (full-length), Pex7 (full-length), the PTS2 sequence of Pot1 (amino acids 1-30) and the Pex7 binding domain of Pex18 (amino acids 192-283).

**Figure S5. Characterization of [R/K]PWE motif of yeast and human PEX39, related to Figure 5**

(A) *Scpex39Δ* cells expressing *Sc*Pex39 with an RPWE-to-AAAA mutation grow like wild-type yeast on glucose. Experiment performed as described in Figure S2A. Data are mean ± SD (n = 3). Error bars may not be visible if the SD is very small. Data for wild-type, *Scpex39Δ*, and *Scpex39Δ* + pPEX39 are the same as shown in Figure S2A.

(B) Immunoblots of two additional biological replicates for cellular fractionation of *Scpex39Δ* cells expressing the *Sc*Pex39 RPWE-to-AAAA mutant. Experiments were performed as described for Figure 5D. PNS, post-nuclear supernatant; S, cytosolic supernatant; OP, organellar pellet.

(C) Alignment of PEX7 orthologs for determination of relative sequence conservation. Multiple sequence alignment of PEX7 orthologs from the indicated species via ClustalΩ (v.1.2.4) was used for coloring of PEX7 sequence conservation in the surface representation of AlphaFold model.

(D) Quantification of mature and precursor forms of PHYH, ACAA1, and AGPS upon overexpression of *Hs*PEX39 with a mutated KPWE motif. Band intensities of immunoblots prepared per Figure 5G were quantified using ImageJ. The data for cells overexpressing GAPDH and wild-type *Hs*PEX39 are the same as shown previously in Figure 3B and are used here as comparisons for overexpression of *Hs*PEX39 with mutation of the KPWE motif to AAAA [*Hs*PEX39(4A)]. The 3 overexpression lines were processed together in the same experiments to generate these data. Data are mean ± SEM (n = 3), and multiplicity-adjusted P values were calculated using ordinary, one-way ANOVA with post-hoc Tukey's test.

(E) Quantification of PEX7 and PEX5 upon overexpression of *Hs*PEX39 with a mutated KPWE motif. Legend information same as (D) except the data for cells overexpressing GAPDH and wild-type *Hs*PEX39 are the same as shown previously in Figure S3A.

**Figure S6. Characterization of the KPWE Motif in the N-terminus of PEX13, related to Figure 6**

(A) ScanProsite analysis of generic [R/K]-P-W-[E/Q] motifs in yeast and humans. Using the webtool ScanProsite (https://prosite.expasy.org/scanprosite/), the UniProtKB sequence database of human or yeast were searched for proteins harboring the short linear motif [R/K]-P-W-[E/Q]. Hits were then compared with proteins listed as yeast and human PEX7 interactors in the BioGRID database.

(B) Confidence measures for modeling the PEX7-PEX13 interaction for human and yeast orthologs. Predicted aligned error plots (top) and predicted LDDT (local distance difference test) plots (bottom) of human (left) and yeast (right) AlphaFold models. To predict human protein complexes, PEX7 (full-length) and the N-terminus of PEX13 (amino acids 1-55) were used. Protein complexes from yeast were predicted using the amino acid sequence of Pex7 (full-length) and the N-terminus of Pex13 (amino acids 1-55).

(C) NtPEX13 inhibits peroxisomal import of ACAA1 *in vitro*. $^{35}$S-ACAA1 was subjected to *in vitro* import assays at 37ºC in the presence of increasing concentrations of NtPEX13 or NtPEX13(4A) as indicated. After incubation, reactions were treated with trypsin and organelles were isolated by centrifugation and analyzed by SDS-PAGE and autoradiography. Precursor and mature forms of ACAA1 denoted by open and solid red arrowheads. I, 5% of the reticulocyte lysate containing the $^{35}$S-labeled protein used in each reaction.

(D) NtPEX13 does not inhibit PTS1-protein import *in vitro*. $^{35}$S-PEX5(C11K) was subjected to *in vitro* import assays at 37ºC in the presence or absence of NtPEX13 and in the presence of either ATP or AMP-PNP, as indicated. After incubation, organelle pellet (OP) and cytosolic supernatant (S) fractions were isolated by centrifugation and analyzed by SDS-PAGE and autoradiography. "PEX5" and "Ub-PEX5" indicate the migration of non-modified PEX5 and monoubiquitinated PEX5, respectively. AMP-PNP inhibits the ATPases PEX1/PEX6 and thereby inhibits the transfer of monoubiquitinated PEX5 from peroxisomes into the cytosolic supernatant [S6]. I, 5% of the reticulocyte lysate containing the $^{35}$S-labeled protein used in each reaction.

(E) Immunoblots of two additional biological replicates for cellular fractionation of *Scpex13*Δ cells expressing the *Sc*Pex13 KPWE-to-AAAA mutant. Experiments performed as described for Figure 6F. PNS, post-nuclear supernatant; S, cytosolic supernatant; OP, organellar pellet.

**Figure S7. Comparison of structural predictions of yeast and human PEX7, related to Figure 7**

(A) Superimposition of AlphaFold structural models of human and yeast PEX7. Predicted models were directly received from the AlphaFold database (https://alphafold.ebi.ac.uk/) using accession codes AF-A0A7Y7DV00-F1 (*Hs*PEX7) and AF-A0A7L0J873-F1 (*Sc*Pex7) and superimposed using ChimeraX. AlphaFold confidence scores (*i.e.*, strength of structural predictions) are shown according to the predicted local distance difference test (pLDDT) (right). Extended loops of *Sc*Pex7 are highlighted in red and respective amino acids are labelled (bottom).

FIGURE S1, related to FIGURE 1

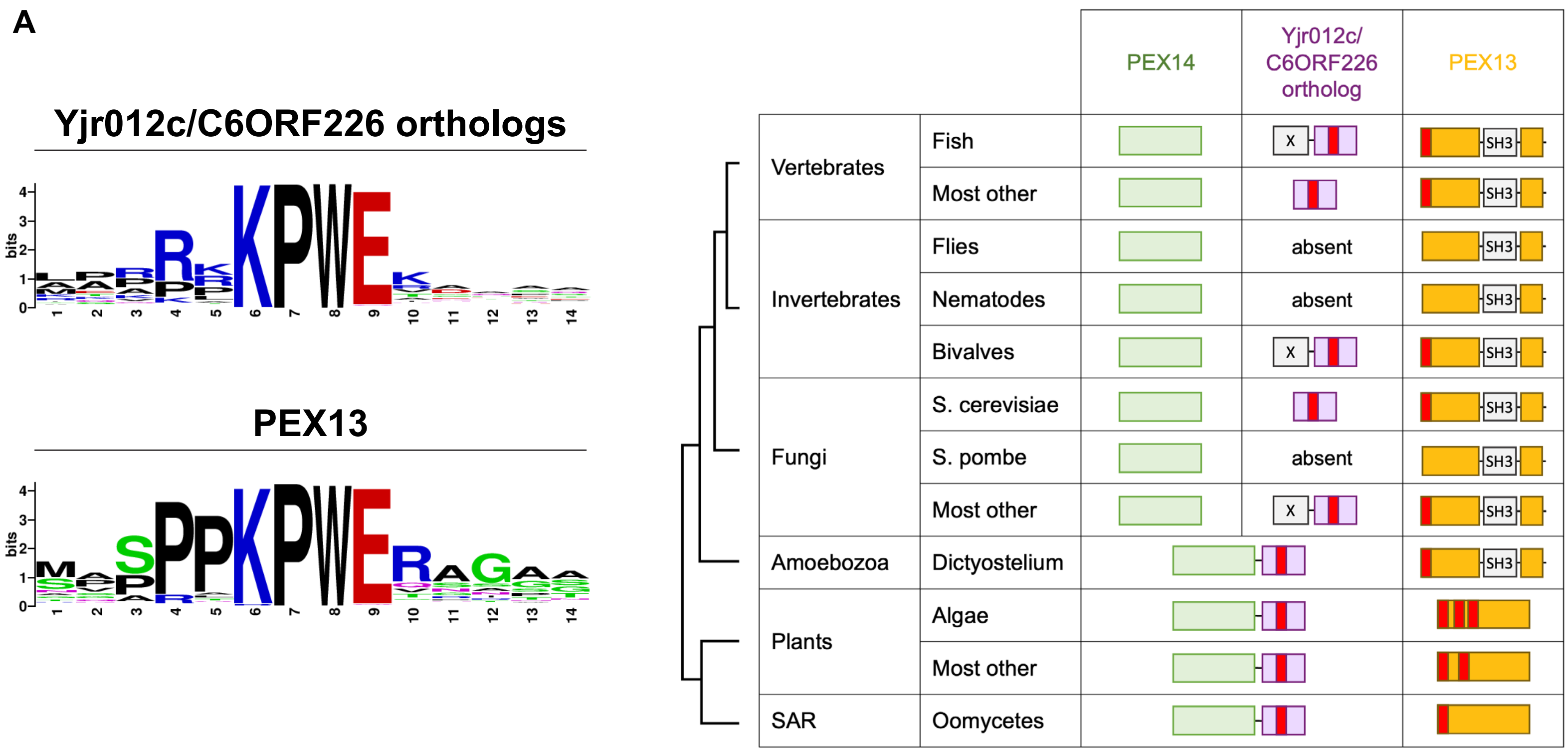


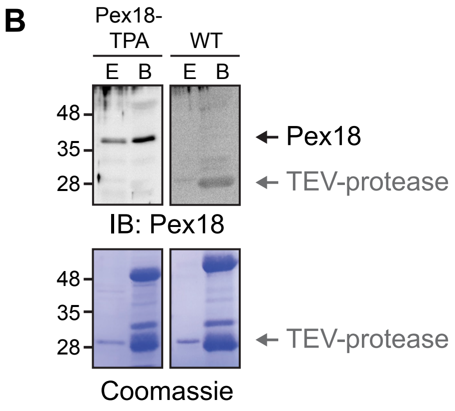


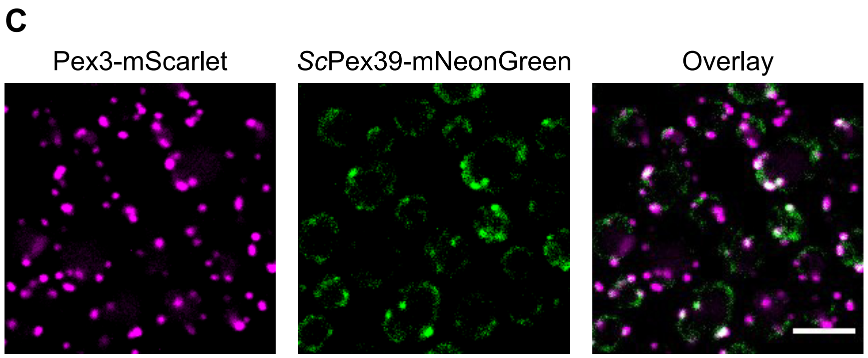


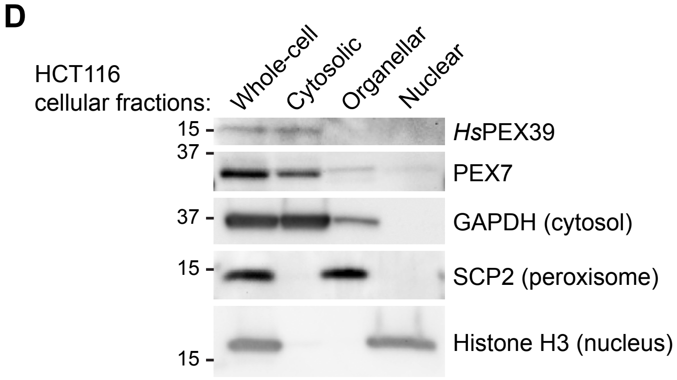

**FIGURE S2, related to FIGURE 2**

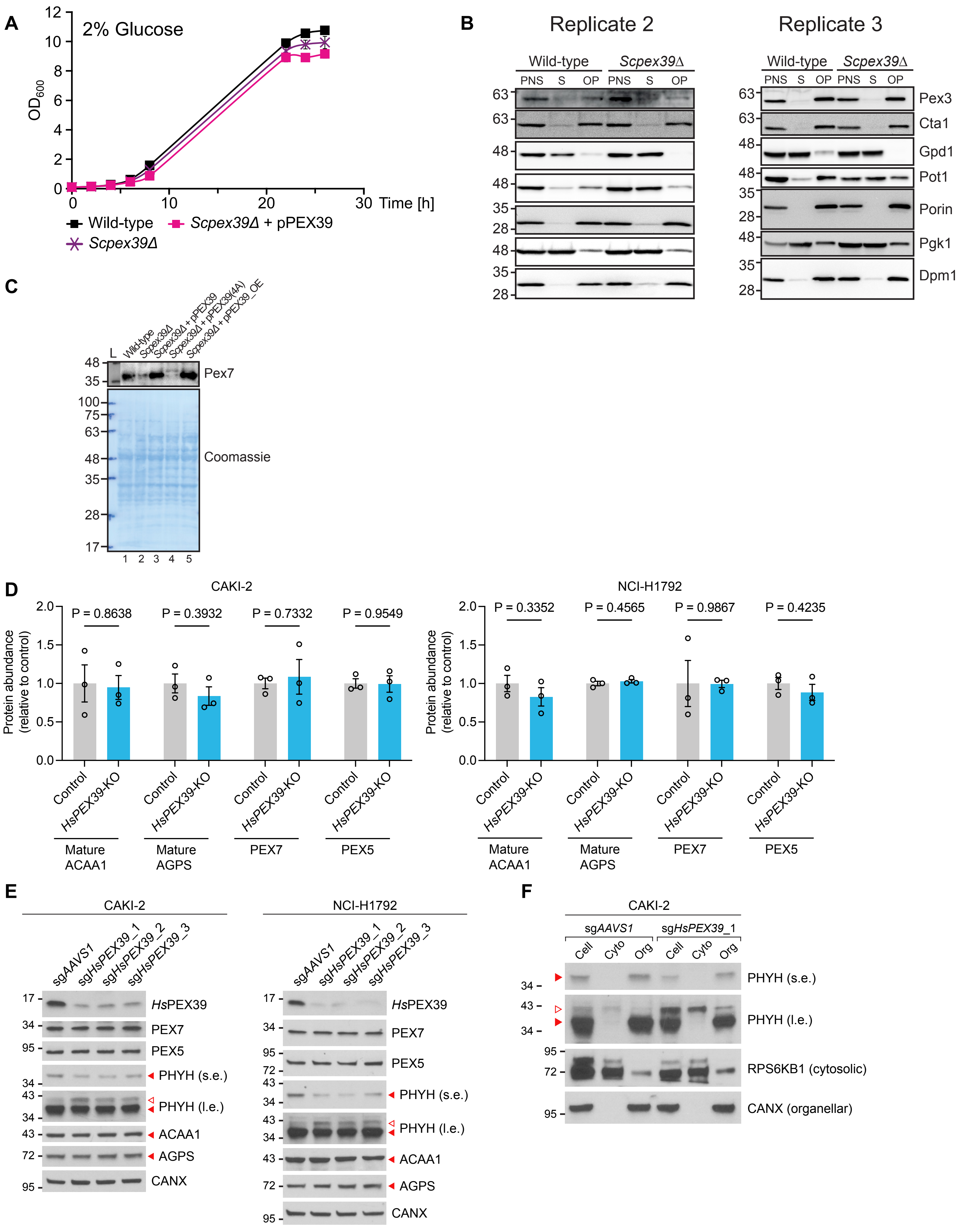

**FIGURE S3, related to FIGURE 3**

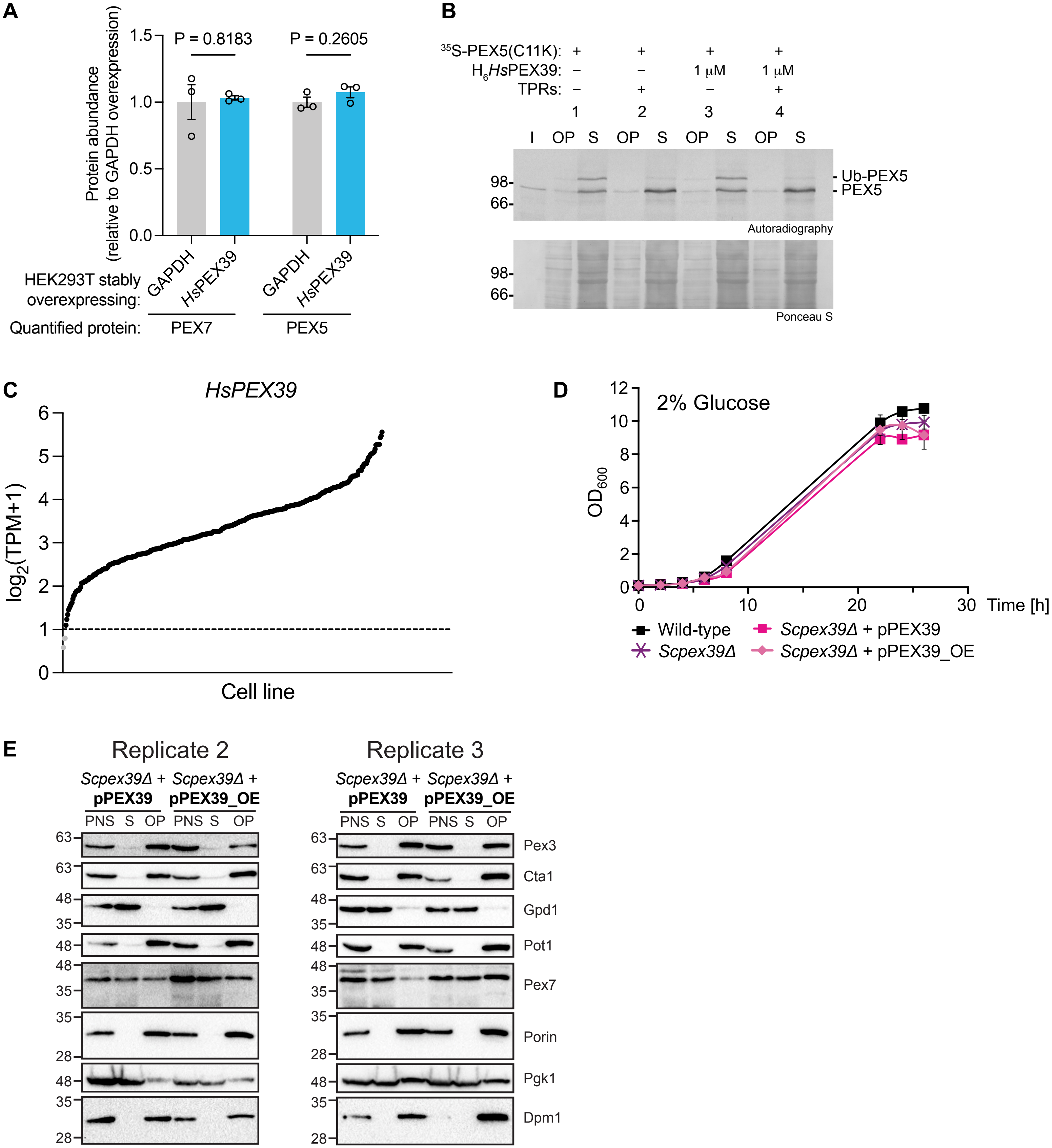

**FIGURE S4, related to FIGURE 4**

**A**

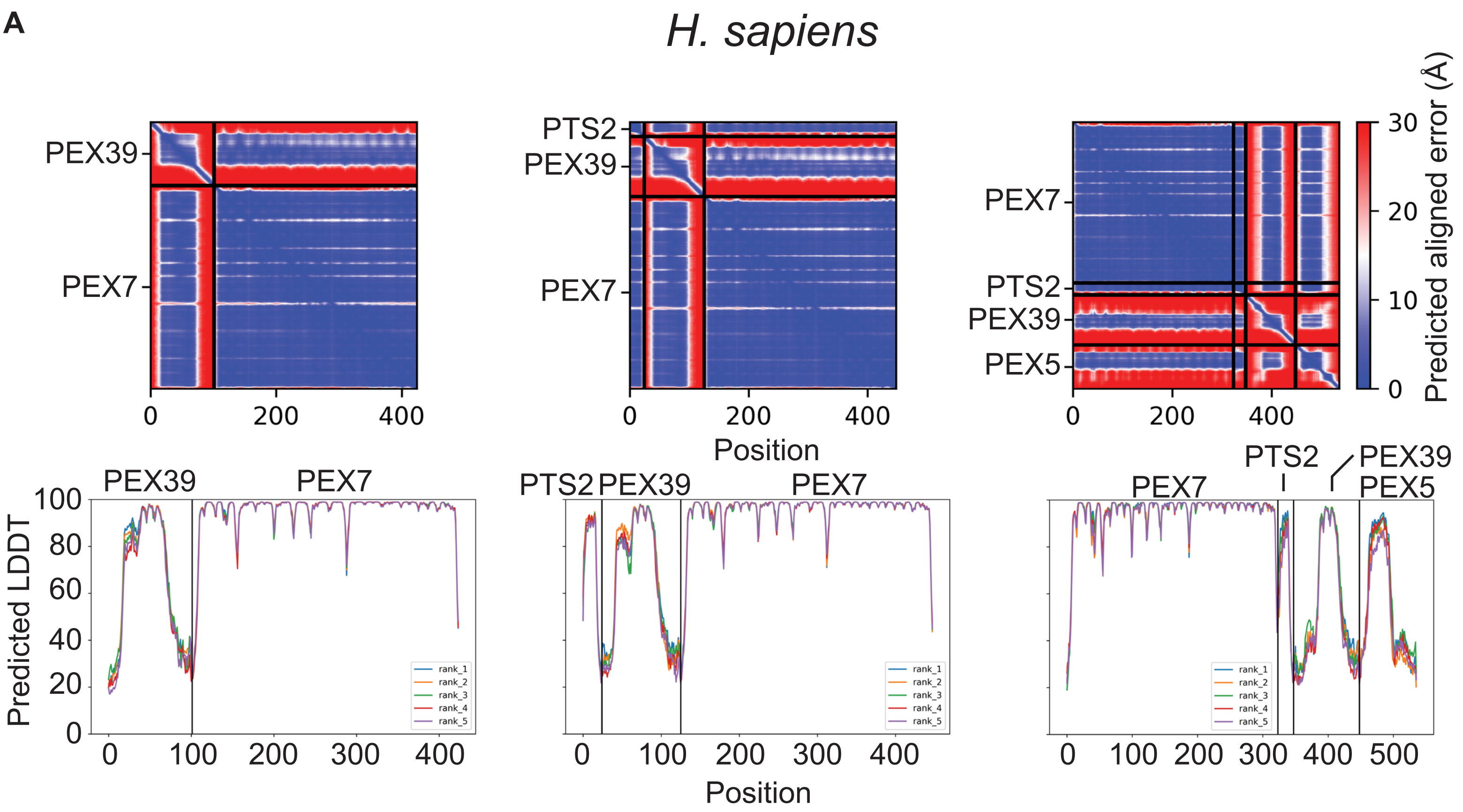


**B**

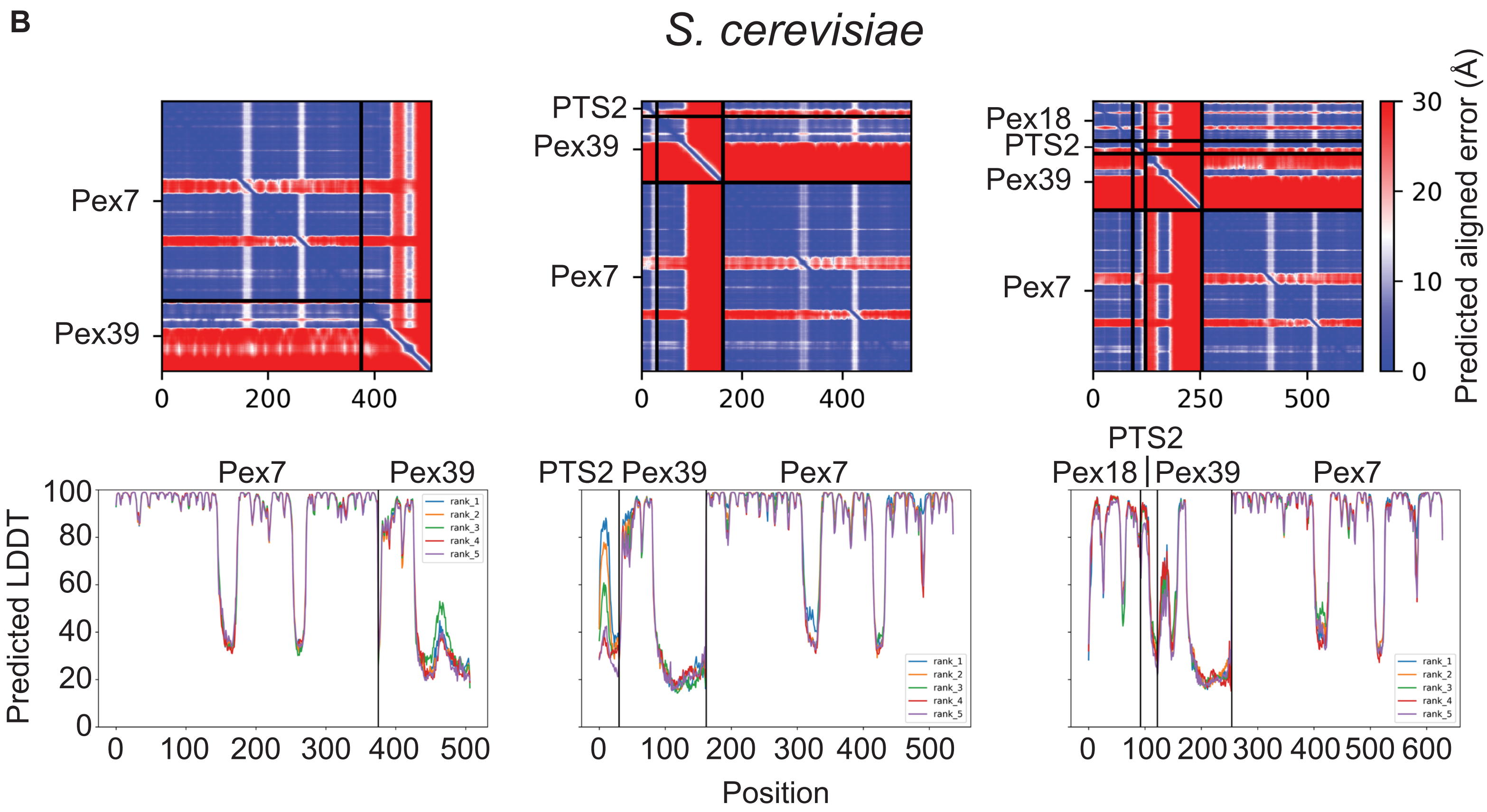

#### FIGURE S5, related to FIGURE 5

A

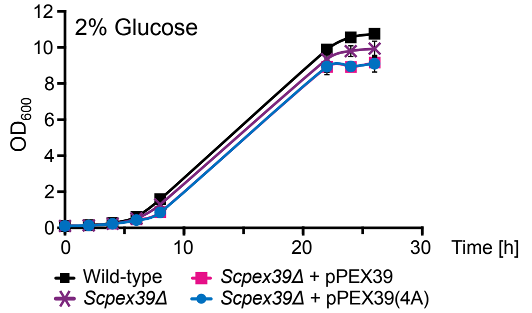


B

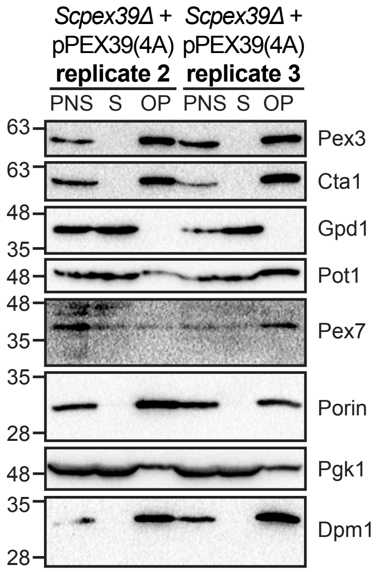


C

```
CLUSTAL O(1.2.4) multiple sequence alignment


tr|A0A6I8SMH6|A0A6I8SMH6_XENTR      MAVSGDGRLHAVRIPDRHGYAVEFSPYKPSTLACATSQNYGISGCGTLVVLEQSEGGIAV    60
sp|O00628|PEX7_HUMAN                MSAVCGGAARMLRTPGRHGYAAEFSPYLPGRLACATAQHYGIAGCGTLLILDPDEAGLRL    60
sp|Q9XF57|PEX7_ARATH                ---------MPVFKAPFNGYSVKFSPFYESRLAVATAQNFGILGNGRIHVLELAPGAPGV    51
sp|Q54WA3|PEX7_DICDI                ---------MKRFHSHFNGYSVEFSPFEEQRLACVTSQHFGIIGNGRQYILDVLDRDIGA    51
sp|P39108|PEX7_YEAST                --------MLRYHMQGFSGYGVQYSPFFDNRLAVAAGSNFGLVGNGKLFILEIDRSGRIV    52
sp|O59894|PEX7_PICPA                --------MFKFQTNGFSGYAVRYSPFYDNKIAVATSANYGLVGNGRLYVLSIMDDGNII    52
                                                      **...:**:    :* .:. ::*: * *   :*.

tr|A0A6I8SMH6|A0A6I8SMH6_XENTR      RRRLKKGHFFVSLKVLGCTEVPRIAESLQQEVSSFDWTDALFDVTWSEISENIVVTSSGD    120
sp|O00628|PEX7_HUMAN                -------------------------------FRSFDWNDGLFDVTWSENNEHVLITCSGD    89
sp|Q9XF57|PEX7_ARATH                -----------------------------TESVSYDTADAVYDVCWSESHDSVLIAAIGD    82
sp|Q54WA3|PEX7_DICDI                -------------------------------FKTFDTRDGLYDCTWSEENECHVASSSGD    80
sp|P39108|PEX7_YEAST                ------------------------------EVNSFLTQDCLFDLAWNESHENQVLVAQGD    82
sp|O59894|PEX7_PICPA                ------------------------------TDISYDTQDGLFGVAWSETNENHVLTSSGD    82
                                                                     ::   * ::.  *.*  :  :  . **

tr|A0A6I8SMH6|A0A6I8SMH6_XENTR      GSLQLWDITKPQ--GPLQVFKEHTQEVYSVDWSQTRGEQLIVSGSWDHTVKLWDPSFGKP    178
sp|O00628|PEX7_HUMAN                GSLQLWDTAKAA--GPLQVYKEHAQEVYSVDWSQTRGEQLVVSGSWDQTVKLWDPTVGKS    147
sp|Q9XF57|PEX7_ARATH                GSVKIYDTALPPPSNPIRSFQEHAREVQSVDYNPTRRDSF-LTSSWDDTVKLWAMDRPAS    141
sp|Q54WA3|PEX7_DICDI                GSIKIWDTQAPSGGRPIKSFEEHTKEVYSVDWNLVTKDTF-ITGSWDQSIKIWNPRMDRS    139
sp|P39108|PEX7_YEAST                GTLRLFDTTFKE--FPIAIFKEHEREVFSCNWNLVNRQNF-LSSSWDGSIKIWSPLRKQS    139
sp|O59894|PEX7_PICPA                GCVSLFDTTLKD--YPVMKFTEHQREVFSVDWSNIDKNLF-CSASWDGSVKVWSPGSNRN    139
                                    * : ::*        *:  : ** :** * ::.      : :  :.*** ::*:*

tr|A0A6I8SMH6|A0A6I8SMH6_XENTR      LCTF------------------------------------TGHENIIYSTIWSPHIPGC    201
sp|O00628|PEX7_HUMAN                LCTF------------------------------------RGHESIIYSTIWSPHIPGC    170
sp|Q9XF57|PEX7_ARATH                VRTF------------------------------------KEHAYCVYQAVWNPKHGDV    164
sp|Q54WA3|PEX7_DICDI                LKTF------------------------------------REHRYCIYSAIWSPRNAHL    162
sp|P39108|PEX7_YEAST                LMTLTPRPLEI-----TKMVDPLNA-----IILKKKSFTGISKNRNCVYQAQFSPHDQNL    189
sp|O59894|PEX7_PICPA                TSLLTLRSLASREEKTGRIEKPIPVVQPSQVPMSKTRPNIRNDNNDCVYDAKFSFHDPNI    199
                                       :                                       :   :*.: :. :

tr|A0A6I8SMH6|A0A6I8SMH6_XENTR      FASASGDQSLRIWDMKTPVSKV---VIPAHQAEILSCDWCKYDQNLLVTGAVDCSLKGWD    258
sp|O00628|PEX7_HUMAN                FASASGDQTLRIWDVKAAGVRI---VIPAHQAEILSCDWCKYNENLLVTGAVDCSLRGWD    227
sp|Q9XF57|PEX7_ARATH                FASASGDCTLRIWDVREPGSTM---IIPAHDFEILSCDWNKYDDCILATSSVDKTVKVWD    221
sp|Q54WA3|PEX7_DICDI                FASVSGDRTLKIWDSRDNRSLN---TIKAHDHEILTCDWNKYNDKEVVTGSVDKTIRIWD    219
sp|P39108|PEX7_YEAST                VLSCSGNSYASLFDIRLPSGKNQNNFLVHSGLEALTCDFNKYRPYVVATGGVDNAIRIWD    249
sp|O59894|PEX7_PICPA                IMSCNSDSHLQLWDTRLPNPL-FMDFVAHNGLEALSCDFNRYRPFVVASAGVDKLAKVWD    258
                                    . * ..:    ::* :          :     * *:**: :*    :.:..**   : **

tr|A0A6I8SMH6|A0A6I8SMH6_XENTR      LRTVRQPVF-------------------ELRGHNYAIRRVKFSPFHANIVASCSYDFTVR    299
sp|O00628|PEX7_HUMAN                LRNVRQPVF-------------------ELLGHTYAIRRVKFSPFHASVLASCSYDFTVR    268
sp|Q9XF57|PEX7_ARATH                VRSYRVPLA-------------------VLNGHGYAVRKVKFSPHRRSLIASCSYDMSVC    262
sp|Q54WA3|PEX7_DICDI                IRYPDRPTT-------------------ILRGHTYAVRRIKCSPHSESMLASCSYDMSVI    260
sp|P39108|PEX7_YEAST                IRMLNKNESATIKRTVPGQLHNSSCINEIPNAHGLAIRKVTWSPHHSNILMSASYDMTCR    309
sp|O59894|PEX7_PICPA                TRMIQPNVHSRPPRALN-----------KFMGHEFAIRKLAWSPHGPTQLLTCSYDMTVR    307
                                     *                             .*   *:*::  **.    . : :.***::

tr|A0A6I8SMH6|A0A6I8SMH6_XENTR      LWDFSKS-SS---------------LLETVNHHTEFVCGLDFSIL-TPGQIADCAWDETV    342
sp|O00628|PEX7_HUMAN                FWNFSKP-DS---------------LLETVEHHTEFTCGLDFSLQ-SPTQVADCSWDETI    311
sp|Q9XF57|PEX7_ARATH                LWDYMVE-DA---------------LVGRYDHHTEFAVGIDMSVL-VEGLMASTGWDELV    305
sp|Q54WA3|PEX7_DICDI                VWDRAREQDP---------------IIARMDHHTEFVVGLDWNMF-IDGQMASCSWDEQV    304
sp|P39108|PEX7_YEAST                IWRDLSNDGAKETYKTN-STDATKGSIFNFTQHSEFVFGADWSLWGKPGYVASTAWDGNL    368
sp|O59894|PEX7_PICPA                VWNDSPSPTSRVGLLDGASQPHAPPCSKIFSAHTEFVMGCDWSLWGEPGWVVTTGWDEMV    367
                                    .*                              *:**.  * *  .:       :.   .**    :

tr|A0A6I8SMH6|A0A6I8SMH6_XENTR      KIYFPSCLSLP-    353
sp|O00628|PEX7_HUMAN                KIYDPACLTIPA    323
sp|Q9XF57|PEX7_ARATH                YVWQQGMDPRAS    317
sp|Q54WA3|PEX7_DICDI                CVWNLGRPGQFR    316
sp|P39108|PEX7_YEAST                FVWNGLG-----    375
sp|O59894|PEX7_PICPA                YVWNTQRLQ---    376
                                     ::
```

D

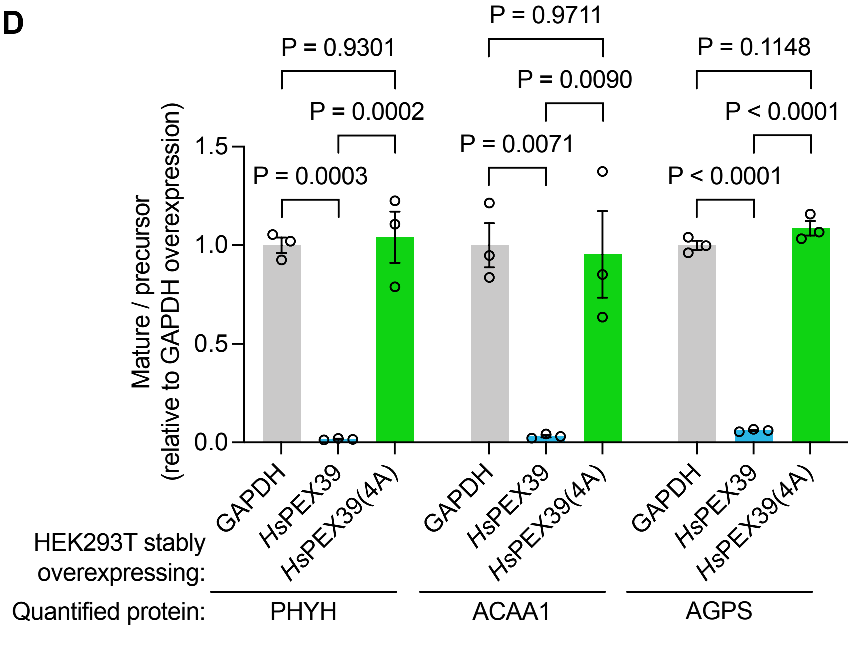


E

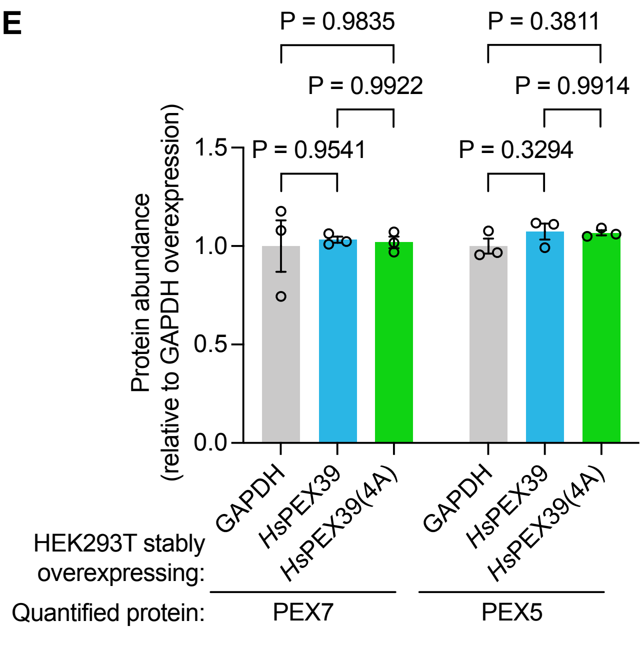

**FIGURE S6, related to FIGURE 6**

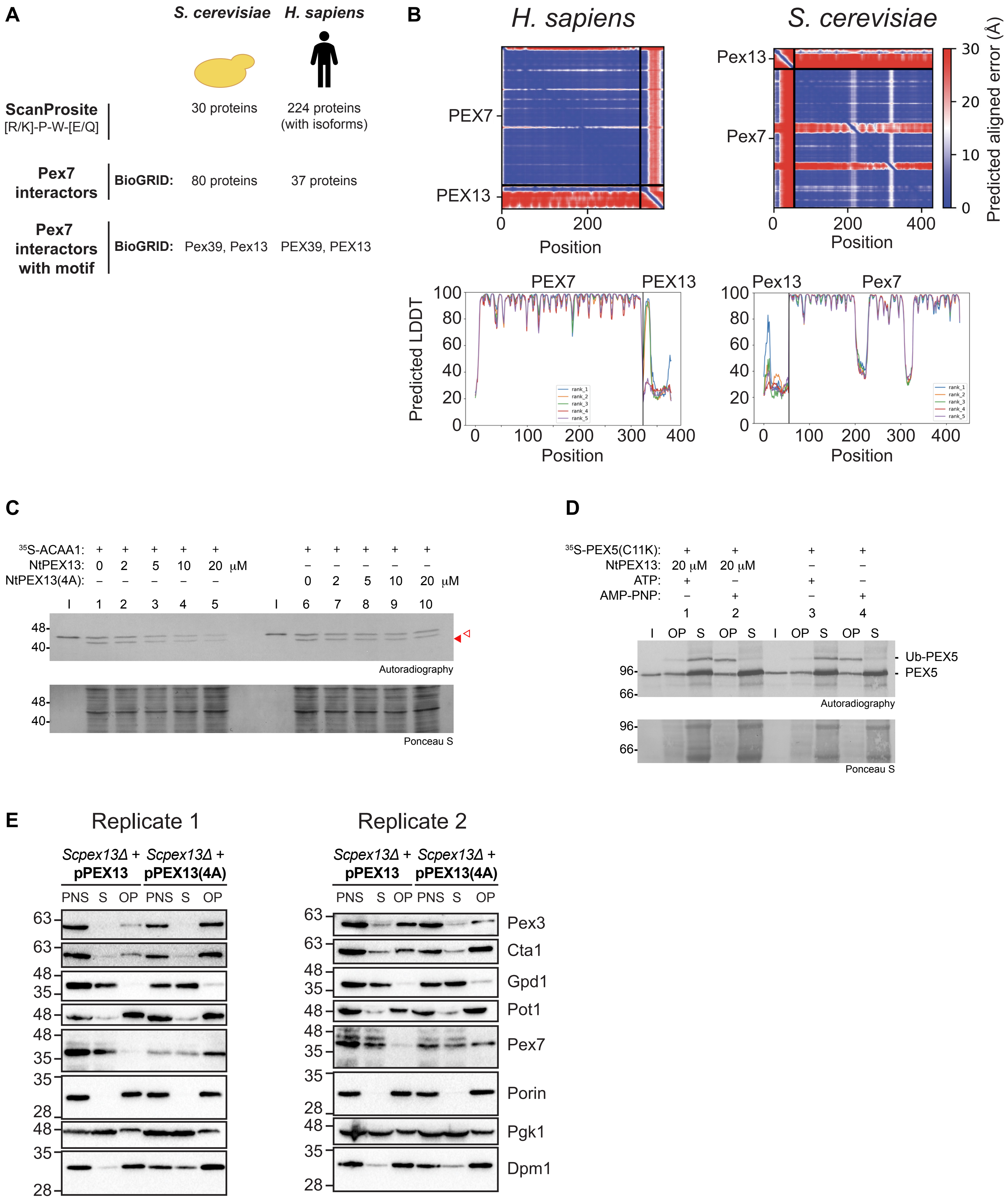

**FIGURE S7, related to FIGURE 7**

**A**

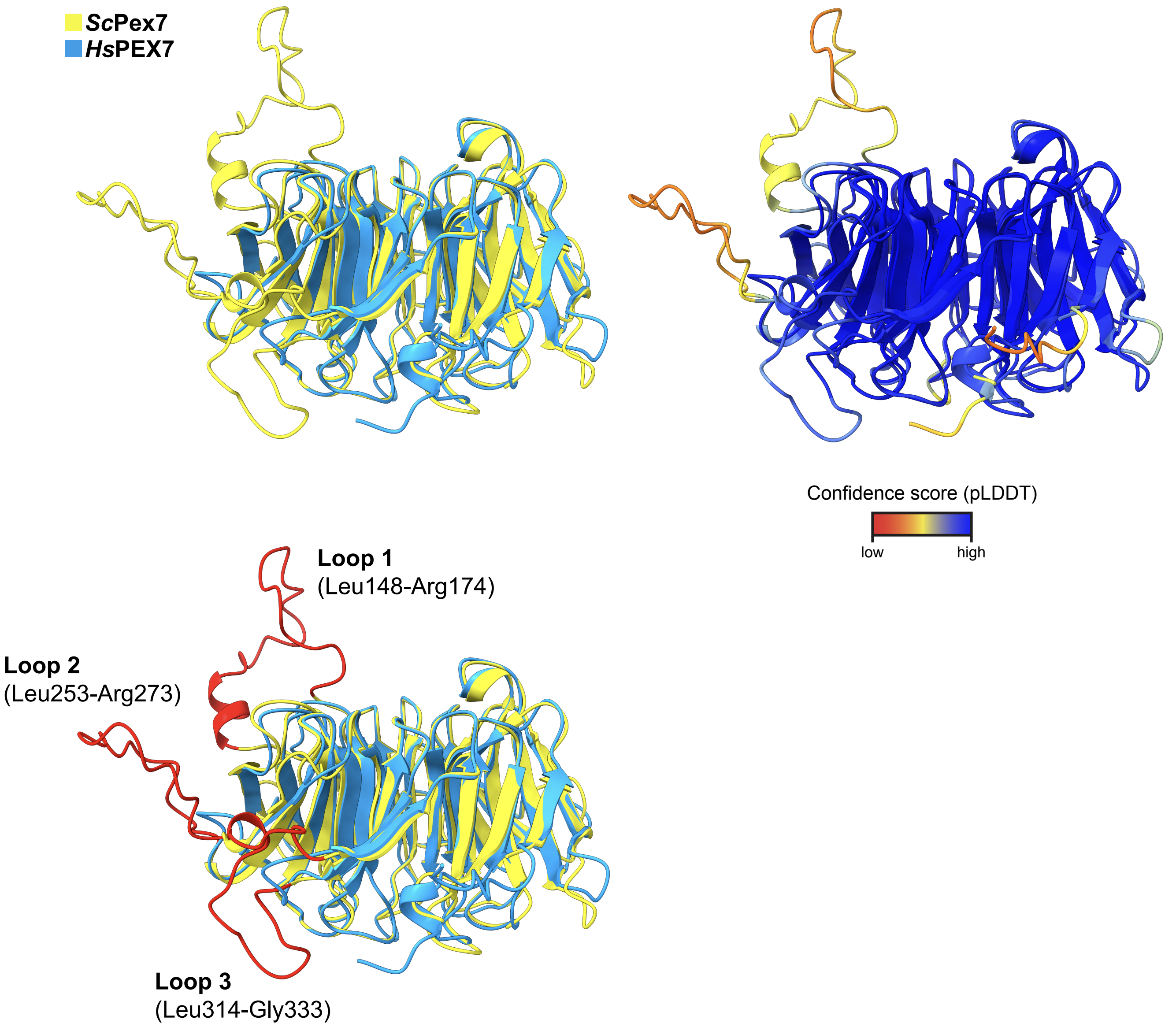
